## Supplementary Note 1 for "MS-DIAL 5 multimodal mass spectrometry data mining unveils lipidome complexities"

### Separation Workflow

Created by https://lipidomicstandards.org, version v2.3.2

#### Overall study design

| | | | |
|---|---|---|---|
| Title of the study | Overview of MS-DIAL lipid annotation for EAD-MS/MS | | |
| Document creation date | 02/05/2024 | Corresponding Email | |
| Principle investigator | Hiroshi Tsugawa | Is the workflow targeted or untargeted? | Untargeted |
| Institution | Tokyo University of Agriculture and Technology | Clinical | No |

#### Lipid extraction

| | | | |
|---|---|---|---|
| Extraction method | This reporting checklist is not for the description of sample analysis, but for the description of MS-DIAL 5 algorithm. | Were internal standards added prior extraction? | No |
| pH adjustment | None | | |

#### Analytical platform

| | | | |
|---|---|---|---|
| Which solvents were used | NA | Mass resolution for detected ion at MS1 | High resolution |
| Number of separation dimensions | One dimension | Resolution at m/z 200 at MS1 | 35000 |
| Separation type 1 | LC | Mass accuracy in ppm at MS1 | 2.5 |
| Separation mode 1 (liquid) | RP | Mass window for precursor ion isolation (in Da total isolation window) | 1 |
| Detector | Mass spectrometer | Mass resolution for detected ion at MS2 | High resolution |
| MS type | QTOF | Resolution at m/z 200 at MS2 | 25000 |
| MS vendor | SCIEX | Mass accuracy in ppm at MS2 | 5 |
| Ion source | ESI | Was/Were additional dimension/techniques used | Yes |
| MS Level | MS1, MS2 | | |

#### Quality control

| | | | |
|---|---|---|---|
| Blanks | No | Quality control | No |

#### Method qualification and validation

| | |
|---|---|
| Method validation | No |

#### Reporting

| | | | |
|---|---|---|---|
| Are reported raw data uploaded into repository? | Yes | Summary data | Identification data |
| Link to repository / ID to entry | http://prime.psc.riken.jp/menta.cgi/prime/drop_index, DM0054 | Raw data/upload | Yes |
| Are metadata available? | Yes | Additional comments | The uploaded files are msp files containing MS/MS spectra of lipid standards. |

### Lipid Class Descriptions

#### 1) BMP[M+NH4]+ / Lipid identification

| | | | |
|---|---|---|---|
| Lipid class | BMP | Did you presume assumptions for identification? | No |
| Derivatization | - | Check isomer overlap | No |
| MS Level for identification | MS1, MS2 | RT verified by standard | Yes |
| Identification level | Double bond position | Separation of isobaric/isomeric interferece confirmed | Yes |
| Polarity mode | Positive | Model for separation prediction | Yes |
| Type of positive (precursor)ion | [M+NH4]+ | Additional dimension/techniques | EAD |
| Fragments for identification | **Fragment name**<br>FA1(+C3H6O2)<br>FA2(+C3H6O2)<br>HG(GP,155)<br>-HG(GP,172)<br>FA1(+C3H5O4P)<br>FA2(+C3H5O4P) | How was/were the additional dimension(s) used? | To determine sn- positions |
| Isotope correction at MS1 | No | Was a model used to predict lipid molecule separation? | No |
| Isotope correction at MS2 | No | Lipid Identification Software | MS-DIAL |
| MS1 verified by standard | Yes | Data manipulation | Smoothing, Centroiding |
| MS2 verified by standard | Yes | Nomenclature for intact lipid molecule | Yes |
| Background check at MS1 | Yes | Nomenclature for fragment ions | No |
| Background check at MS2 | No | Further identification remarks | The reverse dot product similarity value, where the in silico spectrum is used for the library template, is used as the correlation coefficient value. The candidates are ranked by the reverse dot product score and the candidate with the highest similarity value is described as the representative C=C isomer candidate for the EAD-MS/MS spectrum. |

#### 1) BMP[M+NH4]+ / Lipid quantification

| | | | |
|---|---|---|---|
| Quantitative | No | Batch correction | No |
| Normalization to reference | No | Further quantification remarks | - |

#### 2) Acylcarnitine (CAR)[M+H]+ / Lipid identification

| | | | |
|---|---|---|---|
| Lipid class | Acylcarnitine (CAR) | Did you presume assumptions for identification? | No |
| Derivatization | - | Check isomer overlap | No |
| MS Level for identification | MS1, MS2 | RT verified by standard | Yes |
| Identification level | Double bond position | Separation of isobaric/isomeric interferece confirmed | Yes |
| Polarity mode | Positive | Model for separation prediction | Yes |
| Type of positive (precursor)ion | [M+H]+ | Additional dimension/techniques | EAD |
| Fragments for identification<br>**Fragment name**<br>Characteristic fragment (C4H5O2+) | | How was/were the additional dimension(s) used? | To determine doublebond positions |
| Isotope correction at MS1 | No | Was a model used to predict lipid molecule separation? | No |
| Isotope correction at MS2 | No | Lipid Identification Software | MS-DIAL |
| MS1 verified by standard | Yes | Data manipulation | Smoothing, Centroiding |
| MS2 verified by standard | Yes | Nomenclature for intact lipid molecule | Yes |
| Background check at MS1 | Yes | Nomenclature for fragment ions | No |
| Background check at MS2 | No | Further identification remarks | The reverse dot product similarity value, where the in silico spectrum is used for the library template, is used as the correlation coefficient value. The candidates are ranked by the reverse dot product score and the candidate with the highest similarity value is described as the representative C=C isomer candidate for the EAD-MS/MS spectrum. |

#### 2) Acylcarnitine (CAR)[M+H]+ / Lipid quantification

| | | | |
|---|---|---|---|
| Quantitative | No | Batch correction | No |
| Normalization to reference | No | Further quantification remarks | - |

#### 3) CL[M+NH4]+ / Lipid identification

| | | | |
|---|---|---|---|
| Lipid class | CL | Did you presume assumptions for identification? | No |
| Derivatization | - | Check isomer overlap | No |
| MS Level for identification | MS1, MS2 | RT verified by standard | Yes |
| Identification level | Molecular species level | Separation of isobaric/isomeric interferece confirmed | Yes |
| Polarity mode | Positive | Model for separation prediction | Yes |
| Type of positive (precursor)ion | [M+NH4]+ | Additional dimension/techniques | EAD |
| Fragments for identification | | How was/were the additional dimension(s) used? | To determine sn- positions |
| Isotope correction at MS1 | No | Was a model used to predict lipid molecule separation? | No |
| Isotope correction at MS2 | No | Lipid Identification Software | MS-DIAL |
| MS1 verified by standard | Yes | Data manipulation | Smoothing, Centroiding |
| MS2 verified by standard | Yes | Nomenclature for intact lipid molecule | Yes |
| Background check at MS1 | Yes | Nomenclature for fragment ions | No |
| Background check at MS2 | No | Further identification remarks | The reverse dot product similarity value, where the in silico spectrum is used for the library template, is used as the correlation coefficient value. The candidates are ranked by the reverse dot product score and the candidate with the highest similarity value is described as the representative C=C isomer candidate for the EAD-MS/MS spectrum. |

| Fragment name |
|---|
| Dehydro-monoacyl glycerols |
| Fatty acyl fragment |
| Dehydro-diacyl glycerols |

#### 3) CL[M+NH4]+ / Lipid quantification

| | | | |
|---|---|---|---|
| Quantitative | No | Batch correction | No |
| Normalization to reference | No | Further quantification remarks | - |

#### 4) DG[M+NH4]+ / Lipid identification

| | | | |
|---|---|---|---|
| Lipid class | DG | Did you presume assumptions for identification? | No |
| Derivatization | - | Check isomer overlap | No |
| MS Level for identification | MS1, MS2 | RT verified by standard | Yes |
| Identification level | Double bond position | Separation of isobaric/isomeric interferece confirmed | Yes |
| Polarity mode | Positive | Model for separation prediction | Yes |
| Type of positive (precursor)ion | [M+NH4]+ | Additional dimension/techniques | EAD |
| Fragments for identification | **Fragment name**<br>-(H2O+NH3,35)<br>The reverse dot product similarity value, where the in silico spectrum is used for the library template, is used as the correlation coefficient value. The candidates are ranked by the reverse dot product score and the candidate with the highest similarity value is described as the representative C=C isomer candidate for the EAD-MS/MS spectrum.<br>-FA1(-H)-(H2O+NH3)<br>-FA2(-H)-(H2O+NH3) | How was/were the additional dimension(s) used? | To determine doublebond positions |
| Isotope correction at MS1 | No | Was a model used to predict lipid molecule separation? | No |
| Isotope correction at MS2 | No | Lipid Identification Software | MS-DIAL |
| MS1 verified by standard | Yes | Data manipulation | Smoothing, Centroiding |
| MS2 verified by standard | Yes | Nomenclature for intact lipid molecule | Yes |
| Background check at MS1 | Yes | Nomenclature for fragment ions | No |
| Background check at MS2 | No | Further identification remarks | The reverse dot product similarity value, where the in silico spectrum is used for the library template, is used as the correlation coefficient value. The candidates are ranked by the reverse dot product score and the candidate with the highest similarity value is described as the representative C=C isomer candidate for the EAD-MS/MS spectrum. |

#### 4) DG[M+NH4]+ / Lipid quantification

| | | | |
|---|---|---|---|
| Quantitative | No | Batch correction | No |
| Normalization to reference | No | Further quantification remarks | - |

#### 5) DG[M+Na]+ / Lipid identification

| | | | |
|---|---|---|---|
| Lipid class | DG | Did you presume assumptions for identification? | No |
| Derivatization | - | Check isomer overlap | No |
| MS Level for identification | MS1, MS2 | RT verified by standard | Yes |
| Identification level | Double bond position | Separation of isobaric/isomeric interferece confirmed | Yes |
| Polarity mode | Positive | Model for separation prediction | Yes |
| Type of positive (precursor)ion | [M+Na]+ | Additional dimension/techniques | EAD |
| Fragments for identification | Fragment name: -FA1(-H)-(H2O+NH3); -FA2(-H)-(H2O+NH3) | How was/were the additional dimension(s) used? | To determine the sn- and doublebond positions |
| Isotope correction at MS1 | No | Was a model used to predict lipid molecule separation? | No |
| Isotope correction at MS2 | No | Lipid Identification Software | MS-DIAL |
| MS1 verified by standard | Yes | Data manipulation | Smoothing, Centroiding |
| MS2 verified by standard | Yes | Nomenclature for intact lipid molecule | Yes |
| Background check at MS1 | Yes | Nomenclature for fragment ions | No |
| Background check at MS2 | No | Further identification remarks | The reverse dot product similarity value, where the in silico spectrum is used for the library template, is used as the correlation coefficient value. The candidates are ranked by the reverse dot product score and the candidate with the highest similarity value is described as the representative C=C isomer candidate for the EAD-MS/MS spectrum. |

#### 5) DG[M+Na]+ / Lipid quantification

| | | | |
|---|---|---|---|
| Quantitative | No | Batch correction | No |
| Normalization to reference | No | Further quantification remarks | - |

#### 6) Diacylglyceryl hydroxymethyl-N,N,N-trimethyl-beta-alanine (DGTA)[M+H]+ / Lipid identification

| | | | |
|---|---|---|---|
| Lipid class | Diacylglyceryl hydroxymethyl-N,N,N-trimethyl-beta-alanine (DGTA) | Did you presume assumptions for identification? | No |
| Derivatization | - | Check isomer overlap | No |
| MS Level for identification | MS1, MS2 | RT verified by standard | Yes |
| Identification level | Double bond position | Separation of isobaric/isomeric interferece confirmed | Yes |
| Polarity mode | Positive | Model for separation prediction | Yes |
| Type of positive (precursor)ion | [M+H]+ | Additional dimension/techniques | EAD |
| Fragments for identification | (see fragment table below) | How was/were the additional dimension(s) used? | To determine sn- and doublebond positions |
| Isotope correction at MS1 | No | Was a model used to predict lipid molecule separation? | No |
| Isotope correction at MS2 | No | Lipid Identification Software | MS-DIAL |
| MS1 verified by standard | No | Data manipulation | Smoothing, Centroiding |
| MS2 verified by standard | No | Nomenclature for intact lipid molecule | Yes |
| Background check at MS1 | Yes | Nomenclature for fragment ions | No |
| Background check at MS2 | No | Further identification remarks | The reverse dot product similarity value, where the in silico spectrum is used for the library template, is used as the correlation coefficient value. The candidates are ranked by the reverse dot product score and the candidate with the highest similarity value is described as the representative C=C isomer candidate for the EAD-MS/MS spectrum. |

| Fragment name |
|---|
| Characteristic fragment (C7H14NO2+) |
| HG + C3H6+ |
| HG + C2H4O+ |

#### 6) Diacylglyceryl hydroxymethyl-N,N,N-trimethyl-beta-alanine (DGTA)[M+H]+ / Lipid quantification

| | | | |
|---|---|---|---|
| Quantitative | No | Batch correction | No |
| Normalization to reference | No | Further quantification remarks | - |

#### 7) Diacylglyceryl trimethylhomoserine (DGTS)[M+H]+ / Lipid identification

| | | | |
|---|---|---|---|
| Lipid class | Diacylglyceryl trimethylhomoserine (DGTS) | Did you presume assumptions for identification? | No |
| Derivatization | - | Check isomer overlap | No |
| MS Level for identification | MS1, MS2 | RT verified by standard | Yes |
| Identification level | Double bond position | Separation of isobaric/isomeric interferece confirmed | Yes |
| Polarity mode | Positive | Model for separation prediction | Yes |
| Type of positive (precursor)ion | [M+H]+ | Additional dimension/techniques | EAD |
| Fragments for identification | | How was/were the additional dimension(s) used? | To determine sn- and doublebond positions |

| Fragment name |
|---|
| Characteristic fragment (C6H12NO2+) |
| HG + C3H6+ |
| HG + C2H4O+ |

| | | | |
|---|---|---|---|
| Isotope correction at MS1 | No | Was a model used to predict lipid molecule separation? | No |
| Isotope correction at MS2 | No | Lipid Identification Software | MS-DIAL |
| MS1 verified by standard | No | Data manipulation | Smoothing, Centroiding |
| MS2 verified by standard | No | Nomenclature for intact lipid molecule | Yes |
| Background check at MS1 | Yes | Nomenclature for fragment ions | No |
| Background check at MS2 | No | Further identification remarks | The reverse dot product similarity value, where the in silico spectrum is used for the library template, is used as the correlation coefficient value. The candidates are ranked by the reverse dot product score and the candidate with the highest similarity value is described as the representative C=C isomer candidate for the EAD-MS/MS spectrum. |

#### 7) Diacylglyceryl trimethylhomoserine (DGTS)[M+H]+ / Lipid quantification

| | | | |
|---|---|---|---|
| Quantitative | No | Batch correction | No |
| Normalization to reference | No | Further quantification remarks | - |

#### 8) N, N-dimethylethylenediamine derivatized fatty acid (DMEDFA)[M+H]+ / Lipid identification

| | | | |
|---|---|---|---|
| Lipid class | N, N-dimethylethylenediamine derivatized fatty acid (DMEDFA) | Did you presume assumptions for identification? | No |
| Derivatization | - | Check isomer overlap | No |
| MS Level for identification | MS1, MS2 | RT verified by standard | Yes |
| Identification level | Double bond position | Separation of isobaric/isomeric interferece confirmed | Yes |
| Polarity mode | Positive | Model for separation prediction | Yes |
| Type of positive (precursor)ion | [M+H]+ | Additional dimension/techniques | EAD |
| Fragments for identification<br>**Fragment name**<br>NL of C2NH7 | | How was/were the additional dimension(s) used? | To determine doublebond positions |
| Isotope correction at MS1 | No | Was a model used to predict lipid molecule separation? | No |
| Isotope correction at MS2 | No | Lipid Identification Software | MS-DIAL |
| MS1 verified by standard | No | Data manipulation | Smoothing, Centroiding |
| MS2 verified by standard | No | Nomenclature for intact lipid molecule | Yes |
| Background check at MS1 | Yes | Nomenclature for fragment ions | No |
| Background check at MS2 | No | Further identification remarks | The reverse dot product similarity value, where the in silico spectrum is used for the library template, is used as the correlation coefficient value. The candidates are ranked by the reverse dot product score and the candidate with the highest similarity value is described as the representative C=C isomer candidate for the EAD-MS/MS spectrum. |

#### 8) N, N-dimethylethylenediamine derivatized fatty acid (DMEDFA)[M+H]+ / Lipid quantification

| | | | |
|---|---|---|---|
| Quantitative | No | Batch correction | No |
| Normalization to reference | No | Further quantification remarks | - |

#### 9) N, N-dimethylethylenediamine derivatized fatty acid ester of hydroxyl fatty acid (DMED-FAHFA)[M+H]+ / Lipid identification

| | | | |
|---|---|---|---|
| Lipid class | N, N-dimethylethylenediamine derivatized fatty acid ester of hydroxyl fatty acid (DMEDFAHFA) | Did you presume assumptions for identification? | No |
| Derivatization | - | Check isomer overlap | No |
| MS Level for identification | MS1, MS2 | RT verified by standard | Yes |
| Identification level | Double bond position | Separation of isobaric/isomeric interferece confirmed | Yes |
| Polarity mode | Positive | Model for separation prediction | Yes |
| Type of positive (precursor)ion | [M+H]+ | Additional dimension/techniques | EAD |
| Fragments for identification | **Fragment name**<br>The reverse dot product similarity value, where the in silico spectrum is used for the library template, is used as the correlation coefficient value. The candidates are ranked by the reverse dot product score and the candidate with the highest similarity value is described as the representative C=C isomer candidate for the EAD-MS/MS spectrum.<br>HFA -H2O +DMED<br>HFA -H2O +C2H5N<br>Specific fragments detached at OH position on HFA | How was/were the additional dimension(s) used? | To determine hydroxyl moiety position and doublebond positions |
| Isotope correction at MS1 | No | Was a model used to predict lipid molecule separation? | No |
| Isotope correction at MS2 | No | Lipid Identification Software | MS-DIAL |
| MS1 verified by standard | No | Data manipulation | Smoothing, Centroiding |
| MS2 verified by standard | No | Nomenclature for intact lipid molecule | Yes |
| Background check at MS1 | Yes | Nomenclature for fragment ions | No |
| Background check at MS2 | No | Further identification remarks | The reverse dot product similarity value, where the in silico spectrum is used for the library template, is used as the correlation coefficient value. The candidates are ranked by the reverse dot product score and the candidate with the highest similarity value is described as the representative C=C isomer candidate for the EAD-MS/MS spectrum. |

#### 9) N, N-dimethylethylenediamine derivatized fatty acid ester of hydroxyl fatty acid (DMED-FAHFA)[M+H]+ / Lipid quantification

| | | | |
|---|---|---|---|
| Quantitative | No | Batch correction | No |
| Normalization to reference | No | Further quantification remarks | - |

#### 10) PC O[M+H]+ / Lipid identification

| | | | |
|---|---|---|---|
| Lipid class | PC O | Did you presume assumptions for identification? | No |
| Derivatization | - | Check isomer overlap | No |
| MS Level for identification | MS1, MS2 | RT verified by standard | Yes |
| Identification level | Double bond position | Separation of isobaric/isomeric interferece confirmed | Yes |
| Polarity mode | Positive | Model for separation prediction | Yes |
| Type of positive (precursor)ion | [M+H]+ | Additional dimension/techniques | EAD |
| Fragments for identification | | How was/were the additional dimension(s) used? | To determine sn- and doublebond positions |

| Fragment name |
|---|
| HG(PC,184) |
| NL of FA |
| NL of Ether |
| HG + C3H5 |
| HG + C2H3O |

| | | | |
|---|---|---|---|
| Isotope correction at MS1 | No | Was a model used to predict lipid molecule separation? | No |
| Isotope correction at MS2 | No | Lipid Identification Software | MS-DIAL |
| MS1 verified by standard | Yes | Data manipulation | Smoothing, Centroiding |
| MS2 verified by standard | Yes | Nomenclature for intact lipid molecule | Yes |
| Background check at MS1 | Yes | Nomenclature for fragment ions | No |
| Background check at MS2 | No | Further identification remarks | The reverse dot product similarity value, where the in silico spectrum is used for the library template, is used as the correlation coefficient value. The candidates are ranked by the reverse dot product score and the candidate with the highest similarity value is described as the representative C=C isomer candidate for the EAD-MS/MS spectrum. |

#### 10) PC O[M+H]+ / Lipid quantification

| | | | |
|---|---|---|---|
| Quantitative | No | Batch correction | No |
| Normalization to reference | No | Further quantification remarks | - |

#### 11) PC P[M+H]+ / Lipid identification

| | | | |
|---|---|---|---|
| Lipid class | PC P | Did you presume assumptions for identification? | No |
| Derivatization | - | Check isomer overlap | No |
| MS Level for identification | MS1, MS2 | RT verified by standard | Yes |
| Identification level | Double bond position | Separation of isobaric/isomeric interferece confirmed | Yes |
| Polarity mode | Positive | Model for separation prediction | Yes |
| Type of positive (precursor)ion | [M+H]+ | Additional dimension/techniques | EAD |
| Fragments for identification | | How was/were the additional dimension(s) used? | To determine sn- and doublebond positions |

| Fragment name |
|---|
| HG(PC,184) |
| NL of FA |
| NL of Ether |
| HG + C3H5 |
| HG + C2H3O |

| | | | |
|---|---|---|---|
| Isotope correction at MS1 | No | Was a model used to predict lipid molecule separation? | No |
| Isotope correction at MS2 | No | Lipid Identification Software | MS-DIAL |
| MS1 verified by standard | Yes | Data manipulation | Smoothing, Centroiding |
| MS2 verified by standard | Yes | Nomenclature for intact lipid molecule | Yes |
| Background check at MS1 | Yes | Nomenclature for fragment ions | No |
| Background check at MS2 | No | Further identification remarks | The reverse dot product similarity value, where the in silico spectrum is used for the library template, is used as the correlation coefficient value. The candidates are ranked by the reverse dot product score and the candidate with the highest similarity value is described as the representative C=C isomer candidate for the EAD-MS/MS spectrum. |

#### 11) PC P[M+H]+ / Lipid quantification

| | | | |
|---|---|---|---|
| Quantitative | No | Batch correction | No |
| Normalization to reference | No | Further quantification remarks | - |

#### 12) PE O[M+H]+ / Lipid identification

| | | | |
|---|---|---|---|
| Lipid class | PE O | Did you presume assumptions for identification? | No |
| Derivatization | - | Check isomer overlap | No |
| MS Level for identification | MS1, MS2 | RT verified by standard | Yes |
| Identification level | Double bond position | Separation of isobaric/isomeric interferece confirmed | Yes |
| Polarity mode | Positive | Model for separation prediction | Yes |
| Type of positive (precursor)ion | [M+H]+ | Additional dimension/techniques | EAD |
| Fragments for identification | | How was/were the additional dimension(s) used? | To determine doublebond positions |

| Fragment name |
|---|
| -HG(PE,141) |
| NL of FA |
| NL of Ether |
| HG + C3H6+ |
| HG + C2H4O+ |

| | | | |
|---|---|---|---|
| Isotope correction at MS1 | No | Was a model used to predict lipid molecule separation? | No |
| Isotope correction at MS2 | No | Lipid Identification Software | MS-DIAL |
| MS1 verified by standard | Yes | Data manipulation | Smoothing, Centroiding |
| MS2 verified by standard | Yes | Nomenclature for intact lipid molecule | Yes |
| Background check at MS1 | Yes | Nomenclature for fragment ions | No |
| Background check at MS2 | No | Further identification remarks | The reverse dot product similarity value, where the in silico spectrum is used for the library template, is used as the correlation coefficient value. The candidates are ranked by the reverse dot product score and the candidate with the highest similarity value is described as the representative C=C isomer candidate for the EAD-MS/MS spectrum. |

#### 12) PE O[M+H]+ / Lipid quantification

| | | | |
|---|---|---|---|
| Quantitative | No | Batch correction | No |
| Normalization to reference | No | Further quantification remarks | - |

#### 13) PE P[M+H]+ / Lipid identification

| | | | |
|---|---|---|---|
| Lipid class | PE P | Did you presume assumptions for identification? | No |
| Derivatization | - | Check isomer overlap | No |
| MS Level for identification | MS1, MS2 | RT verified by standard | Yes |
| Identification level | Double bond position | Separation of isobaric/isomeric interferece confirmed | Yes |
| Polarity mode | Positive | Model for separation prediction | Yes |
| Type of positive (precursor)ion | [M+H]+ | Additional dimension/techniques | EAD |
| Fragments for identification | | How was/were the additional dimension(s) used? | To determine doublebond positions |

| Fragment name |
|---|
| -HG(PE,141) |
| -FA1(-H) |
| -FA2+(C3H5O2) |
| HG+ |
| HG + C3H6+ |
| HG + C2H4O+ |

| | | | |
|---|---|---|---|
| Isotope correction at MS1 | No | Was a model used to predict lipid molecule separation? | No |
| Isotope correction at MS2 | No | Lipid Identification Software | MS-DIAL |
| MS1 verified by standard | Yes | Data manipulation | Smoothing, Centroiding |
| MS2 verified by standard | Yes | Nomenclature for intact lipid molecule | Yes |
| Background check at MS1 | Yes | Nomenclature for fragment ions | No |
| Background check at MS2 | No | Further identification remarks | The reverse dot product similarity value, where the in silico spectrum is used for the library template, is used as the correlation coefficient value. The candidates are ranked by the reverse dot product score and the candidate with the highest similarity value is described as the representative C=C isomer candidate for the EAD-MS/MS spectrum. |

#### 13) PE P[M+H]+ / Lipid quantification

| | | | |
|---|---|---|---|
| Quantitative | No | Batch correction | No |
| Normalization to reference | No | Further quantification remarks | - |

#### 14) Hemibismonoacylglycerophosphate (HBMP)[M+NH4]+ / Lipid identification

| | | | |
|---|---|---|---|
| Lipid class | Hemibismonoacylglycerophosphate (HBMP) | Did you presume assumptions for identification? | No |
| Derivatization | - | Check isomer overlap | No |
| MS Level for identification | MS1, MS2 | RT verified by standard | Yes |
| Identification level | Molecular species level | Separation of isobaric/isomeric interferece confirmed | Yes |
| Polarity mode | Positive | Model for separation prediction | Yes |
| Type of positive (precursor)ion | [M+NH4]+ | Additional dimension/techniques | EAD |
| Fragments for identification | | How was/were the additional dimension(s) used? | To determine sn- positions |

| Fragment name |
|---|
| Dehydro-monoacyl glycerols |
| Dehydro-diacyl glycerol |
| NL of FA |

| | | | |
|---|---|---|---|
| Isotope correction at MS1 | No | Was a model used to predict lipid molecule separation? | No |
| Isotope correction at MS2 | No | Lipid Identification Software | MS-DIAL |
| MS1 verified by standard | No | Data manipulation | Smoothing, Centroiding |
| MS2 verified by standard | No | Nomenclature for intact lipid molecule | Yes |
| Background check at MS1 | Yes | Nomenclature for fragment ions | No |
| Background check at MS2 | No | Further identification remarks | The reverse dot product similarity value, where the in silico spectrum is used for the library template, is used as the correlation coefficient value. The candidates are ranked by the reverse dot product score and the candidate with the highest similarity value is described as the representative C=C isomer candidate for the EAD-MS/MS spectrum. |

#### 14) Hemibismonoacylglycerophosphate (HBMP)[M+NH4]+ / Lipid quantification

| | | | |
|---|---|---|---|
| Quantitative | No | Batch correction | No |
| Normalization to reference | No | Further quantification remarks | - |

#### 15) Lysodiacylglyceryl hydroxymethyl-N,N,N-trimethyl-beta-alanine (LDGTA)[M+H]+ / Lipid identification

| | | | |
|---|---|---|---|
| Lipid class | Lysodiacylglyceryl hydroxymethyl-N,N,N-trimethyl-beta-alanine (LDGTA) | Did you presume assumptions for identification? | No |
| Derivatization | - | Check isomer overlap | No |
| MS Level for identification | MS1, MS2 | RT verified by standard | Yes |
| Identification level | Double bond position | Separation of isobaric/isomeric interferece confirmed | Yes |
| Polarity mode | Positive | Model for separation prediction | Yes |
| Type of positive (precursor)ion | [M+H]+ | Additional dimension/techniques | EAD |
| Fragments for identification | **Fragment name**<br>Characteristic fragment (C7H14NO2+)<br>HG + C2H4O+ | How was/were the additional dimension(s) used? | To determine doublebond positions |
| Isotope correction at MS1 | No | Was a model used to predict lipid molecule separation? | No |
| Isotope correction at MS2 | No | Lipid Identification Software | MS-DIAL |
| MS1 verified by standard | No | Data manipulation | Smoothing, Centroiding |
| MS2 verified by standard | No | Nomenclature for intact lipid molecule | Yes |
| Background check at MS1 | Yes | Nomenclature for fragment ions | No |
| Background check at MS2 | No | Further identification remarks | The reverse dot product similarity value, where the in silico spectrum is used for the library template, is used as the correlation coefficient value. The candidates are ranked by the reverse dot product score and the candidate with the highest similarity value is described as the representative C=C isomer candidate for the EAD-MS/MS spectrum. |

#### 15) Lysodiacylglyceryl hydroxymethyl-N,N,N-trimethyl-beta-alanine (LDGTA)[M+H]+ / Lipid quantification

| | | | |
|---|---|---|---|
| Quantitative | No | Batch correction | No |
| Normalization to reference | No | Further quantification remarks | - |

#### 16) Lysoiacylglyceryl trimethylhomoserine (LDGTS)[M+H]+ / Lipid identification

| | | | |
|---|---|---|---|
| Lipid class | Lysoiacylglyceryl trimethylhomoserine (LDGTS) | Did you presume assumptions for identification? | No |
| Derivatization | - | Check isomer overlap | No |
| MS Level for identification | MS1, MS2 | RT verified by standard | Yes |
| Identification level | Double bond position | Separation of isobaric/isomeric interferece confirmed | Yes |
| Polarity mode | Positive | Model for separation prediction | Yes |
| Type of positive (precursor)ion | [M+H]+ | Additional dimension/techniques | EAD |
| Fragments for identification | | How was/were the additional dimension(s) used? | To determine doublebond positions |

| Fragment name |
|---|
| Characteristic fragment (C6H12NO2+) |
| HG + C2H4O+ |

| | | | |
|---|---|---|---|
| Isotope correction at MS1 | No | Was a model used to predict lipid molecule separation? | No |
| Isotope correction at MS2 | No | Lipid Identification Software | MS-DIAL |
| MS1 verified by standard | No | Data manipulation | Smoothing, Centroiding |
| MS2 verified by standard | No | Nomenclature for intact lipid molecule | Yes |
| Background check at MS1 | Yes | Nomenclature for fragment ions | No |
| Background check at MS2 | No | Further identification remarks | The reverse dot product similarity value, where the in silico spectrum is used for the library template, is used as the correlation coefficient value. The candidates are ranked by the reverse dot product score and the candidate with the highest similarity value is described as the representative C=C isomer candidate for the EAD-MS/MS spectrum. |

#### 16) Lysoiacylglyceryl trimethylhomoserine (LDGTS)[M+H]+ / Lipid quantification

| | | | |
|---|---|---|---|
| Quantitative | No | Batch correction | No |
| Normalization to reference | No | Further quantification remarks | - |

#### 17) LPC[M+H]+ / Lipid identification

| | | | |
|---|---|---|---|
| Lipid class | LPC | Did you presume assumptions for identification? | No |
| Derivatization | - | Check isomer overlap | No |
| MS Level for identification | MS1, MS2 | RT verified by standard | Yes |
| Identification level | Double bond position | Separation of isobaric/isomeric interferece confirmed | Yes |
| Polarity mode | Positive | Model for separation prediction | Yes |
| Type of positive (precursor)ion | [M+H]+ | Additional dimension/techniques | EAD |
| Fragments for identification | | How was/were the additional dimension(s) used? | To determine sn- and doublebond positions |

| Fragment name |
|---|
| HG(PC,184) |
| (C5H13NO,104) |
| HG + C3H5 |
| HG + C2H3O |
| NL of FA |

| | | | |
|---|---|---|---|
| Isotope correction at MS1 | No | Was a model used to predict lipid molecule separation? | No |
| Isotope correction at MS2 | No | Lipid Identification Software | MS-DIAL |
| MS1 verified by standard | Yes | Data manipulation | Smoothing, Centroiding |
| MS2 verified by standard | Yes | Nomenclature for intact lipid molecule | Yes |
| Background check at MS1 | Yes | Nomenclature for fragment ions | No |
| Background check at MS2 | No | Further identification remarks | The reverse dot product similarity value, where the in silico spectrum is used for the library template, is used as the correlation coefficient value. The candidates are ranked by the reverse dot product score and the candidate with the highest similarity value is described as the representative C=C isomer candidate for the EAD-MS/MS spectrum. |

#### 17) LPC[M+H]+ / Lipid quantification

| | | | |
|---|---|---|---|
| Quantitative | No | Batch correction | No |
| Normalization to reference | No | Further quantification remarks | - |

#### 18) LPE[M+H]+ / Lipid identification

| | | | |
|---|---|---|---|
| Lipid class | LPE | Did you presume assumptions for identification? | No |
| Derivatization | - | Check isomer overlap | No |
| MS Level for identification | MS1, MS2 | RT verified by standard | Yes |
| Identification level | Double bond position | Separation of isobaric/isomeric interferece confirmed | Yes |
| Polarity mode | Positive | Model for separation prediction | Yes |
| Type of positive (precursor)ion | [M+H]+ | Additional dimension/techniques | EAD |
| Fragments for identification | **Fragment name**<br>-HG(PE,141)<br>HG + C3H6+<br>HG + C2H4O+<br>HG+<br>NL of FA | How was/were the additional dimension(s) used? | To determine sn- and doublebond positions |
| Isotope correction at MS1 | No | Was a model used to predict lipid molecule separation? | No |
| Isotope correction at MS2 | No | Lipid Identification Software | MS-DIAL |
| MS1 verified by standard | Yes | Data manipulation | Smoothing, Centroiding |
| MS2 verified by standard | Yes | Nomenclature for intact lipid molecule | Yes |
| Background check at MS1 | Yes | Nomenclature for fragment ions | No |
| Background check at MS2 | No | Further identification remarks | The reverse dot product similarity value, where the in silico spectrum is used for the library template, is used as the correlation coefficient value. The candidates are ranked by the reverse dot product score and the candidate with the highest similarity value is described as the representative C=C isomer candidate for the EAD-MS/MS spectrum. |

#### 18) LPE[M+H]+ / Lipid quantification

| | | | |
|---|---|---|---|
| Quantitative | No | Batch correction | No |
| Normalization to reference | No | Further quantification remarks | - |

#### 19) LPG[M+H]+ / Lipid identification

| | | | |
|---|---|---|---|
| Lipid class | LPG | Did you presume assumptions for identification? | No |
| Derivatization | - | Check isomer overlap | No |
| MS Level for identification | MS1, MS2 | RT verified by standard | Yes |
| Identification level | sn Position | Separation of isobaric/isomeric interferece confirmed | Yes |
| Polarity mode | Positive | Model for separation prediction | Yes |
| Type of positive (precursor)ion | [M+H]+ | Additional dimension/techniques | EAD |
| Fragments for identification | | How was/were the additional dimension(s) used? | To determine sn- position |

| Fragment name |
|---|
| -HG(PG,172) |
| HG + C3H6+ |
| HG + C2H4O+ |
| HG+ |
| HG+ -H2O |
| NL of FA |

| | | | |
|---|---|---|---|
| Isotope correction at MS1 | No | Was a model used to predict lipid molecule separation? | No |
| Isotope correction at MS2 | No | Lipid Identification Software | MS-DIAL |
| MS1 verified by standard | Yes | Data manipulation | Smoothing, Centroiding |
| MS2 verified by standard | Yes | Nomenclature for intact lipid molecule | Yes |
| Background check at MS1 | Yes | Nomenclature for fragment ions | No |
| Background check at MS2 | No | Further identification remarks | The reverse dot product similarity value, where the in silico spectrum is used for the library template, is used as the correlation coefficient value. The candidates are ranked by the reverse dot product score and the candidate with the highest similarity value is described as the representative C=C isomer candidate for the EAD-MS/MS spectrum. |

#### 19) LPG[M+H]+ / Lipid quantification

| | | | |
|---|---|---|---|
| Quantitative | No | Batch correction | No |
| Normalization to reference | No | Further quantification remarks | - |

#### 20) LPI[M+NH4]+ / Lipid identification

| | | | |
|---|---|---|---|
| Lipid class | LPI | Did you presume assumptions for identification? | No |
| Derivatization | - | Check isomer overlap | No |
| MS Level for identification | MS1, MS2 | RT verified by standard | Yes |
| Identification level | sn Position | Separation of isobaric/isomeric interferece confirmed | Yes |
| Polarity mode | Positive | Model for separation prediction | Yes |
| Type of positive (precursor)ion | [M+NH4]+ | Additional dimension/techniques | EAD |
| Fragments for identification | | How was/were the additional dimension(s) used? | To determine sn- position |

| Fragment name |
|---|
| -(H2O,18) |
| GP(155) + H2O |
| -HG(PI,260) |
| NL of FA |
| HG + C3H6+ |
| HG + C2H4O+ |

| | | | |
|---|---|---|---|
| Isotope correction at MS1 | No | Was a model used to predict lipid molecule separation? | No |
| Isotope correction at MS2 | No | Lipid Identification Software | MS-DIAL |
| MS1 verified by standard | Yes | Data manipulation | Smoothing, Centroiding |
| MS2 verified by standard | Yes | Nomenclature for intact lipid molecule | Yes |
| Background check at MS1 | Yes | Nomenclature for fragment ions | No |
| Background check at MS2 | No | Further identification remarks | The reverse dot product similarity value, where the in silico spectrum is used for the library template, is used as the correlation coefficient value. The candidates are ranked by the reverse dot product score and the candidate with the highest similarity value is described as the representative C=C isomer candidate for the EAD-MS/MS spectrum. |

#### 20) LPI[M+NH4]+ / Lipid quantification

| | | | |
|---|---|---|---|
| Quantitative | No | Batch correction | No |
| Normalization to reference | No | Further quantification remarks | - |

#### 21) LPS[M+H]+ / Lipid identification

| | | | |
|---|---|---|---|
| Lipid class | LPS | Did you presume assumptions for identification? | No |
| Derivatization | - | Check isomer overlap | No |
| MS Level for identification | MS1, MS2 | RT verified by standard | Yes |
| Identification level | Double bond position | Separation of isobaric/isomeric interferece confirmed | Yes |
| Polarity mode | Positive | Model for separation prediction | Yes |
| Type of positive (precursor)ion | [M+H]+ | Additional dimension/techniques | EAD |
| Fragments for identification | Fragment name: -(C3H7NO3,105); HG+; HG + C3H6+; HG + C2H4O+; NL of FA | How was/were the additional dimension(s) used? | To determine sn- and doublebond positions |
| Isotope correction at MS1 | No | Was a model used to predict lipid molecule separation? | No |
| Isotope correction at MS2 | No | Lipid Identification Software | MS-DIAL |
| MS1 verified by standard | Yes | Data manipulation | Smoothing, Centroiding |
| MS2 verified by standard | Yes | Nomenclature for intact lipid molecule | Yes |
| Background check at MS1 | Yes | Nomenclature for fragment ions | No |
| Background check at MS2 | No | Further identification remarks | The reverse dot product similarity value, where the in silico spectrum is used for the library template, is used as the correlation coefficient value. The candidates are ranked by the reverse dot product score and the candidate with the highest similarity value is described as the representative C=C isomer candidate for the EAD-MS/MS spectrum. |

#### 21) LPS[M+H]+ / Lipid quantification

| | | | |
|---|---|---|---|
| Quantitative | No | Batch correction | No |
| Normalization to reference | No | Further quantification remarks | - |

#### 22) PC[M+H]+ / Lipid identification

| | | | |
|---|---|---|---|
| Lipid class | PC | Did you presume assumptions for identification? | No |
| Derivatization | - | Check isomer overlap | No |
| MS Level for identification | MS1, MS2 | RT verified by standard | Yes |
| Identification level | Double bond position | Separation of isobaric/isomeric interferece confirmed | Yes |
| Polarity mode | Positive | Model for separation prediction | Yes |
| Type of positive (precursor)ion | [M+H]+ | Additional dimension/techniques | EAD |
| Fragments for identification | | How was/were the additional dimension(s) used? | To determine sn- and doublebond positions |

| Fragment name |
|---|
| HG(PC,184) |
| HG + C3H5 |
| HG + C2H3O |
| NL of HG |
| NL of FA |

| | | | |
|---|---|---|---|
| Isotope correction at MS1 | No | Was a model used to predict lipid molecule separation? | No |
| Isotope correction at MS2 | No | Lipid Identification Software | MS-DIAL |
| MS1 verified by standard | Yes | Data manipulation | Smoothing, Centroiding |
| MS2 verified by standard | Yes | Nomenclature for intact lipid molecule | Yes |
| Background check at MS1 | Yes | Nomenclature for fragment ions | No |
| Background check at MS2 | No | Further identification remarks | The reverse dot product similarity value, where the in silico spectrum is used for the library template, is used as the correlation coefficient value. The candidates are ranked by the reverse dot product score and the candidate with the highest similarity value is described as the representative C=C isomer candidate for the EAD-MS/MS spectrum. |

#### 22) PC[M+H]+ / Lipid quantification

| | | | |
|---|---|---|---|
| Quantitative | No | Batch correction | No |
| Normalization to reference | No | Further quantification remarks | - |

#### 23) PC[M+Na]+ / Lipid identification

| | | | |
|---|---|---|---|
| Lipid class | PC | Did you presume assumptions for identification? | No |
| Derivatization | - | Check isomer overlap | No |
| MS Level for identification | MS1, MS2 | RT verified by standard | Yes |
| Identification level | Double bond position | Separation of isobaric/isomeric interferece confirmed | Yes |
| Polarity mode | Positive | Model for separation prediction | Yes |
| Type of positive (precursor)ion | [M+Na]+ | Additional dimension/techniques | EAD |
| Fragments for identification | | How was/were the additional dimension(s) used? | To determine sn- and doublebond positions |

| Fragment name |
|---|
| HG(PC,184) + Na+ |
| HG + C3H5Na |
| HG + C2H3ONa |
| NL of HG |
| NL of FA |
| NL of C3H9N |

| | | | |
|---|---|---|---|
| Isotope correction at MS1 | No | Was a model used to predict lipid molecule separation? | No |
| Isotope correction at MS2 | No | Lipid Identification Software | MS-DIAL |
| MS1 verified by standard | Yes | Data manipulation | Smoothing, Centroiding |
| MS2 verified by standard | Yes | Nomenclature for intact lipid molecule | Yes |
| Background check at MS1 | Yes | Nomenclature for fragment ions | No |
| Background check at MS2 | No | Further identification remarks | The reverse dot product similarity value, where the in silico spectrum is used for the library template, is used as the correlation coefficient value. The candidates are ranked by the reverse dot product score and the candidate with the highest similarity value is described as the representative C=C isomer candidate for the EAD-MS/MS spectrum. |

#### 23) PC[M+Na]+ / Lipid quantification

| | | | |
|---|---|---|---|
| Quantitative | No | Batch correction | No |
| Normalization to reference | No | Further quantification remarks | - |

#### 24) PE[M+H]+ / Lipid identification

| | | | |
|---|---|---|---|
| Lipid class | PE | Did you presume assumptions for identification? | No |
| Derivatization | - | Check isomer overlap | No |
| MS Level for identification | MS1, MS2 | RT verified by standard | Yes |
| Identification level | Double bond position | Separation of isobaric/isomeric interferece confirmed | Yes |
| Polarity mode | Positive | Model for separation prediction | Yes |
| Type of positive (precursor)ion | [M+H]+ | Additional dimension/techniques | EAD |
| Fragments for identification | | How was/were the additional dimension(s) used? | To determine sn- and doublebond positions |

| Fragment name |
|---|
| -HG(PE,141) |
| HG+ |
| HG + C3H6+ |
| HG + C2H4O+ |
| NL of FA |
| NL of HG and FA |

| | | | |
|---|---|---|---|
| Isotope correction at MS1 | No | Was a model used to predict lipid molecule separation? | No |
| Isotope correction at MS2 | No | Lipid Identification Software | MS-DIAL |
| MS1 verified by standard | Yes | Data manipulation | Smoothing, Centroiding |
| MS2 verified by standard | Yes | Nomenclature for intact lipid molecule | Yes |
| Background check at MS1 | Yes | Nomenclature for fragment ions | No |
| Background check at MS2 | No | Further identification remarks | The reverse dot product similarity value, where the in silico spectrum is used for the library template, is used as the correlation coefficient value. The candidates are ranked by the reverse dot product score and the candidate with the highest similarity value is described as the representative C=C isomer candidate for the EAD-MS/MS spectrum. |

#### 24) PE[M+H]+ / Lipid quantification

| | | | |
|---|---|---|---|
| Quantitative | No | Batch correction | No |
| Normalization to reference | No | Further quantification remarks | - |

#### 25) PE[M+Na]+ / Lipid identification

| | | | |
|---|---|---|---|
| Lipid class | PE | Did you presume assumptions for identification? | No |
| Derivatization | - | Check isomer overlap | No |
| MS Level for identification | MS1, MS2 | RT verified by standard | Yes |
| Identification level | Double bond position | Separation of isobaric/isomeric interferece confirmed | Yes |
| Polarity mode | Positive | Model for separation prediction | Yes |
| Type of positive (precursor)ion | [M+Na]+ | Additional dimension/techniques | EAD |
| Fragments for identification | Fragment name: -HG(PE,141); HG+Na+; HG + C3H5Na+; HG + C2H3ONa+; NL of FA | How was/were the additional dimension(s) used? | To determine sn- and doublebond positions |
| Isotope correction at MS1 | No | Was a model used to predict lipid molecule separation? | No |
| Isotope correction at MS2 | No | Lipid Identification Software | MS-DIAL |
| MS1 verified by standard | Yes | Data manipulation | Smoothing, Centroiding |
| MS2 verified by standard | Yes | Nomenclature for intact lipid molecule | Yes |
| Background check at MS1 | Yes | Nomenclature for fragment ions | No |
| Background check at MS2 | No | Further identification remarks | The reverse dot product similarity value, where the in silico spectrum is used for the library template, is used as the correlation coefficient value. The candidates are ranked by the reverse dot product score and the candidate with the highest similarity value is described as the representative C=C isomer candidate for the EAD-MS/MS spectrum. |

#### 25) PE[M+Na]+ / Lipid quantification

| | | | |
|---|---|---|---|
| Quantitative | No | Batch correction | No |
| Normalization to reference | No | Further quantification remarks | - |

#### 26) PG[M+NH4]+ / Lipid identification

| | | | |
|---|---|---|---|
| Lipid class | PG | Did you presume assumptions for identification? | No |
| Derivatization | - | Check isomer overlap | No |
| MS Level for identification | MS1, MS2 | RT verified by standard | Yes |
| Identification level | Double bond position | Separation of isobaric/isomeric interferece confirmed | Yes |
| Polarity mode | Positive | Model for separation prediction | Yes |
| Type of positive (precursor)ion | [M+NH4]+ | Additional dimension/techniques | EAD |
| Fragments for identification | | How was/were the additional dimension(s) used? | To determine doublebond positions |

| Fragment name |
|---|
| -HG(PG,172) |
| HG+ |
| NL of FA |
| NL of HG and FA |
| HG + C3H6+ |
| HG + C2H4O+ |

| | | | |
|---|---|---|---|
| Isotope correction at MS1 | No | Was a model used to predict lipid molecule separation? | No |
| Isotope correction at MS2 | No | Lipid Identification Software | MS-DIAL |
| MS1 verified by standard | Yes | Data manipulation | Smoothing, Centroiding |
| MS2 verified by standard | Yes | Nomenclature for intact lipid molecule | Yes |
| Background check at MS1 | Yes | Nomenclature for fragment ions | No |
| Background check at MS2 | No | Further identification remarks | The reverse dot product similarity value, where the in silico spectrum is used for the library template, is used as the correlation coefficient value. The candidates are ranked by the reverse dot product score and the candidate with the highest similarity value is described as the representative C=C isomer candidate for the EAD-MS/MS spectrum. |

#### 26) PG[M+NH4]+ / Lipid quantification

| | | | |
|---|---|---|---|
| Quantitative | No | Batch correction | No |
| Normalization to reference | No | Further quantification remarks | - |

#### 27) PG[M+Na]+ / Lipid identification

| | | | |
|---|---|---|---|
| Lipid class | PG | Did you presume assumptions for identification? | No |
| Derivatization | - | Check isomer overlap | No |
| MS Level for identification | MS1, MS2 | RT verified by standard | Yes |
| Identification level | Double bond position | Separation of isobaric/isomeric interferece confirmed | Yes |
| Polarity mode | Positive | Model for separation prediction | Yes |
| Type of positive (precursor)ion | [M+Na]+ | Additional dimension/techniques | EAD |
| Fragments for identification | | How was/were the additional dimension(s) used? | To determine sn- and doublebond positions |
| Isotope correction at MS1 | No | Was a model used to predict lipid molecule separation? | No |
| Isotope correction at MS2 | No | Lipid Identification Software | MS-DIAL |
| MS1 verified by standard | Yes | Data manipulation | Smoothing, Centroiding |
| MS2 verified by standard | Yes | Nomenclature for intact lipid molecule | Yes |
| Background check at MS1 | Yes | Nomenclature for fragment ions | No |
| Background check at MS2 | No | Further identification remarks | The reverse dot product similarity value, where the in silico spectrum is used for the library template, is used as the correlation coefficient value. The candidates are ranked by the reverse dot product score and the candidate with the highest similarity value is described as the representative C=C isomer candidate for the EAD-MS/MS spectrum. |

| Fragment name |
|---|
| -HG(PG,172) |
| HG + Na+ |
| NL of FA |
| HG + C3H5Na+ |
| HG + C2H3ONa+ |

#### 27) PG[M+Na]+ / Lipid quantification

| | | | |
|---|---|---|---|
| Quantitative | No | Batch correction | No |
| Normalization to reference | No | Further quantification remarks | - |

#### 28) PI[M+NH4]+ / Lipid identification

| | | | |
|---|---|---|---|
| Lipid class | PI | Did you presume assumptions for identification? | No |
| Derivatization | - | Check isomer overlap | No |
| MS Level for identification | MS1, MS2 | RT verified by standard | Yes |
| Identification level | Double bond position | Separation of isobaric/isomeric interferece confirmed | Yes |
| Polarity mode | Positive | Model for separation prediction | Yes |
| Type of positive (precursor)ion | [M+NH4]+ | Additional dimension/techniques | EAD |
| Fragments for identification | | How was/were the additional dimension(s) used? | To determine doublebond positions |

| Fragment name |
|---|
| HG+ |
| HG + C3H5 |
| HG + C2H3O |
| NL of HG and FA |
| NL of FA |
| NL of inositol |
| -HG(PI,260) |

| | | | |
|---|---|---|---|
| Isotope correction at MS1 | No | Was a model used to predict lipid molecule separation? | No |
| Isotope correction at MS2 | No | Lipid Identification Software | MS-DIAL |
| MS1 verified by standard | Yes | Data manipulation | Smoothing, Centroiding |
| MS2 verified by standard | Yes | Nomenclature for intact lipid molecule | Yes |
| Background check at MS1 | Yes | Nomenclature for fragment ions | No |
| Background check at MS2 | No | Further identification remarks | The reverse dot product similarity value, where the in silico spectrum is used for the library template, is used as the correlation coefficient value. The candidates are ranked by the reverse dot product score and the candidate with the highest similarity value is described as the representative C=C isomer candidate for the EAD-MS/MS spectrum. |

#### 28) PI[M+NH4]+ / Lipid quantification

| | | | |
|---|---|---|---|
| Quantitative | No | Batch correction | No |
| Normalization to reference | No | Further quantification remarks | - |

#### 29) PI[M+Na]+ / Lipid identification

| | | | |
|---|---|---|---|
| Lipid class | PI | Did you presume assumptions for identification? | No |
| Derivatization | - | Check isomer overlap | No |
| MS Level for identification | MS1, MS2 | RT verified by standard | Yes |
| Identification level | sn Position | Separation of isobaric/isomeric interferece confirmed | Yes |
| Polarity mode | Positive | Model for separation prediction | Yes |
| Type of positive (precursor)ion | [M+Na]+ | Additional dimension/techniques | EAD |
| Fragments for identification | **Fragment name**<br>HG + Na+<br>HG + C3H5Na+<br>HG + C2H3ONa+<br>NL of FA<br>NL of inositol | How was/were the additional dimension(s) used? | To determine sn- position |
| Isotope correction at MS1 | No | Was a model used to predict lipid molecule separation? | No |
| Isotope correction at MS2 | No | Lipid Identification Software | MS-DIAL |
| MS1 verified by standard | Yes | Data manipulation | Smoothing, Centroiding |
| MS2 verified by standard | Yes | Nomenclature for intact lipid molecule | Yes |
| Background check at MS1 | Yes | Nomenclature for fragment ions | No |
| Background check at MS2 | No | Further identification remarks | The reverse dot product similarity value, where the in silico spectrum is used for the library template, is used as the correlation coefficient value. The candidates are ranked by the reverse dot product score and the candidate with the highest similarity value is described as the representative C=C isomer candidate for the EAD-MS/MS spectrum. |

#### 29) PI[M+Na]+ / Lipid quantification

| | | | |
|---|---|---|---|
| Quantitative | No | Batch correction | No |
| Normalization to reference | No | Further quantification remarks | - |

#### 30) PS[M+H]+ / Lipid identification

| | | | |
|---|---|---|---|
| Lipid class | PS | Did you presume assumptions for identification? | No |
| Derivatization | - | Check isomer overlap | No |
| MS Level for identification | MS1, MS2 | RT verified by standard | Yes |
| Identification level | Double bond position | Separation of isobaric/isomeric interferece confirmed | Yes |
| Polarity mode | Positive | Model for separation prediction | Yes |
| Type of positive (precursor)ion | [M+H]+ | Additional dimension/techniques | EAD |
| Fragments for identification | | How was/were the additional dimension(s) used? | To determine doublebond positions |

| Fragment name |
|---|
| -HG(PS,185) |
| HG+ |
| NL of FA |
| NL of HG and FA |
| HG + C3H6+ |
| HG + C2H4O+ |

| | | | |
|---|---|---|---|
| Isotope correction at MS1 | No | Was a model used to predict lipid molecule separation? | No |
| Isotope correction at MS2 | No | Lipid Identification Software | MS-DIAL |
| MS1 verified by standard | Yes | Data manipulation | Smoothing, Centroiding |
| MS2 verified by standard | Yes | Nomenclature for intact lipid molecule | Yes |
| Background check at MS1 | Yes | Nomenclature for fragment ions | No |
| Background check at MS2 | No | Further identification remarks | The reverse dot product similarity value, where the in silico spectrum is used for the library template, is used as the correlation coefficient value. The candidates are ranked by the reverse dot product score and the candidate with the highest similarity value is described as the representative C=C isomer candidate for the EAD-MS/MS spectrum. |

#### 30) PS[M+H]+ / Lipid quantification

| | | | |
|---|---|---|---|
| Quantitative | No | Batch correction | No |
| Normalization to reference | No | Further quantification remarks | - |

#### 31) PS[M+Na]+ / Lipid identification

| | | | |
|---|---|---|---|
| Lipid class | PS | Did you presume assumptions for identification? | No |
| Derivatization | - | Check isomer overlap | No |
| MS Level for identification | MS1, MS2 | RT verified by standard | Yes |
| Identification level | sn Position | Separation of isobaric/isomeric interferece confirmed | Yes |
| Polarity mode | Positive | Model for separation prediction | Yes |
| Type of positive (precursor)ion | [M+Na]+ | Additional dimension/techniques | EAD |
| Fragments for identification | | How was/were the additional dimension(s) used? | To determine sn- position |

| Fragment name |
|---|
| -HG(PS,185) |
| HG + Na+ |
| NL of FA |
| HG + C3H5Na+ |
| HG + C2H3ONa+ |
| NL of HG and FA |

| | | | |
|---|---|---|---|
| Isotope correction at MS1 | No | Was a model used to predict lipid molecule separation? | No |
| Isotope correction at MS2 | No | Lipid Identification Software | MS-DIAL |
| MS1 verified by standard | Yes | Data manipulation | Smoothing, Centroiding |
| MS2 verified by standard | Yes | Nomenclature for intact lipid molecule | Yes |
| Background check at MS1 | Yes | Nomenclature for fragment ions | No |
| Background check at MS2 | No | Further identification remarks | The reverse dot product similarity value, where the in silico spectrum is used for the library template, is used as the correlation coefficient value. The candidates are ranked by the reverse dot product score and the candidate with the highest similarity value is described as the representative C=C isomer candidate for the EAD-MS/MS spectrum. |

#### 31) PS[M+Na]+ / Lipid quantification

| | | | |
|---|---|---|---|
| Quantitative | No | Batch correction | No |
| Normalization to reference | No | Further quantification remarks | - |

#### 32) TG[M+NH4]+ / Lipid identification

| | | | |
|---|---|---|---|
| Lipid class | TG | Did you presume assumptions for identification? | No |
| Derivatization | - | Check isomer overlap | No |
| MS Level for identification | MS1, MS2 | RT verified by standard | Yes |
| Identification level | Double bond position | Separation of isobaric/isomeric interferece confirmed | Yes |
| Polarity mode | Positive | Model for separation prediction | Yes |
| Type of positive (precursor)ion | [M+NH4]+ | Additional dimension/techniques | EAD |
| Fragments for identification | Fragment name: NL of FA; FA + C3H5O2+; Fatty Acyl | How was/were the additional dimension(s) used? | To determine doublebond positions |
| Isotope correction at MS1 | No | Was a model used to predict lipid molecule separation? | No |
| Isotope correction at MS2 | No | Lipid Identification Software | MS-DIAL |
| MS1 verified by standard | Yes | Data manipulation | Smoothing, Centroiding |
| MS2 verified by standard | Yes | Nomenclature for intact lipid molecule | Yes |
| Background check at MS1 | Yes | Nomenclature for fragment ions | No |
| Background check at MS2 | No | Further identification remarks | The reverse dot product similarity value, where the in silico spectrum is used for the library template, is used as the correlation coefficient value. The candidates are ranked by the reverse dot product score and the candidate with the highest similarity value is described as the representative C=C isomer candidate for the EAD-MS/MS spectrum. |

#### 32) TG[M+NH4]+ / Lipid quantification

| | | | |
|---|---|---|---|
| Quantitative | No | Batch correction | No |
| Normalization to reference | No | Further quantification remarks | - |

#### 33) TG[M+Na]+ / Lipid identification

| | | | |
|---|---|---|---|
| Lipid class | TG | Did you presume assumptions for identification? | No |
| Derivatization | - | Check isomer overlap | No |
| MS Level for identification | MS1, MS2 | RT verified by standard | Yes |
| Identification level | Double bond position | Separation of isobaric/isomeric interferece confirmed | Yes |
| Polarity mode | Positive | Model for separation prediction | Yes |
| Type of positive (precursor)ion | [M+Na]+ | Additional dimension/techniques | EAD |
| Fragments for identification | Fragment name: NL of FA; Fatty Acyl; FA + C3H5ONa+; FA + C3H3O2Na+ | How was/were the additional dimension(s) used? | To determine sn- and doublebond positions |
| Isotope correction at MS1 | No | Was a model used to predict lipid molecule separation? | No |
| Isotope correction at MS2 | No | Lipid Identification Software | MS-DIAL |
| MS1 verified by standard | Yes | Data manipulation | Smoothing, Centroiding |
| MS2 verified by standard | Yes | Nomenclature for intact lipid molecule | Yes |
| Background check at MS1 | Yes | Nomenclature for fragment ions | No |
| Background check at MS2 | No | Further identification remarks | The reverse dot product similarity value, where the in silico spectrum is used for the library template, is used as the correlation coefficient value. The candidates are ranked by the reverse dot product score and the candidate with the highest similarity value is described as the representative C=C isomer candidate for the EAD-MS/MS spectrum. |

#### 33) TG[M+Na]+ / Lipid quantification

| | | | |
|---|---|---|---|
| Quantitative | No | Batch correction | No |
| Normalization to reference | No | Further quantification remarks | - |

#### 34) Ceramide non-hydroxyfatty acid-sphingosine (Cer_NS)[M+H]+ / Lipid identification

| | | | |
|---|---|---|---|
| Lipid class | Ceramide non-hydroxyfatty acid-sphingosine (Cer_NS) | Did you presume assumptions for identification? | No |
| Derivatization | - | Check isomer overlap | No |
| MS Level for identification | MS1, MS2 | RT verified by standard | Yes |
| Identification level | Double bond position | Separation of isobaric/isomeric interferece confirmed | Yes |
| Polarity mode | Positive | Model for separation prediction | Yes |
| Type of positive (precursor)ion | [M+H]+ | Additional dimension/techniques | EAD |
| Fragments for identification | | How was/were the additional dimension(s) used? | To determine doublebond positions |

| Fragment name |
|---|
| Neutral loss of H2O |
| Neutral loss of 2H2O |
| Neutral loss of CH4O2 |
| Fatty acyl amide +C2H3O+ fragment |
| Fatty acyl amide +C2H2+ fragment |
| Fatty acyl amide+ fragment |
| Sphingosine -CH4O2 fragment |
| Sphingosine -2H2O fragment |
| Sphingosine -H2O fragment |

| | | | |
|---|---|---|---|
| Isotope correction at MS1 | No | Was a model used to predict lipid molecule separation? | No |
| Isotope correction at MS2 | No | Lipid Identification Software | MS-DIAL |
| MS1 verified by standard | Yes | Data manipulation | Smoothing, Centroiding |
| MS2 verified by standard | Yes | Nomenclature for intact lipid molecule | Yes |
| Background check at MS1 | Yes | Nomenclature for fragment ions | No |
| Background check at MS2 | No | Further identification remarks | The reverse dot product similarity value, where the in silico spectrum is used for the library template, is used as the correlation coefficient value. The candidates are ranked by the reverse dot product score and the candidate with the highest similarity value is described as the representative C=C isomer candidate for the EAD-MS/MS spectrum. |

#### 34) Ceramide non-hydroxyfatty acid-sphingosine (Cer_NS)[M+H]+ / Lipid quantification

| | | | |
|---|---|---|---|
| Quantitative | No | Batch correction | No |
| Normalization to reference | No | Further quantification remarks | - |

#### 35) Ceramide non-hydroxyfatty acid-sphingosine (Cer_NS)[M+Na]+ / Lipid identification

| | | | |
|---|---|---|---|
| Lipid class | Ceramide non-hydroxyfatty acid-sphingosine (Cer_NS) | Did you presume assumptions for identification? | No |
| Derivatization | - | Check isomer overlap | No |
| MS Level for identification | MS1, MS2 | RT verified by standard | Yes |
| Identification level | Double bond position | Separation of isobaric/isomeric interference confirmed | Yes |
| Polarity mode | Positive | Model for separation prediction | Yes |
| Type of positive (precursor)ion | [M+Na]+ | Additional dimension/techniques | EAD |
| Fragments for identification | | How was/were the additional dimension(s) used? | To determine doublebond positions |

| Fragment name |
|---|
| Neutral loss of CH3O |
| Fatty acyl amide +C2H2ONa fragment |

The reverse dot product similarity value, where the in silico spectrum is used for the library template, is used as the correlation coefficient value. The candidates are ranked by the reverse dot product score and the candidate with the highest similarity value is described as the representative C=C isomer candidate for the EAD-MS/MS spectrum.

| | | | |
|---|---|---|---|
| Isotope correction at MS1 | No | Was a model used to predict lipid molecule separation? | No |
| Isotope correction at MS2 | No | Lipid Identification Software | MS-DIAL |
| MS1 verified by standard | Yes | Data manipulation | Smoothing, Centroiding |
| MS2 verified by standard | Yes | Nomenclature for intact lipid molecule | Yes |
| Background check at MS1 | Yes | Nomenclature for fragment ions | No |
| Background check at MS2 | No | Further identification remarks | The reverse dot product similarity value, where the in silico spectrum is used for the library template, is used as the correlation coefficient value. The candidates are ranked by the reverse dot product score and the candidate with the highest similarity value is described as the representative C=C isomer candidate for the EAD-MS/MS spectrum. |

#### 35) Ceramide non-hydroxyfatty acid-sphingosine (Cer_NS)[M+Na]+ / Lipid quantification

| | | | |
|---|---|---|---|
| Quantitative | No | Batch correction | No |
| Normalization to reference | No | Further quantification remarks | - |

#### 36) Ceramide non-hydroxyfatty acid-dihydrosphingosine (Cer_NDS)[M+H]+ / Lipid identification

| | | | |
|---|---|---|---|
| Lipid class | Ceramide non-hydroxyfatty acid-dihydrosphingosine (Cer_NDS) | Did you presume assumptions for identification? | No |
| Derivatization | - | Check isomer overlap | No |
| MS Level for identification | MS1, MS2 | RT verified by standard | Yes |
| Identification level | Double bond position | Separation of isobaric/isomeric interferece confirmed | Yes |
| Polarity mode | Positive | Model for separation prediction | Yes |
| Type of positive (precursor)ion | [M+H]+ | Additional dimension/techniques | EAD |
| Fragments for identification | | How was/were the additional dimension(s) used? | To determine doublebond positions |

| Fragment name |
|---|
| Neutral loss of H2O |
| Neutral loss of 2H2O |
| Neutral loss of CH4O2 |
| Fatty acyl amide +C2H3O+ fragment |
| Fatty acyl amide +C2H2+ fragment |
| Fatty acyl amide+ fragment |
| Sphingosine -CH4O2 fragment |
| Sphingosine -2H2O fragment |
| Sphingosine -H2O fragment |

| | | | |
|---|---|---|---|
| Isotope correction at MS1 | No | Was a model used to predict lipid molecule separation? | No |
| Isotope correction at MS2 | No | Lipid Identification Software | MS-DIAL |
| MS1 verified by standard | No | Data manipulation | Smoothing, Centroiding |
| MS2 verified by standard | No | Nomenclature for intact lipid molecule | Yes |
| Background check at MS1 | Yes | Nomenclature for fragment ions | No |
| Background check at MS2 | No | Further identification remarks | The reverse dot product similarity value, where the in silico spectrum is used for the library template, is used as the correlation coefficient value. The candidates are ranked by the reverse dot product score and the candidate with the highest similarity value is described as the representative C=C isomer candidate for the EAD-MS/MS spectrum. |

#### 36) Ceramide non-hydroxyfatty acid-dihydrosphingosine (Cer_NDS)[M+H]+ / Lipid quantification

| | | | |
|---|---|---|---|
| Quantitative | No | Batch correction | No |
| Normalization to reference | No | Further quantification remarks | - |

#### 37) Ceramide non-hydroxyfatty acid-dihydrosphingosine (Cer_NDS)[M+Na]+ / Lipid identification

| | | | |
|---|---|---|---|
| Lipid class | Ceramide non-hydroxyfatty acid-dihydrosphingosine (Cer_NDS) | Did you presume assumptions for identification? | No |
| Derivatization | - | Check isomer overlap | No |
| MS Level for identification | MS1, MS2 | RT verified by standard | Yes |
| Identification level | Double bond position | Separation of isobaric/isomeric interference confirmed | Yes |
| Polarity mode | Positive | Model for separation prediction | Yes |
| Type of positive (precursor)ion | [M+Na]+ | Additional dimension/techniques | EAD |
| Fragments for identification | | How was/were the additional dimension(s) used? | To determine doublebond positions |
| Isotope correction at MS1 | No | Was a model used to predict lipid molecule separation? | No |
| Isotope correction at MS2 | No | Lipid Identification Software | MS-DIAL |
| MS1 verified by standard | No | Data manipulation | Smoothing, Centroiding |
| MS2 verified by standard | No | Nomenclature for intact lipid molecule | Yes |
| Background check at MS1 | Yes | Nomenclature for fragment ions | No |
| Background check at MS2 | No | Further identification remarks | The reverse dot product similarity value, where the in silico spectrum is used for the library template, is used as the correlation coefficient value. The candidates are ranked by the reverse dot product score and the candidate with the highest similarity value is described as the representative C=C isomer candidate for the EAD-MS/MS spectrum. |

Fragments for identification:

| Fragment name |
|---|
| Neutral loss of CH3O |
| Fatty acyl amide +C2H2ONa fragment |

The reverse dot product similarity value, where the in silico spectrum is used for the library template, is used as the correlation coefficient value. The candidates are ranked by the reverse dot product score and the candidate with the highest similarity value is described as the representative C=C isomer candidate for the EAD-MS/MS spectrum.

#### 37) Ceramide non-hydroxyfatty acid-dihydrosphingosine (Cer_NDS)[M+Na]+ / Lipid quantification

| | | | |
|---|---|---|---|
| Quantitative | No | Batch correction | No |
| Normalization to reference | No | Further quantification remarks | - |

#### 38) Ceramide non-hydroxyfatty acid-phytospingosine(Cer_NP)[M+H]+ / Lipid identification

| | | | |
|---|---|---|---|
| Lipid class | Ceramide non-hydroxyfatty acid-phytospingosine(Cer_NP) | Did you presume assumptions for identification? | No |
| Derivatization | - | Check isomer overlap | No |
| MS Level for identification | MS1, MS2 | RT verified by standard | Yes |
| Identification level | Double bond position | Separation of isobaric/isomeric interferece confirmed | Yes |
| Polarity mode | Positive | Model for separation prediction | Yes |
| Type of positive (precursor)ion | [M+H]+ | Additional dimension/techniques | EAD |
| Fragments for identification | | How was/were the additional dimension(s) used? | To determine doublebond positions |

| Fragment name |
|---|
| Neutral loss of H2O |
| Neutral loss of 2H2O |
| Neutral loss of CH4O2 |
| Fatty acyl amide +C2H3O+ fragment |
| Fatty acyl amide +C2H2+ fragment |
| Fatty acyl amide+ fragment |
| Phytosphingosine -CH4O2 fragment |
| Phytosphingosine -2H2O fragment |
| Phytosphingosine -H2O fragment |
| Fatty acyl amide +C3H4O+ fragment |
| Fatty acyl amide +C3H5O2+ fragment |
| Phytosphingosine -CH6O2 fragment |

| | | | |
|---|---|---|---|
| Isotope correction at MS1 | No | Was a model used to predict lipid molecule separation? | No |
| Isotope correction at MS2 | No | Lipid Identification Software | MS-DIAL |
| MS1 verified by standard | Yes | Data manipulation | Smoothing, Centroiding |
| MS2 verified by standard | Yes | Nomenclature for intact lipid molecule | Yes |
| Background check at MS1 | Yes | Nomenclature for fragment ions | No |
| Background check at MS2 | No | Further identification remarks | The reverse dot product similarity value, where the in silico spectrum is used for the library template, is used as the correlation coefficient value. The candidates are ranked by the reverse dot product score and the candidate with the highest similarity value is described as the representative C=C isomer candidate for the EAD-MS/MS spectrum. |

#### 38) Ceramide non-hydroxyfatty acid-phytospingosine(Cer_NP)[M+H]+ / Lipid quantification

| | | | |
|---|---|---|---|
| Quantitative | No | Batch correction | No |
| Normalization to reference | No | Further quantification remarks | - |

#### 39) Ceramide alpha-hydroxy fatty acid-sphingosine (Cer_AS)[M+H]+ / Lipid identification

| | | | |
|---|---|---|---|
| Lipid class | Ceramide alpha-hydroxy fatty acid-sphingosine (Cer_AS) | Did you presume assumptions for identification? | No |
| Derivatization | - | Check isomer overlap | No |
| MS Level for identification | MS1, MS2 | RT verified by standard | Yes |
| Identification level | Double bond position | Separation of isobaric/isomeric interferece confirmed | Yes |
| Polarity mode | Positive | Model for separation prediction | Yes |
| Type of positive (precursor)ion | [M+H]+ | Additional dimension/techniques | EAD |
| Fragments for identification | | How was/were the additional dimension(s) used? | To determine doublebond positions |

| Fragment name |
|---|
| Neutral loss of H2O |
| Neutral loss of 2H2O |
| Neutral loss of CH4O2 |
| Alpha-hydroxy fatty acyl amide +C2H3O+ fragment |
| Alpha-hydroxy fatty acyl amide +C2H2+ fragment |
| Alpha-hydroxy fatty acyl amide+ fragment |
| Sphingosine -CH4O2 fragment |
| Sphingosine -2H2O fragment |
| Sphingosine -H2O fragment |

| | | | |
|---|---|---|---|
| Isotope correction at MS1 | No | Was a model used to predict lipid molecule separation? | No |
| Isotope correction at MS2 | No | Lipid Identification Software | MS-DIAL |
| MS1 verified by standard | Yes | Data manipulation | Smoothing, Centroiding |
| MS2 verified by standard | Yes | Nomenclature for intact lipid molecule | Yes |
| Background check at MS1 | Yes | Nomenclature for fragment ions | No |
| Background check at MS2 | No | Further identification remarks | The reverse dot product similarity value, where the in silico spectrum is used for the library template, is used as the correlation coefficient value. The candidates are ranked by the reverse dot product score and the candidate with the highest similarity value is described as the representative C=C isomer candidate for the EAD-MS/MS spectrum. |

#### 39) Ceramide alpha-hydroxy fatty acid-sphingosine (Cer_AS)[M+H]+ / Lipid quantification

| | | | |
|---|---|---|---|
| Quantitative | No | Batch correction | No |
| Normalization to reference | No | Further quantification remarks | - |

#### 40) Hexosylceramide non-hydroxyfatty acid-sphingosine (HexCer_NS)[M+H]+ / Lipid identification

| | | | |
|---|---|---|---|
| Lipid class | Hexosylceramide non-hydroxyfatty acid-sphingosine (HexCer_NS) | Did you presume assumptions for identification? | No |
| Derivatization | - | Check isomer overlap | No |
| MS Level for identification | MS1, MS2 | RT verified by standard | Yes |
| Identification level | Double bond position | Separation of isobaric/isomeric interferece confirmed | Yes |
| Polarity mode | Positive | Model for separation prediction | Yes |
| Type of positive (precursor)ion | [M+H]+ | Additional dimension/techniques | EAD |
| Fragments for identification | | How was/were the additional dimension(s) used? | To determine doublebond positions |

| Fragment name |
|---|
| Neutral loss of H2O |
| Neutral loss of hexose |
| Neutral loss of hexose and H2O |
| Neutral loss of hexose and 2H2O |
| Fatty acyl amide +C2H5O + hexose fragment |
| Fatty acyl amide +C2H3O+ fragment |
| Fatty acyl amide +C2H2+ fragment |
| Fatty acyl amide+ fragment |
| Sphingosine -CH4O2 fragment |
| Sphingosine -2H2O fragment |
| Sphingosine -H2O fragment |
| Hexose + C2H5N+ |

| | | | |
|---|---|---|---|
| Isotope correction at MS1 | No | Was a model used to predict lipid molecule separation? | No |
| Isotope correction at MS2 | No | Lipid Identification Software | MS-DIAL |
| MS1 verified by standard | Yes | Data manipulation | Smoothing, Centroiding |
| MS2 verified by standard | Yes | Nomenclature for intact lipid molecule | Yes |
| Background check at MS1 | Yes | Nomenclature for fragment ions | No |
| Background check at MS2 | No | Further identification remarks | The reverse dot product similarity value, where the in silico spectrum is used for the library template, is used as the correlation coefficient value. The candidates are ranked by the reverse dot product score and the candidate with the highest similarity value is described as the representative C=C isomer candidate for the EAD-MS/MS spectrum. |

#### 40) Hexosylceramide non-hydroxyfatty acid-sphingosine (HexCer_NS)[M+H]+ / Lipid quantification

| | | | |
|---|---|---|---|
| Quantitative | No | Batch correction | No |
| Normalization to reference | No | Further quantification remarks | - |

#### 41) Hexosylceramide non-hydroxyfatty acid-sphingosine (HexCer_NS)[M+Na]+ / Lipid identification

| | | | |
|---|---|---|---|
| Lipid class | Hexosylceramide non-hydroxyfatty acid-sphingosine (HexCer_NS) | Did you presume assumptions for identification? | No |
| Derivatization | - | Check isomer overlap | No |
| MS Level for identification | MS1, MS2 | RT verified by standard | Yes |
| Identification level | Double bond position | Separation of isobaric/isomeric interference confirmed | Yes |
| Polarity mode | Positive | Model for separation prediction | Yes |
| Type of positive (precursor)ion | [M+Na]+ | Additional dimension/techniques | EAD |
| Fragments for identification | (see fragment table below) | How was/were the additional dimension(s) used? | To determine doublebond positions |
| Isotope correction at MS1 | No | Was a model used to predict lipid molecule separation? | No |
| Isotope correction at MS2 | No | Lipid Identification Software | MS-DIAL |
| MS1 verified by standard | Yes | Data manipulation | Smoothing, Centroiding |
| MS2 verified by standard | Yes | Nomenclature for intact lipid molecule | Yes |
| Background check at MS1 | Yes | Nomenclature for fragment ions | No |
| Background check at MS2 | No | Further identification remarks | The reverse dot product similarity value, where the in silico spectrum is used for the library template, is used as the correlation coefficient value. The candidates are ranked by the reverse dot product score and the candidate with the highest similarity value is described as the representative C=C isomer candidate for the EAD-MS/MS spectrum. |

| Fragment name |
|---|
| Neutral loss of C5H10O4 |
| Neutral loss of hexose and H2O |
| Fatty acyl amide +C2H3ONa+ fragment |
| Fatty acyl amide +C2HNa+ fragment |
| Fatty acyl amide+ fragment |
| Hexose + C2H5NNa+ |
| C5H9O4Na+ |

#### 41) Hexosylceramide non-hydroxyfatty acid-sphingosine (HexCer_NS)[M+Na]+ / Lipid quantification

| | | | |
|---|---|---|---|
| Quantitative | No | Batch correction | No |
| Normalization to reference | No | Further quantification remarks | - |

#### 42) Di-hexosylceramide hydroxyfatty acid-sphingosine (Hex2Cer_NS)[M+H]+ / Lipid identification

| | | | |
|---|---|---|---|
| Lipid class | Di-hexosylceramide hydroxyfatty acid-sphingosine (Hex2Cer_NS) | Did you presume assumptions for identification? | No |
| Derivatization | - | Check isomer overlap | No |
| MS Level for identification | MS1, MS2 | RT verified by standard | Yes |
| Identification level | Molecular species level | Separation of isobaric/isomeric interferece confirmed | Yes |
| Polarity mode | Positive | Model for separation prediction | Yes |
| Type of positive (precursor)ion | [M+H]+ | Additional dimension/techniques | EAD |
| Fragments for identification | | How was/were the additional dimension(s) used? | To determine OH positions |

| Fragment name |
|---|
| Neutral loss of H2O |
| Neutral loss of 2hexose |
| Neutral loss of 2hexose and H2O |
| Neutral loss of 2hexose and 2H2O |
| Fatty acyl amide +C2H5O + 2hexose fragment |
| Fatty acyl amide +C2H3O+ fragment |
| Fatty acyl amide +C2H2+ fragment |
| Fatty acyl amide+ fragment |
| Sphingosine -CH4O2 fragment |
| Sphingosine -2H2O fragment |
| Sphingosine -H2O fragment |

| | | | |
|---|---|---|---|
| Isotope correction at MS1 | No | Was a model used to predict lipid molecule separation? | No |
| Isotope correction at MS2 | No | Lipid Identification Software | MS-DIAL |
| MS1 verified by standard | Yes | Data manipulation | Smoothing, Centroiding |
| MS2 verified by standard | Yes | Nomenclature for intact lipid molecule | Yes |
| Background check at MS1 | Yes | Nomenclature for fragment ions | No |
| Background check at MS2 | No | Further identification remarks | The reverse dot product similarity value, where the in silico spectrum is used for the library template, is used as the correlation coefficient value. The candidates are ranked by the reverse dot product score and the candidate with the highest similarity value is described as the representative C=C isomer candidate for the EAD-MS/MS spectrum. |

#### 42) Di-hexosylceramide hydroxyfatty acid-sphingosine (Hex2Cer_NS)[M+H]+ / Lipid quantification

| | | | |
|---|---|---|---|
| Quantitative | No | Batch correction | No |
| Normalization to reference | No | Further quantification remarks | - |

#### 43) SHexCer[M+H]+ / Lipid identification

| | | | |
|---|---|---|---|
| Lipid class | SHexCer | Did you presume assumptions for identification? | No |
| Derivatization | - | Check isomer overlap | No |
| MS Level for identification | MS1, MS2 | RT verified by standard | Yes |
| Identification level | Molecular species level | Separation of isobaric/isomeric interferece confirmed | Yes |
| Polarity mode | Positive | Model for separation prediction | Yes |
| Type of positive (precursor)ion | [M+H]+ | Additional dimension/techniques | EAD |
| Fragments for identification | | How was/were the additional dimension(s) used? | To determine OH positions |

| Fragment name |
|---|
| Neutral loss of SO3 |
| -HG(SHex,98) |
| -HG(SHex,260) |
| -HG(SHex,278) |
| -HG(SHex,242) |
| Fatty acyl amide +C2H2+ fragment |
| Fatty acyl amide+ fragment |
| Sphingosine -2H2O fragment |
| Sphingosine -H2O fragment |
| Sphingosine -CH4O2 fragment |
| Hexose + C2H5N+ |

| | | | |
|---|---|---|---|
| Isotope correction at MS1 | No | Was a model used to predict lipid molecule separation? | No |
| Isotope correction at MS2 | No | Lipid Identification Software | MS-DIAL |
| MS1 verified by standard | Yes | Data manipulation | Smoothing, Centroiding |
| MS2 verified by standard | Yes | Nomenclature for intact lipid molecule | Yes |
| Background check at MS1 | Yes | Nomenclature for fragment ions | No |
| Background check at MS2 | No | Further identification remarks | The reverse dot product similarity value, where the in silico spectrum is used for the library template, is used as the correlation coefficient value. The candidates are ranked by the reverse dot product score and the candidate with the highest similarity value is described as the representative C=C isomer candidate for the EAD-MS/MS spectrum. |

#### 43) SHexCer[M+H]+ / Lipid quantification

| | | | |
|---|---|---|---|
| Quantitative | No | Batch correction | No |
| Normalization to reference | No | Further quantification remarks | - |

#### 44) SHexCer[M+Na]+ / Lipid identification

| | | | |
|---|---|---|---|
| Lipid class | SHexCer | Did you presume assumptions for identification? | No |
| Derivatization | - | Check isomer overlap | No |
| MS Level for identification | MS1, MS2 | RT verified by standard | Yes |
| Identification level | Molecular species level | Separation of isobaric/isomeric interferece confirmed | Yes |
| Polarity mode | Positive | Model for separation prediction | Yes |
| Type of positive (precursor)ion | [M+Na]+ | Additional dimension/techniques | EAD |
| Fragments for identification | | How was/were the additional dimension(s) used? | To determine OH positions |

| Fragment name |
|---|
| Fatty acyl amide +C2H2Na+ fragment |
| Fatty acyl amide +H3Na+ fragment |
| SHex + C2H4NNa+ |
| Hexose + C2H4NNa+ |
| Hexose + Na+ |
| Neutral loss of SO3 |
| -HG(SHex,98) |
| NL of C5H10O7S |
| NL of C6H10O8S |
| -HG(SHex,260) |

| | | | |
|---|---|---|---|
| Isotope correction at MS1 | No | Was a model used to predict lipid molecule separation? | No |
| Isotope correction at MS2 | No | Lipid Identification Software | MS-DIAL |
| MS1 verified by standard | Yes | Data manipulation | Smoothing, Centroiding |
| MS2 verified by standard | Yes | Nomenclature for intact lipid molecule | Yes |
| Background check at MS1 | Yes | Nomenclature for fragment ions | No |
| Background check at MS2 | No | Further identification remarks | The reverse dot product similarity value, where the in silico spectrum is used for the library template, is used as the correlation coefficient value. The candidates are ranked by the reverse dot product score and the candidate with the highest similarity value is described as the representative C=C isomer candidate for the EAD-MS/MS spectrum. |

#### 44) SHexCer[M+Na]+ / Lipid quantification

| | | | |
|---|---|---|---|
| Quantitative | No | Batch correction | No |
| Normalization to reference | No | Further quantification remarks | - |

#### 45) SM[M+H]+ / Lipid identification

| | | | |
|---|---|---|---|
| Lipid class | SM | Did you presume assumptions for identification? | No |
| Derivatization | - | Check isomer overlap | No |
| MS Level for identification | MS1, MS2 | RT verified by standard | Yes |
| Identification level | Double bond position | Separation of isobaric/isomeric interferece confirmed | Yes |
| Polarity mode | Positive | Model for separation prediction | Yes |
| Type of positive (precursor)ion | [M+H]+ | Additional dimension/techniques | EAD |
| Fragments for identification | | How was/were the additional dimension(s) used? | To determine doublebond positions |

| Fragment name |
|---|
| NL of Fatty acyl and NH4 |
| Fatty acyl amide +C2H2 + Header fragment |
| Fatty acyl amide +C2H2+ fragment |
| Sphingosine -2H2O fragment |
| Header + C3H4NO |
| Header + C2H4N |
| HG(PC,184) |

| | | | |
|---|---|---|---|
| Isotope correction at MS1 | No | Was a model used to predict lipid molecule separation? | No |
| Isotope correction at MS2 | No | Lipid Identification Software | MS-DIAL |
| MS1 verified by standard | Yes | Data manipulation | Smoothing, Centroiding |
| MS2 verified by standard | Yes | Nomenclature for intact lipid molecule | Yes |
| Background check at MS1 | Yes | Nomenclature for fragment ions | No |
| Background check at MS2 | No | Further identification remarks | The reverse dot product similarity value, where the in silico spectrum is used for the library template, is used as the correlation coefficient value. The candidates are ranked by the reverse dot product score and the candidate with the highest similarity value is described as the representative C=C isomer candidate for the EAD-MS/MS spectrum. |

#### 45) SM[M+H]+ / Lipid quantification

| | | | |
|---|---|---|---|
| Quantitative | No | Batch correction | No |
| Normalization to reference | No | Further quantification remarks | - |

#### 46) SM[M+Na]+ / Lipid identification

| | | | |
|---|---|---|---|
| Lipid class | SM | Did you presume assumptions for identification? | No |
| Derivatization | - | Check isomer overlap | No |
| MS Level for identification | MS1, MS2 | RT verified by standard | Yes |
| Identification level | Double bond position | Separation of isobaric/isomeric interferece confirmed | Yes |
| Polarity mode | Positive | Model for separation prediction | Yes |
| Type of positive (precursor)ion | [M+Na]+ | Additional dimension/techniques | EAD |
| Fragments for identification | | How was/were the additional dimension(s) used? | To determine doublebond positions |

| Fragment name |
|---|
| NL of C3H9N |
| NL of C5H11N |
| NL of HG |
| Fatty acyl amide +C2H + header +Na fragment |
| Fatty acyl amide +C2H +Na fragment |
| Header + C2H4NNa |
| HG(PC,184) + Na |

| | | | |
|---|---|---|---|
| Isotope correction at MS1 | No | Was a model used to predict lipid molecule separation? | No |
| Isotope correction at MS2 | No | Lipid Identification Software | MS-DIAL |
| MS1 verified by standard | Yes | Data manipulation | Smoothing, Centroiding |
| MS2 verified by standard | Yes | Nomenclature for intact lipid molecule | Yes |
| Background check at MS1 | Yes | Nomenclature for fragment ions | No |
| Background check at MS2 | No | Further identification remarks | The reverse dot product similarity value, where the in silico spectrum is used for the library template, is used as the correlation coefficient value. The candidates are ranked by the reverse dot product score and the candidate with the highest similarity value is described as the representative C=C isomer candidate for the EAD-MS/MS spectrum. |

#### 46) SM[M+Na]+ / Lipid quantification

| | | | |
|---|---|---|---|
| Quantitative | No | Batch correction | No |
| Normalization to reference | No | Further quantification remarks | - |
