## Supplementary Note 2 for "MS-DIAL 5 multimodal mass spectrometry data mining unveils lipidome complexities"

### Separation Workflow

Created by https://lipidomicstandards.org, version v2.3.2

#### Overall study design

| | | | |
|---|---|---|---|
| Title of the study | Characterization of very long chain PUFA (VLC-PUFA) containing PC in the eye tissue of mice using EAD | | |
| Document creation date | 02/07/2024 | Corresponding Email | |
| Principle investigator | Hiroshi Tsugawa | Is the workflow targeted or untargeted? | Untargeted |
| Institution | Tokyo University of Agriculture and Technology | Clinical | No |

#### Lipid extraction

| | | | |
|---|---|---|---|
| Extraction method | 2-phase system | 2-phase system | Bligh&Dyer |
| pH adjustment | None | Were internal standards added prior extraction? | No |

#### Analytical platform

| | | | |
|---|---|---|---|
| Which solvents were used | (A) acetonitrile (ACN):MeOH:H2O (1:1:3, v/v/v) and (B) ACN:IPA (1:9, v/v). Both the solvents contained 10 nM ethylenediaminetetraacetic acid and 5 mM ammonium acetate. | Mass resolution for detected ion at MS1 | High resolution |
| Number of separation dimensions | One dimension | Resolution at m/z 200 at MS1 | 26316 |
| Separation type 1 | LC | Mass accuracy in ppm at MS1 | 1.13 |
| Separation mode 1 (liquid) | RP | Mass window for precursor ion isolation (in Da total isolation window) | 1 |
| Detector | Mass spectrometer | Mass resolution for detected ion at MS2 | High resolution |
| MS type | QTOF | Resolution at m/z 200 at MS2 | 28538 |
| MS vendor | SCIEX | Mass accuracy in ppm at MS2 | 0.76 |
| Ion source | ESI | Was/Were additional dimension/techniques used | No |
| MS Level | MS1, MS2 | | |

#### Quality control

| | | | |
|---|---|---|---|
| Blanks | Yes | Quality control | No |
| Type of Blanks | Extraction blank | | |

#### Method qualification and validation

| | | | |
|---|---|---|---|
| Method validation | Yes | Precison | Yes |
| Lipid recovery | No | Accuracy | Yes |
| Dynamic quantification range | Yes | Guidelines followed | None |
| Limit of quantitation (LOQ)/Limit of detection (LOD) | Yes | | |

#### Reporting

| | | | |
|---|---|---|---|
| Are reported raw data uploaded into repository? | Yes | Summary data | Identification data |
| Link to repository / ID to entry | http://prime.psc.riken.jp/menta.cgi/prime/drop_index, DM0054 | Raw data upload | Yes |
| Are metadata available? | Yes | Additional comments | The resolution and accuracy (ppm) for MS1 are those of m/z 132.9049 which were determined by the mass calibration in SCIEX OS. In addition, the resolution and accuracy for MS2 are those of m/z 185.1284, which were also determined by the mass calibration in SCIEX OS. |

### Sample Descriptions

#### Mouse eye ball / Mouse / Tissues (e.g., liver, heart, brain)

| | | | |
|---|---|---|---|
| Perfusion | No | Freeze-thaw cycles | 0 |
| Provided information | Time to separate plasma/serum (min), Time to freeze (min), Storage time (month), Freeze-thaw cycles | Additives | None |
| Temperature handling original sample | 4-8 °C | Were samples stored under inert gas? | No |
| Instant sample preparation | No | Additional preservation methods | No |
| Time to freeze (min) | 10 | Biobank samples | No |
| Snap freezing in liquid N2 | Yes | Sample homogenization | Yes |
| Storage temperature | -80 °C | Sample homogenization solvent | Methanol |
| Storage time (month) | 1 | | |

Fragments for identification:

| Fragment name |
|---|
| HG(PC,184) |
| HG + C3H5 |
| HG + C2H3O |
| NL of HG |
| NL of FA |

#### 9) PC[M+H]+ / Lipid quantification

| | | | |
|---|---|---|---|
| Quantitative | No | Batch correction | No |
| Normalization to reference | No | Further quantification remarks | - |

| Fragment name |
|---|
| -HG(PS,185) |
| HG+ |
| NL of FA |
| NL of HG and FA |
| HG + C3H6+ |
| HG + C2H4O+ |

#### 13) PS[M+H]+ / Lipid quantification

| | | | |
|---|---|---|---|
| Quantitative | No | Batch correction | No |
| Normalization to reference | No | Further quantification remarks | - |

#### 14) TG[M+NH4]+ / Lipid identification

#### 19) SM[M+H]+ / Lipid quantification

| | | | |
|---|---|---|---|
| Quantitative | No | Batch correction | No |
| Normalization to reference | No | Further quantification remarks | - |

#### 20) Hexosylceramide non-hydroxyfatty acid-dihydrosphingosine (HexCer_NDS)[M+H]+ / Lipid identification

| | | | |
|---|---|---|---|
| Lipid class | Hexosylceramide non-hydroxyfatty acid-dihydrosphingosine (HexCer_NDS) | Did you presume assumptions for identification? | No |
| Derivatization | - | Check isomer overlap | No |
| MS Level for identification | MS1, MS2 | RT verified by standard | No |
| Identification level | Double bond position | Separation of isobaric/isomeric interferece confirmed | No |
| Polarity mode | Positive | Model for separation prediction | No |
| Type of positive (precursor)ion | [M+H]+ | Additional dimension/techniques | EAD |
| Fragments for identification | | How was/were the additional dimension(s) used? | To determine doublebond positions |

| Fragment name |
|---|
| Neutral loss of H2O |
| Neutral loss of hexose |
| Neutral loss of hexose and H2O |
| Neutral loss of hexose and 2H2O |
| Fatty acyl amide +C2H5O + hexose fragment |
| Fatty acyl amide +C2H3O+ fragment |
| Fatty acyl amide +C2H2+ fragment |
| Fatty acyl amide+ fragment |
| Sphingosine -CH4O2 fragment |
| Sphingosine -2H2O fragment |
| Sphingosine -H2O fragment |
| Hexose + C2H5N+ |

| | | | |
|---|---|---|---|
| Isotope correction at MS1 | No | Was a model used to predict lipid molecule separation? | No |
| Isotope correction at MS2 | No | Lipid Identification Software | MS-DIAL |
| MS1 verified by standard | Yes | Data manipulation | Smoothing, Centroiding |
| MS2 verified by standard | Yes | Nomenclature for intact lipid molecule | Yes |
| Background check at MS1 | Yes | Nomenclature for fragment ions | No |
| Background check at MS2 | No | Further identification remarks | The reverse dot product similarity value, where the in silico spectrum is used for the library template, is used as the correlation coefficient value. The candidates are ranked by the reverse dot product score and the candidate with the highest similarity value is described as the representative C=C isomer candidate for the EAD-MS/MS spectrum. |

#### 20) Hexosylceramide non-hydroxyfatty acid-dihydrosphingosine (HexCer_NDS)[M+H]+ / Lipid quantification

| | | | |
|---|---|---|---|
| Quantitative | No | Batch correction | No |
| Normalization to reference | No | Further quantification remarks | - |

#### 21) DG[M+NH4]+ / Lipid identification

| | | | |
|---|---|---|---|
| Lipid class | DG | Background check at MS2 | No |
| Derivatization | - | Did you presume assumptions for identification? | No |
| MS Level for identification | MS1, MS2 | Check isomer overlap | No |
| Identification level | Molecular species level | RT verified by standard | No |
| Polarity mode | Positive | Separation of isobaric/isomeric interferece confirmed | No |
| Type of positive (precursor)ion | [M+NH4]+ | Model for separation prediction | No |
| Fragments for identification | | Additional dimension/techniques | - |

| Fragment name |
|---|
| Dehydro-monoacyl glycerols |
| Neutral loss of H2O |

| | | | |
|---|---|---|---|
| Isotope correction at MS1 | No | Lipid Identification Software | MS-DIAL |
| Isotope correction at MS2 | No | Data manipulation | Smoothing, Centroiding |
| MS1 verified by standard | Yes | Nomenclature for intact lipid molecule | Yes |
| MS2 verified by standard | Yes | Nomenclature for fragment ions | No |
| Background check at MS1 | Yes | Further identification remarks | - |

#### 21) DG[M+NH4]+ / Lipid quantification

| | | | |
|---|---|---|---|
| Quantitative | No | Batch correction | No |
| Normalization to reference | No | Further quantification remarks | - |

#### 22) Acylcarnitine (CAR)[M+H]+ / Lipid identification

| | | | |
|---|---|---|---|
| Lipid class | Acylcarnitine (CAR) | Background check at MS2 | No |
| Derivatization | - | Did you presume assumptions for identification? | No |
| MS Level for identification | MS1, MS2 | Check isomer overlap | No |
| Identification level | Molecular species level | RT verified by standard | No |
| Polarity mode | Positive | Separation of isobaric/isomeric interferece confirmed | No |
| Type of positive (precursor)ion | [M+H]+ | Model for separation prediction | No |
| Fragments for identification | | Additional dimension/techniques | - |

| Fragment name |
|---|
| Characteristic fragment (C4H5O2+) |

| | | | |
|---|---|---|---|
| Isotope correction at MS1 | No | Lipid Identification Software | MS-DIAL |
| Isotope correction at MS2 | No | Data manipulation | Smoothing, Centroiding |
| MS1 verified by standard | No | Nomenclature for intact lipid molecule | Yes |
| MS2 verified by standard | No | Nomenclature for fragment ions | No |
| Background check at MS1 | Yes | Further identification remarks | - |

#### 22) Acylcarnitine (CAR)[M+H]+ / Lipid quantification

| | | | |
|---|---|---|---|
| Quantitative | No | Batch correction | No |
| Normalization to reference | No | Further quantification remarks | - |

#### 23) BMP[M+NH4]+ / Lipid identification

| | | | |
|---|---|---|---|
| Lipid class | BMP | Background check at MS2 | No |
| Derivatization | - | Did you presume assumptions for identification? | No |
| MS Level for identification | MS1, MS2 | Check isomer overlap | No |
| Identification level | Molecular species level | RT verified by standard | No |
| Polarity mode | Positive | Separation of isobaric/isomeric interferece confirmed | No |
| Type of positive (precursor)ion | [M+NH4]+ | Model for separation prediction | No |
| Fragments for identification | | Additional dimension/techniques | - |

| Fragment name |
|---|
| Dehydro-monoacyl glycerols |
| Neutral loss of glycerophosphate |

| | | | |
|---|---|---|---|
| Isotope correction at MS1 | No | Lipid Identification Software | MS-DIAL |
| Isotope correction at MS2 | No | Data manipulation | Smoothing, Centroiding |
| MS1 verified by standard | No | Nomenclature for intact lipid molecule | Yes |
| MS2 verified by standard | No | Nomenclature for fragment ions | No |
| Background check at MS1 | Yes | Further identification remarks | - |

#### 23) BMP[M+NH4]+ / Lipid quantification

| | | | |
|---|---|---|---|
| Quantitative | No | Batch correction | No |
| Normalization to reference | No | Further quantification remarks | - |

#### 24) CE[M+NH4]+ / Lipid identification

| | | | |
|---|---|---|---|
| Lipid class | CE | Background check at MS2 | No |
| Derivatization | - | Did you presume assumptions for identification? | No |
| MS Level for identification | MS1, MS2 | Check isomer overlap | No |
| Identification level | Molecular species level | RT verified by standard | No |
| Polarity mode | Positive | Separation of isobaric/isomeric interferece confirmed | No |
| Type of positive (precursor)ion | [M+NH4]+ | Model for separation prediction | No |
| Fragments for identification | | Additional dimension/techniques | - |

| Fragment name |
|---|
| Neutral loss of fatty acyl |

| | | | |
|---|---|---|---|
| Isotope correction at MS1 | No | Lipid Identification Software | MS-DIAL |
| Isotope correction at MS2 | No | Data manipulation | Smoothing, Centroiding |
| MS1 verified by standard | Yes | Nomenclature for intact lipid molecule | Yes |
| MS2 verified by standard | Yes | Nomenclature for fragment ions | No |
| Background check at MS1 | Yes | Further identification remarks | - |

#### 24) CE[M+NH4]+ / Lipid quantification

| | | | |
|---|---|---|---|
| Quantitative | No | Batch correction | No |
| Normalization to reference | No | Further quantification remarks | - |

#### 25) Coenzyme Q (CoQ)[M+H]+ / Lipid identification

| | | | |
|---|---|---|---|
| Lipid class | Coenzyme Q (CoQ) | Background check at MS2 | No |
| Derivatization | - | Did you presume assumptions for identification? | No |
| MS Level for identification | MS1, MS2 | Check isomer overlap | No |
| Identification level | Molecular species level | RT verified by standard | No |
| Polarity mode | Positive | Separation of isobaric/isomeric interferece confirmed | No |
| Type of positive (precursor)ion | [M+H]+ | Model for separation prediction | No |
| Fragments for identification | | Additional dimension/techniques | - |

| Fragment name |
|---|
| Characteristic fragment (C10H13O4+) |

| | | | |
|---|---|---|---|
| Isotope correction at MS1 | No | Lipid Identification Software | MS-DIAL |
| Isotope correction at MS2 | No | Data manipulation | Smoothing, Centroiding |
| MS1 verified by standard | No | Nomenclature for intact lipid molecule | Yes |
| MS2 verified by standard | No | Nomenclature for fragment ions | No |
| Background check at MS1 | Yes | Further identification remarks | - |

#### 25) Coenzyme Q (CoQ)[M+H]+ / Lipid quantification

| | | | |
|---|---|---|---|
| Quantitative | No | Batch correction | No |
| Normalization to reference | No | Further quantification remarks | - |

#### 26) Hex2Cer[M+H]+ / Lipid identification

| | | | |
|---|---|---|---|
| Lipid class | Hex2Cer | Background check at MS2 | No |
| Derivatization | - | Did you presume assumptions for identification? | No |
| MS Level for identification | MS1, MS2 | Check isomer overlap | No |
| Identification level | Molecular species level | RT verified by standard | No |
| Polarity mode | Positive | Separation of isobaric/isomeric interferece confirmed | No |
| Type of positive (precursor)ion | [M+H]+ | Model for separation prediction | No |
| Fragments for identification | | Additional dimension/techniques | - |

| Fragment name |
|---|
| Neutral loss of hexose |
| Neutral loss of 2hexose |
| Sphingosine -H2O fragment |
| Sphingosine -2H2O fragment |
| Sphingosine -CH4O2 fragment |

| | | | |
|---|---|---|---|
| Isotope correction at MS1 | No | Lipid Identification Software | MS-DIAL |
| Isotope correction at MS2 | No | Data manipulation | Smoothing, Centroiding |
| MS1 verified by standard | No | Nomenclature for intact lipid molecule | Yes |
| MS2 verified by standard | No | Nomenclature for fragment ions | No |
| Background check at MS1 | Yes | Further identification remarks | - |

#### 26) Hex2Cer[M+H]+ / Lipid quantification

| | | | |
|---|---|---|---|
| Quantitative | No | Batch correction | No |
| Normalization to reference | No | Further quantification remarks | - |

#### 27) LPE O[M+H]+ / Lipid identification

| | | | |
|---|---|---|---|
| Lipid class | LPE O | Background check at MS2 | No |
| Derivatization | - | Did you presume assumptions for identification? | No |
| MS Level for identification | MS1, MS2 | Check isomer overlap | No |
| Identification level | Molecular species level | RT verified by standard | No |
| Polarity mode | Positive | Separation of isobaric/isomeric interferece confirmed | No |
| Type of positive (precursor)ion | [M+H]+ | Model for separation prediction | No |
| Fragments for identification | | Additional dimension/techniques | - |

| Fragment name |
|---|
| Neutral loss of C3H8NO4P |
| Neutral loss of C3H10NO5P |

| | | | |
|---|---|---|---|
| Isotope correction at MS1 | No | Lipid Identification Software | MS-DIAL |
| Isotope correction at MS2 | No | Data manipulation | Smoothing, Centroiding |
| MS1 verified by standard | No | Nomenclature for intact lipid molecule | Yes |
| MS2 verified by standard | No | Nomenclature for fragment ions | No |
| Background check at MS1 | Yes | Further identification remarks | - |

#### 27) LPE O[M+H]+ / Lipid quantification

| | | | |
|---|---|---|---|
| Quantitative | No | Batch correction | No |
| Normalization to reference | No | Further quantification remarks | - |

#### 28) Ether-linked triacylglycerol (EtherTG)[M+NH4]+ / Lipid identification

| | | | |
|---|---|---|---|
| Lipid class | Ether-linked triacylglycerol (EtherTG) | Background check at MS2 | No |
| Derivatization | - | Did you presume assumptions for identification? | No |
| MS Level for identification | MS1, MS2 | Check isomer overlap | No |
| Identification level | Molecular species level | RT verified by standard | No |
| Polarity mode | Positive | Separation of isobaric/isomeric interferece confirmed | No |
| Type of positive (precursor)ion | [M+NH4]+ | Model for separation prediction | No |
| Fragments for identification | | Additional dimension/techniques | - |

| Fragment name |
|---|
| Neutral loss of fatty acyl and H2O |
| Neutral loss of alkyl ether |

| | | | |
|---|---|---|---|
| Isotope correction at MS1 | No | Lipid Identification Software | MS-DIAL |
| Isotope correction at MS2 | No | Data manipulation | Smoothing, Centroiding |
| MS1 verified by standard | No | Nomenclature for intact lipid molecule | Yes |
| MS2 verified by standard | No | Nomenclature for fragment ions | No |
| Background check at MS1 | Yes | Further identification remarks | - |

#### 28) Ether-linked triacylglycerol (EtherTG)[M+NH4]+ / Lipid quantification

| | | | |
|---|---|---|---|
| Quantitative | No | Batch correction | No |
| Normalization to reference | No | Further quantification remarks | - |

#### 29) Hexosylceramide hydroxyfatty acid-sphingosine (HexCer_HS)[M+H]+ / Lipid identification

| | | | |
|---|---|---|---|
| Lipid class | Hexosylceramide hydroxyfatty acid-sphingosine (HexCer_HS) | Background check at MS2 | No |
| Derivatization | - | Did you presume assumptions for identification? | No |
| MS Level for identification | MS1, MS2 | Check isomer overlap | No |
| Identification level | Molecular species level | RT verified by standard | No |
| Polarity mode | Positive | Separation of isobaric/isomeric interferece confirmed | No |
| Type of positive (precursor)ion | [M+H]+ | Model for separation prediction | No |
| Fragments for identification | | Additional dimension/techniques | - |

| Fragment name |
|---|
| Neutral loss of hexose |
| Neutral loss of hexose and H2O |
| Sphingosine -H2O fragment |
| Sphingosine -2H2O fragment |
| Sphingosine -CH4O2 fragment |

| | | | |
|---|---|---|---|
| Isotope correction at MS1 | No | Lipid Identification Software | MS-DIAL |
| Isotope correction at MS2 | No | Data manipulation | Smoothing, Centroiding |
| MS1 verified by standard | No | Nomenclature for intact lipid molecule | Yes |
| MS2 verified by standard | No | Nomenclature for fragment ions | No |
| Background check at MS1 | Yes | Further identification remarks | - |

#### 29) Hexosylceramide hydroxyfatty acid-sphingosine (HexCer_HS)[M+H]+ / Lipid quantification

| | | | |
|---|---|---|---|
| Quantitative | No | Batch correction | No |
| Normalization to reference | No | Further quantification remarks | - |

#### 30) LPC[M+H]+ / Lipid identification

| | | | |
|---|---|---|---|
| Lipid class | LPC | Background check at MS2 | No |
| Derivatization | - | Did you presume assumptions for identification? | No |
| MS Level for identification | MS1, MS2 | Check isomer overlap | No |
| Identification level | Molecular species level | RT verified by standard | No |
| Polarity mode | Positive | Separation of isobaric/isomeric interferece confirmed | No |
| Type of positive (precursor)ion | [M+H]+ | Model for separation prediction | No |
| Fragments for identification | | Additional dimension/techniques | - |

| Fragment name |
|---|
| Characteristic fragment (C5H15NO4P+) |

| | | | |
|---|---|---|---|
| Isotope correction at MS1 | No | Lipid Identification Software | MS-DIAL |
| Isotope correction at MS2 | No | Data manipulation | Smoothing, Centroiding |
| MS1 verified by standard | Yes | Nomenclature for intact lipid molecule | Yes |
| MS2 verified by standard | Yes | Nomenclature for fragment ions | No |
| Background check at MS1 | Yes | Further identification remarks | - |

#### 30) LPC[M+H]+ / Lipid quantification

| | | | |
|---|---|---|---|
| Quantitative | No | Batch correction | No |
| Normalization to reference | No | Further quantification remarks | - |

#### 31) LPE[M+H]+ / Lipid identification

| | | | |
|---|---|---|---|
| Lipid class | LPE | Background check at MS2 | No |
| Derivatization | - | Did you presume assumptions for identification? | No |
| MS Level for identification | MS1, MS2 | Check isomer overlap | No |
| Identification level | Molecular species level | RT verified by standard | No |
| Polarity mode | Positive | Separation of isobaric/isomeric interferece confirmed | No |
| Type of positive (precursor)ion | [M+H]+ | Model for separation prediction | No |
| Fragments for identification | | Additional dimension/techniques | - |

| Fragment name |
|---|
| Neutral loss of C2H8NO4P |

| | | | |
|---|---|---|---|
| Isotope correction at MS1 | No | Lipid Identification Software | MS-DIAL |
| Isotope correction at MS2 | No | Data manipulation | Smoothing, Centroiding |
| MS1 verified by standard | Yes | Nomenclature for intact lipid molecule | Yes |
| MS2 verified by standard | Yes | Nomenclature for fragment ions | No |
| Background check at MS1 | Yes | Further identification remarks | - |

#### 31) LPE[M+H]+ / Lipid quantification

| | | | |
|---|---|---|---|
| Quantitative | No | Batch correction | No |
| Normalization to reference | No | Further quantification remarks | - |

#### 32) MG[M+NH4]+ / Lipid identification

| | | | |
|---|---|---|---|
| Lipid class | MG | Background check at MS2 | No |
| Derivatization | - | Did you presume assumptions for identification? | No |
| MS Level for identification | MS1, MS2 | Check isomer overlap | No |
| Identification level | Molecular species level | RT verified by standard | No |
| Polarity mode | Positive | Separation of isobaric/isomeric interferece confirmed | No |
| Type of positive (precursor)ion | [M+NH4]+ | Model for separation prediction | No |
| Fragments for identification | | Additional dimension/techniques | - |

| Fragment name |
|---|
| Neutral loss of H2O |

| | | | |
|---|---|---|---|
| Isotope correction at MS1 | No | Lipid Identification Software | MS-DIAL |
| Isotope correction at MS2 | No | Data manipulation | Smoothing, Centroiding |
| MS1 verified by standard | Yes | Nomenclature for intact lipid molecule | Yes |
| MS2 verified by standard | Yes | Nomenclature for fragment ions | No |
| Background check at MS1 | Yes | Further identification remarks | - |

#### 32) MG[M+NH4]+ / Lipid quantification

| | | | |
|---|---|---|---|
| Quantitative | No | Batch correction | No |
| Normalization to reference | No | Further quantification remarks | - |

#### 33) MGDG[M+NH4]+ / Lipid identification

| | | | |
|---|---|---|---|
| Lipid class | MGDG | Background check at MS2 | No |
| Derivatization | - | Did you presume assumptions for identification? | No |
| MS Level for identification | MS1, MS2 | Check isomer overlap | No |
| Identification level | Molecular species level | RT verified by standard | No |
| Polarity mode | Positive | Separation of isobaric/isomeric interferece confirmed | No |
| Type of positive (precursor)ion | [M+NH4]+ | Model for separation prediction | No |
| Fragments for identification | | Additional dimension/techniques | - |

| Fragment name |
|---|
| -HG(Hex,180) |
| -HG(Hex,180) -H2O and -SN1 acyl chain |
| -HG(Hex,180) -H2O and -SN2 acyl chain |

| | | | |
|---|---|---|---|
| Isotope correction at MS1 | No | Lipid Identification Software | MS-DIAL |
| Isotope correction at MS2 | No | Data manipulation | Smoothing, Centroiding |
| MS1 verified by standard | No | Nomenclature for intact lipid molecule | Yes |
| MS2 verified by standard | No | Nomenclature for fragment ions | No |
| Background check at MS1 | Yes | Further identification remarks | - |

#### 33) MGDG[M+NH4]+ / Lipid quantification

| | | | |
|---|---|---|---|
| Quantitative | No | Batch correction | No |
| Normalization to reference | No | Further quantification remarks | - |

#### 34) PC[M+H]+ / Lipid identification

| | | | |
|---|---|---|---|
| Lipid class | PC | Background check at MS2 | No |
| Derivatization | - | Did you presume assumptions for identification? | No |
| MS Level for identification | MS1, MS2 | Check isomer overlap | No |
| Identification level | Molecular species level | RT verified by standard | No |
| Polarity mode | Positive | Separation of isobaric/isomeric interferece confirmed | No |
| Type of positive (precursor)ion | [M+H]+ | Model for separation prediction | No |
| Fragments for identification | | Additional dimension/techniques | - |

| Fragment name |
|---|
| HG(PC,184) |
| NL of SN1 fatty acyl |
| NL of SN2 fatty acyl |
| NL of SN1 fatty acyl and H2O |
| NL of SN2 fatty acyl and H2O |

| | | | |
|---|---|---|---|
| Isotope correction at MS1 | No | Lipid Identification Software | MS-DIAL |
| Isotope correction at MS2 | No | Data manipulation | Smoothing, Centroiding |
| MS1 verified by standard | Yes | Nomenclature for intact lipid molecule | Yes |
| MS2 verified by standard | Yes | Nomenclature for fragment ions | No |
| Background check at MS1 | Yes | Further identification remarks | - |

#### 34) PC[M+H]+ / Lipid quantification

| | | | |
|---|---|---|---|
| Quantitative | No | Batch correction | No |
| Normalization to reference | No | Further quantification remarks | - |

#### 35) PE[M+H]+ / Lipid identification

| | | | |
|---|---|---|---|
| Lipid class | PE | Background check at MS2 | No |
| Derivatization | - | Did you presume assumptions for identification? | No |
| MS Level for identification | MS1, MS2 | Check isomer overlap | No |
| Identification level | Molecular species level | RT verified by standard | No |
| Polarity mode | Positive | Separation of isobaric/isomeric interferece confirmed | No |
| Type of positive (precursor)ion | [M+H]+ | Model for separation prediction | No |
| Fragments for identification | | Additional dimension/techniques | - |

| Fragment name |
|---|
| -HG(PE,141) |
| Fatty acyl fragment |

| | | | |
|---|---|---|---|
| Isotope correction at MS1 | No | Lipid Identification Software | MS-DIAL |
| Isotope correction at MS2 | No | Data manipulation | Smoothing, Centroiding |
| MS1 verified by standard | Yes | Nomenclature for intact lipid molecule | Yes |
| MS2 verified by standard | Yes | Nomenclature for fragment ions | No |
| Background check at MS1 | Yes | Further identification remarks | - |

#### 35) PE[M+H]+ / Lipid quantification

| | | | |
|---|---|---|---|
| Quantitative | No | Batch correction | No |
| Normalization to reference | No | Further quantification remarks | - |

#### 36) PG[M+NH4]+ / Lipid identification

| | | | |
|---|---|---|---|
| Lipid class | PG | Background check at MS2 | No |
| Derivatization | - | Did you presume assumptions for identification? | No |
| MS Level for identification | MS1, MS2 | Check isomer overlap | No |
| Identification level | Molecular species level | RT verified by standard | No |
| Polarity mode | Positive | Separation of isobaric/isomeric interferece confirmed | No |
| Type of positive (precursor)ion | [M+NH4]+ | Model for separation prediction | No |
| Fragments for identification | | Additional dimension/techniques | - |

| Fragment name |
|---|
| -HG(PG,172) |
| Fatty acyl fragment |

| | | | |
|---|---|---|---|
| Isotope correction at MS1 | No | Lipid Identification Software | MS-DIAL |
| Isotope correction at MS2 | No | Data manipulation | Smoothing, Centroiding |
| MS1 verified by standard | Yes | Nomenclature for intact lipid molecule | Yes |
| MS2 verified by standard | Yes | Nomenclature for fragment ions | No |
| Background check at MS1 | Yes | Further identification remarks | - |

#### 36) PG[M+NH4]+ / Lipid quantification

| | | | |
|---|---|---|---|
| Quantitative | No | Batch correction | No |
| Normalization to reference | No | Further quantification remarks | - |

#### 37) PI[M+NH4]+ / Lipid identification

| | | | |
|---|---|---|---|
| Lipid class | PI | Background check at MS2 | No |
| Derivatization | - | Did you presume assumptions for identification? | No |
| MS Level for identification | MS1, MS2 | Check isomer overlap | No |
| Identification level | Molecular species level | RT verified by standard | No |
| Polarity mode | Positive | Separation of isobaric/isomeric interferece confirmed | No |
| Type of positive (precursor)ion | [M+NH4]+ | Model for separation prediction | No |
| Fragments for identification | | Additional dimension/techniques | - |

| Fragment name |
|---|
| -HG(PI,260) |
| NL of SN1 acyl chain |
| NL of SN1 acyl chain and H2O |
| NL of SN2 acyl chain |
| NL of SN2 acyl chain and H2O |

| | | | |
|---|---|---|---|
| Isotope correction at MS1 | No | Lipid Identification Software | MS-DIAL |
| Isotope correction at MS2 | No | Data manipulation | Smoothing, Centroiding |
| MS1 verified by standard | Yes | Nomenclature for intact lipid molecule | Yes |
| MS2 verified by standard | Yes | Nomenclature for fragment ions | No |
| Background check at MS1 | Yes | Further identification remarks | - |

#### 37) PI[M+NH4]+ / Lipid quantification

| | | | |
|---|---|---|---|
| Quantitative | No | Batch correction | No |
| Normalization to reference | No | Further quantification remarks | - |

#### 38) PS[M+H]+ / Lipid identification

| | | | |
|---|---|---|---|
| Lipid class | PS | Background check at MS2 | No |
| Derivatization | - | Did you presume assumptions for identification? | No |
| MS Level for identification | MS1, MS2 | Check isomer overlap | No |
| Identification level | Molecular species level | RT verified by standard | No |
| Polarity mode | Positive | Separation of isobaric/isomeric interferece confirmed | No |
| Type of positive (precursor)ion | [M+H]+ | Model for separation prediction | No |
| Fragments for identification | | Additional dimension/techniques | - |

| Fragment name |
|---|
| -HG(PS,185) |
| NL of fatty acyl chain |
| NL of fatty acyl chain and H2O |

| | | | |
|---|---|---|---|
| Isotope correction at MS1 | No | Lipid Identification Software | MS-DIAL |
| Isotope correction at MS2 | No | Data manipulation | Smoothing, Centroiding |
| MS1 verified by standard | Yes | Nomenclature for intact lipid molecule | Yes |
| MS2 verified by standard | Yes | Nomenclature for fragment ions | No |
| Background check at MS1 | Yes | Further identification remarks | - |

#### 38) PS[M+H]+ / Lipid quantification

| | | | |
|---|---|---|---|
| Quantitative | No | Batch correction | No |
| Normalization to reference | No | Further quantification remarks | - |

#### 39) SPB[M+H]+ / Lipid identification

| | | | |
|---|---|---|---|
| Lipid class | SPB | Background check at MS2 | No |
| Derivatization | - | Did you presume assumptions for identification? | No |
| MS Level for identification | MS1, MS2 | Check isomer overlap | No |
| Identification level | Molecular species level | RT verified by standard | No |
| Polarity mode | Positive | Separation of isobaric/isomeric interferece confirmed | No |
| Type of positive (precursor)ion | [M+H]+ | Model for separation prediction | No |
| Fragments for identification | | Additional dimension/techniques | - |

| Fragment name |
|---|
| Neutral loss of H2O |
| Neutral loss of 2H2O |
| Neutral loss of CH4O2 |

| | | | |
|---|---|---|---|
| Isotope correction at MS1 | No | Lipid Identification Software | MS-DIAL |
| Isotope correction at MS2 | No | Data manipulation | Smoothing, Centroiding |
| MS1 verified by standard | No | Nomenclature for intact lipid molecule | Yes |
| MS2 verified by standard | No | Nomenclature for fragment ions | No |
| Background check at MS1 | Yes | Further identification remarks | - |

#### 39) SPB[M+H]+ / Lipid quantification

| | | | |
|---|---|---|---|
| Quantitative | No | Batch correction | No |
| Normalization to reference | No | Further quantification remarks | - |

#### 40) TG[M+NH4]+ / Lipid identification

| | | | |
|---|---|---|---|
| Lipid class | TG | Background check at MS2 | No |
| Derivatization | - | Did you presume assumptions for identification? | No |
| MS Level for identification | MS1, MS2 | Check isomer overlap | No |
| Identification level | Molecular species level | RT verified by standard | No |
| Polarity mode | Positive | Separation of isobaric/isomeric interferece confirmed | No |
| Type of positive (precursor)ion | [M+NH4]+ | Model for separation prediction | No |
| Fragments for identification | | Additional dimension/techniques | - |

| Fragment name |
|---|
| Neutral loss of acyl and H2O |

| | | | |
|---|---|---|---|
| Isotope correction at MS1 | No | Lipid Identification Software | MS-DIAL |
| Isotope correction at MS2 | No | Data manipulation | Smoothing, Centroiding |
| MS1 verified by standard | Yes | Nomenclature for intact lipid molecule | Yes |
| MS2 verified by standard | Yes | Nomenclature for fragment ions | No |
| Background check at MS1 | Yes | Further identification remarks | - |

#### 40) TG[M+NH4]+ / Lipid quantification

| | | | |
|---|---|---|---|
| Quantitative | No | Batch correction | No |
| Normalization to reference | No | Further quantification remarks | - |

#### 41) Triacylglycerol estolides (TG_EST)[M+NH4]+ / Lipid identification

| | | | |
|---|---|---|---|
| Lipid class | Triacylglycerol estolides (TG_EST) | Background check at MS2 | No |
| Derivatization | - | Did you presume assumptions for identification? | No |
| MS Level for identification | MS1, MS2 | Check isomer overlap | No |
| Identification level | Molecular species level | RT verified by standard | No |
| Polarity mode | Positive | Separation of isobaric/isomeric interferece confirmed | No |
| Type of positive (precursor)ion | [M+NH4]+ | Model for separation prediction | No |
| Fragments for identification | | Additional dimension/techniques | - |

| Fragment name |
|---|
| Neutral loss of acyl and H2O |
| Neutral loss of oxidized acyl and H2O |
| Neutral loss of FAHFA |

| | | | |
|---|---|---|---|
| Isotope correction at MS1 | No | Lipid Identification Software | MS-DIAL |
| Isotope correction at MS2 | No | Data manipulation | Smoothing, Centroiding |
| MS1 verified by standard | No | Nomenclature for intact lipid molecule | Yes |
| MS2 verified by standard | No | Nomenclature for fragment ions | No |
| Background check at MS1 | Yes | Further identification remarks | - |

#### 41) Triacylglycerol estolides (TG_EST)[M+NH4]+ / Lipid quantification

| | | | |
|---|---|---|---|
| Quantitative | No | Batch correction | No |
| Normalization to reference | No | Further quantification remarks | - |

#### 42) PE P[M+H]+ / Lipid identification

| | | | |
|---|---|---|---|
| Lipid class | PE P | Background check at MS2 | No |
| Derivatization | - | Did you presume assumptions for identification? | No |
| MS Level for identification | MS1, MS2 | Check isomer overlap | No |
| Identification level | Molecular species level | RT verified by standard | No |
| Polarity mode | Positive | Separation of isobaric/isomeric interferece confirmed | No |
| Type of positive (precursor)ion | [M+H]+ | Model for separation prediction | No |
| Fragments for identification | | Additional dimension/techniques | - |

| Fragment name |
|---|
| Neutral loss of C2H8NO4P |
| Alkyl ether +C2H8NO3P fragment |
| Dehydro-monoacyl glycerols |

| | | | |
|---|---|---|---|
| Isotope correction at MS1 | No | Lipid Identification Software | MS-DIAL |
| Isotope correction at MS2 | No | Data manipulation | Smoothing, Centroiding |
| MS1 verified by standard | No | Nomenclature for intact lipid molecule | Yes |
| MS2 verified by standard | No | Nomenclature for fragment ions | No |
| Background check at MS1 | Yes | Further identification remarks | - |

#### 42) PE P[M+H]+ / Lipid quantification

| | | | |
|---|---|---|---|
| Quantitative | No | Batch correction | No |
| Normalization to reference | No | Further quantification remarks | - |

#### 43) N-acyl ethanolamines (NAE)[M+H]+ / Lipid identification

| | | | |
|---|---|---|---|
| Lipid class | N-acyl ethanolamines (NAE) | Background check at MS2 | No |
| Derivatization | - | Did you presume assumptions for identification? | No |
| MS Level for identification | MS1, MS2 | Check isomer overlap | No |
| Identification level | Molecular species level | RT verified by standard | No |
| Polarity mode | Positive | Separation of isobaric/isomeric interferece confirmed | No |
| Type of positive (precursor)ion | [M+H]+ | Model for separation prediction | No |
| Fragments for identification | | Additional dimension/techniques | - |

| Fragment name |
|---|
| Neutral loss of 2H2O |

| | | | |
|---|---|---|---|
| Isotope correction at MS1 | No | Lipid Identification Software | MS-DIAL |
| Isotope correction at MS2 | No | Data manipulation | Smoothing, Centroiding |
| MS1 verified by standard | No | Nomenclature for intact lipid molecule | Yes |
| MS2 verified by standard | No | Nomenclature for fragment ions | No |
| Background check at MS1 | Yes | Further identification remarks | - |

#### 43) N-acyl ethanolamines (NAE)[M+H]+ / Lipid quantification

| | | | |
|---|---|---|---|
| Quantitative | No | Batch correction | No |
| Normalization to reference | No | Further quantification remarks | - |

#### 44) ST[M+NH4]+ / Lipid identification

| | | | |
|---|---|---|---|
| Lipid class | ST | Background check at MS2 | No |
| Derivatization | - | Did you presume assumptions for identification? | No |
| MS Level for identification | MS1, MS2 | Check isomer overlap | No |
| Identification level | Species level | RT verified by standard | No |
| Polarity mode | Positive | Separation of isobaric/isomeric interferece confirmed | No |
| Type of positive (precursor)ion | [M+NH4]+ | Model for separation prediction | No |
| Fragments for identification | | Additional dimension/techniques | - |

| Fragment name |
|---|
| precursor m/z |

| | | | |
|---|---|---|---|
| Isotope correction at MS1 | No | Lipid Identification Software | MS-DIAL |
| Isotope correction at MS2 | No | Data manipulation | Smoothing, Centroiding |
| MS1 verified by standard | No | Nomenclature for intact lipid molecule | Yes |
| MS2 verified by standard | No | Nomenclature for fragment ions | No |
| Background check at MS1 | Yes | Further identification remarks | - |

#### 44) ST[M+NH4]+ / Lipid quantification

| | | | |
|---|---|---|---|
| Quantitative | No | Batch correction | No |
| Normalization to reference | No | Further quantification remarks | - |

#### 45) PC O[M+H]+ / Lipid identification

| | | | |
|---|---|---|---|
| Lipid class | PC O | Background check at MS2 | No |
| Derivatization | - | Did you presume assumptions for identification? | No |
| MS Level for identification | MS1, MS2 | Check isomer overlap | No |
| Identification level | Molecular species level | RT verified by standard | No |
| Polarity mode | Positive | Separation of isobaric/isomeric interferece confirmed | No |
| Type of positive (precursor)ion | [M+H]+ | Model for separation prediction | No |
| Fragments for identification | | Additional dimension/techniques | - |

| Fragment name |
|---|
| HG(PC,184) |
| NL of fatty acyl chain |

| | | | |
|---|---|---|---|
| Isotope correction at MS1 | No | Lipid Identification Software | MS-DIAL |
| Isotope correction at MS2 | No | Data manipulation | Smoothing, Centroiding |
| MS1 verified by standard | No | Nomenclature for intact lipid molecule | Yes |
| MS2 verified by standard | No | Nomenclature for fragment ions | No |
| Background check at MS1 | Yes | Further identification remarks | - |

#### 45) PC O[M+H]+ / Lipid quantification

| | | | |
|---|---|---|---|
| Quantitative | No | Batch correction | No |
| Normalization to reference | No | Further quantification remarks | - |

#### 46) SM[M+H]+ / Lipid identification

| | | | |
|---|---|---|---|
| Lipid class | SM | Background check at MS2 | No |
| Derivatization | - | Did you presume assumptions for identification? | No |
| MS Level for identification | MS1, MS2 | Check isomer overlap | No |
| Identification level | Molecular species level | RT verified by standard | No |
| Polarity mode | Positive | Separation of isobaric/isomeric interferece confirmed | No |
| Type of positive (precursor)ion | [M+H]+ | Model for separation prediction | No |
| Fragments for identification | | Additional dimension/techniques | - |

| Fragment name |
|---|
| HG(PC,184) |
| Sphingosine fragment |

| | | | |
|---|---|---|---|
| Isotope correction at MS1 | No | Lipid Identification Software | MS-DIAL |
| Isotope correction at MS2 | No | Data manipulation | Smoothing, Centroiding |
| MS1 verified by standard | No | Nomenclature for intact lipid molecule | Yes |
| MS2 verified by standard | No | Nomenclature for fragment ions | No |
| Background check at MS1 | Yes | Further identification remarks | - |

#### 46) SM[M+H]+ / Lipid quantification

| | | | |
|---|---|---|---|
| Quantitative | No | Batch correction | No |
| Normalization to reference | No | Further quantification remarks | - |

#### 47) Ether-linked diacylglycerol (EtherDG)[M+NH4]+ / Lipid identification

| | | | |
|---|---|---|---|
| Lipid class | Ether-linked diacylglycerol (EtherDG) | Background check at MS2 | No |
| Derivatization | - | Did you presume assumptions for identification? | No |
| MS Level for identification | MS1, MS2 | Check isomer overlap | No |
| Identification level | Molecular species level | RT verified by standard | No |
| Polarity mode | Positive | Separation of isobaric/isomeric interferece confirmed | No |
| Type of positive (precursor)ion | [M+NH4]+ | Model for separation prediction | No |
| Fragments for identification | | Additional dimension/techniques | - |

| Fragment name |
|---|
| NL of fatty acyl chain |

| | | | |
|---|---|---|---|
| Isotope correction at MS1 | No | Lipid Identification Software | MS-DIAL |
| Isotope correction at MS2 | No | Data manipulation | Smoothing, Centroiding |
| MS1 verified by standard | No | Nomenclature for intact lipid molecule | Yes |
| MS2 verified by standard | No | Nomenclature for fragment ions | No |
| Background check at MS1 | Yes | Further identification remarks | - |

#### 47) Ether-linked diacylglycerol (EtherDG)[M+NH4]+ / Lipid quantification

| | | | |
|---|---|---|---|
| Quantitative | No | Batch correction | No |
| Normalization to reference | No | Further quantification remarks | - |

#### 48) Ceramide hydroxy fatty acid-sphingosine (Cer_HS)[M+H]+ / Lipid identification

| | | | |
|---|---|---|---|
| Lipid class | Ceramide hydroxy fatty acid-sphingosine (Cer_HS) | Background check at MS2 | No |
| Derivatization | - | Did you presume assumptions for identification? | No |
| MS Level for identification | MS1, MS2 | Check isomer overlap | No |
| Identification level | Molecular species level | RT verified by standard | No |
| Polarity mode | Positive | Separation of isobaric/isomeric interferece confirmed | No |
| Type of positive (precursor)ion | [M+H]+ | Model for separation prediction | No |
| Fragments for identification | | Additional dimension/techniques | - |

| Fragment name |
|---|
| NL of H2O |
| NL of hydroxy fatty acyl chain |
| NL of hydroxy fatty acyl chain and H2O |
| NL of hydroxy fatty acyl chain and CH2O |

| | | | |
|---|---|---|---|
| Isotope correction at MS1 | No | Lipid Identification Software | MS-DIAL |
| Isotope correction at MS2 | No | Data manipulation | - |
| MS1 verified by standard | No | Nomenclature for intact lipid molecule | Yes |
| MS2 verified by standard | No | Nomenclature for fragment ions | No |
| Background check at MS1 | Yes | Further identification remarks | - |

#### 48) Ceramide hydroxy fatty acid-sphingosine (Cer_HS)[M+H]+ / Lipid quantification

| | | | |
|---|---|---|---|
| Quantitative | No | Batch correction | No |
| Normalization to reference | No | Further quantification remarks | - |
