## Supplementary Note 1 for "MS-DIAL 5 multimodal mass spectrometry data mining unveils lipidome complexities"

### Separation Workflow

Created by https://lipidomicstandards.org, version v2.0.0

#### Overall study design

| | | | |
|---|---|---|---|
| Title of the study | HeLa lipid profiling for HeLa cells with the supplementation of VLC-PUFA (FA 32:6) | | |
| Document creation date | 02/07/2024 | Corresponding Email | |
| Principle investigator | Hiroshi Tsugawa | Is the workflow targeted or untargeted? | Untargeted |
| Institution | Tokyo University of Agriculture and Technology | Clinical | No |

#### Lipid extraction

| | | | |
|---|---|---|---|
| Extraction method | 2-phase system | 2-phase system | Bligh&Dyer |
| pH adjustment | None | Were internal standards added prior extraction? | No |

#### Analytical platform

| | | | |
|---|---|---|---|
| Number of separation dimensions | One dimension | Ion source | ESI |
| Separation Type 1 | LC | MS Level | MS1, MS2 |
| Separation Mode 1 | RP | Mass resolution for detected ion at MS1 | High resolution |
| Separation window (1) for lipid analyte selection (±) in minutes | 1.5 | Resolution at m/z 200 at MS1 | 25071 |
| RT verified by standard | Yes | Mass accuracy in ppm at MS1 | 0.00549 |
| CCS verified by standard | No | Mass window for precursor ion isolation (in Da total isolation window) | 1 |
| Separation of isobaric/isomeric interferece confirmed | No | Mass resolution for detected ion at MS2 | High resolution |
| Model for separation prediction | No | Resolution at m/z 200 at MS2 | 27858 |
| MS type | QTOF | Mass accuracy in ppm at MS2 | 0.09463 |
| MS vendor | SCIEX | Was/Were additional dimension/techniques used | No |

#### Quality control

| | | | |
|---|---|---|---|
| Blanks | Yes | Quality control | No |
| Type of Blanks | Extraction blank, Injection blank | | |

#### Method qualification and validation

| | | | |
|---|---|---|---|
| Method validation | Yes | Precison | Yes |
| Lipid recovery | Yes | Accuracy | Yes |
| Dynamic quantification range | Yes | Guidelines followed | None |
| Limit of quantitation (LOQ)/Limit of detection (LOD) | Yes | | |

#### Reporting

| | | | |
|---|---|---|---|
| Are reported raw data uploaded into repository? | Yes | Raw data upload | Yes |
| Are metadata available? | Yes | Additional comments | The raw data is available under the index of DM0054 at http://prime.psc.riken.jp/menta.cgi/prime/drop[illegible]dex. The resolution and accuracy (ppm) for MS1 are those of m/z 132.9049 which were determined by the mass calibration in SCIEX OS. In addition, the resolution and accuracy for MS2 are those of m/z 185.1284, which were also determined by the mass calibration in SCIEX OS. |
| Summary data | Quantification and identification data | | |

### Sample Descriptions

#### HeLa cells / Human / Cells

| | | | |
|---|---|---|---|
| Provided information | Time to freeze (min), Storage time (month), Freeze-thaw cycles | Storage time (month) | 1 |
| Temperature handling original sample | 4-8 °C | Freeze-thaw cycles | 0 |
| Instant sample preparation | Yes | Additives | None |
| Time to freeze (min) | 10 | Were samples stored under inert gas? | No |
| Snap freezing in liquid N2 | No | Additional preservation methods | No |
| Storage temperature | -80 °C | Biobank samples | No |

### Lipid Class Descriptions

#### 1) FA[M-H]- / Lipid identification

| | | | |
|---|---|---|---|
| Lipid class | FA | Check isomer overlap | No |
| Derivatization | - | RT verified by standard | Yes |
| MS Level for identification | MS1 | Separation of isobaric/isomeric interferece confirmed | Yes |
| Identification level | Species level | Model for separation prediction | Yes |
| Polarity mode | Negative | Additional dimension/techniques | - |
| Type of negative (precursor)ion | [M-H]- | Lipid Identification Software | MS-DIAL |
| Isotope correction at MS1 | No | Data manipulation | Smoothing, Centroiding |
| MS1 verified by standard | Yes | Nomenclature for intact lipid molecule | Yes |
| Background check at MS1 | Yes | Further identification remarks | - |
| Did you presume assumptions for identification? | No | | |

#### 1) FA[M-H]- / Lipid quantification

| | | | |
|---|---|---|---|
| Quantitative | Yes | Limit of quantification | No |
| MS Level for quantification | MS1 | Normalization to reference | No |
| Internal lipid standard(s) MS1 | | Lipid Quantification Software | MS-DIAL |

| Internal standard | Endogenous subclass |
|---|---|
| FA 18:0(d3) | FA subclass |

| | | | |
|---|---|---|---|
| Type of quantification | Internal standard amount | Batch correction | No |
| Response correction | No | Further quantification remarks | - |
| Type I isotope correction | No | | |

#### 2) DG[M+NH4]+ / Lipid identification

| | | | |
|---|---|---|---|
| Lipid class | DG | Background check at MS2 | No |
| Derivatization | - | Did you presume assumptions for identification? | No |
| MS Level for identification | MS1, MS2 | Check isomer overlap | No |
| Identification level | Molecular species level | RT verified by standard | Yes |
| Polarity mode | Positive | Separation of isobaric/isomeric interferece confirmed | Yes |
| Type of positive (precursor)ion | [M+NH4]+ | Model for separation prediction | Yes |
| Fragments for identification | | Additional dimension/techniques | - |

| Fragment name |
|---|
| Dehydro-monoacyl glycerols |
| Neutral loss of H2O |

| | | | |
|---|---|---|---|
| Isotope correction at MS1 | No | Lipid Identification Software | MS-DIAL |
| Isotope correction at MS2 | No | Data manipulation | Smoothing, Centroiding |
| MS1 verified by standard | Yes | Nomenclature for intact lipid molecule | Yes |
| MS2 verified by standard | Yes | Nomenclature for fragment ions | No |
| Background check at MS1 | Yes | Further identification remarks | - |

#### 2) DG[M+NH4]+ / Lipid quantification

| | | | |
|---|---|---|---|
| Quantitative | Yes | Limit of quantification | No |
| MS Level for quantification | MS1 | Normalization to reference | No |
| Internal lipid standard(s) MS1 | | Lipid Quantification Software | MS-DIAL |

| Internal standard | Endogenous subclass |
|---|---|
| DG 15:0_18:1(d7) | DG subclass |

| | | | |
|---|---|---|---|
| Type of quantification | Internal standard amount | Batch correction | No |
| Response correction | No | Further quantification remarks | - |
| Type I isotope correction | No | | |

#### 3) DGDG[M+CH3COO]- / Lipid identification

| | | | |
|---|---|---|---|
| Lipid class | DGDG | Background check at MS2 | No |
| Derivatization | - | Did you presume assumptions for identification? | No |
| MS Level for identification | MS1, MS2 | Check isomer overlap | No |
| Identification level | Molecular species level | RT verified by standard | Yes |
| Polarity mode | Negative | Separation of isobaric/isomeric interferece confirmed | Yes |
| Type of negative (precursor)ion | [M+CH3COO]- | Model for separation prediction | Yes |
| Fragments for identification | | Additional dimension/techniques | - |

| Fragment name |
|---|
| Fatty acid fragment |
| Neutral loss of fatty acids |

| | | | |
|---|---|---|---|
| Isotope correction at MS1 | No | Lipid Identification Software | MS-DIAL |
| Isotope correction at MS2 | No | Data manipulation | Smoothing, Centroiding |
| MS1 verified by standard | No | Nomenclature for intact lipid molecule | Yes |
| MS2 verified by standard | No | Nomenclature for fragment ions | No |
| Background check at MS1 | Yes | Further identification remarks | - |

#### 3) DGDG[M+CH3COO]- / Lipid quantification

| | | | |
|---|---|---|---|
| Quantitative | Yes | Limit of quantification | No |
| MS Level for quantification | MS1 | Normalization to reference | No |
| Internal lipid standard(s) MS1 | | Lipid Quantification Software | MS-DIAL |

| Internal standard | Endogenous subclass |
|---|---|
| LPC 18:1(d7) | DGDG subclass |

| | | | |
|---|---|---|---|
| Type of quantification | Internal standard amount | Batch correction | No |
| Response correction | No | Further quantification remarks | - |
| Type I isotope correction | No | | |

#### 4) Sulfonolipid (SL)[M-H]- / Lipid identification

| | | | |
|---|---|---|---|
| Lipid class | Sulfonolipid (SL) | Background check at MS2 | No |
| Derivatization | - | Did you presume assumptions for identification? | No |
| MS Level for identification | MS1, MS2 | Check isomer overlap | No |
| Identification level | Molecular species level | RT verified by standard | Yes |
| Polarity mode | Negative | Separation of isobaric/isomeric interferece confirmed | Yes |
| Type of negative (precursor)ion | [M-H]- | Model for separation prediction | Yes |
| Fragments for identification | | Additional dimension/techniques | - |

| Fragment name |
|---|
| Sulfite |
| Neutral loss of N-acyl chain |

| | | | |
|---|---|---|---|
| Isotope correction at MS1 | No | Lipid Identification Software | MS-DIAL |
| Isotope correction at MS2 | No | Data manipulation | Smoothing, Centroiding |
| MS1 verified by standard | No | Nomenclature for intact lipid molecule | Yes |
| MS2 verified by standard | No | Nomenclature for fragment ions | No |
| Background check at MS1 | Yes | Further identification remarks | - |

#### 4) Sulfonolipid (SL)[M-H]- / Lipid quantification

| | | | |
|---|---|---|---|
| Quantitative | Yes | Limit of quantification | No |
| MS Level for quantification | MS1 | Normalization to reference | No |
| Internal lipid standard(s) MS1 | | Lipid Quantification Software | MS-DIAL |

| Internal standard | Endogenous subclass |
|---|---|
| Cer 18:1;2O/15:0(d7) | SL subclass |

| | | | |
|---|---|---|---|
| Type of quantification | Internal standard amount | Batch correction | No |
| Response correction | No | Further quantification remarks | - |
| Type I isotope correction | No | | |

#### 5) Acyl diacylglyceryl glucuronide(ADGGA)[M-H]- / Lipid identification

| | | | |
|---|---|---|---|
| Lipid class | Acyl diacylglyceryl glucuronide(ADGGA) | Background check at MS2 | No |
| Derivatization | - | Did you presume assumptions for identification? | No |
| MS Level for identification | MS1, MS2 | Check isomer overlap | No |
| Identification level | Molecular species level | RT verified by standard | Yes |
| Polarity mode | Negative | Separation of isobaric/isomeric interferece confirmed | Yes |
| Type of negative (precursor)ion | [M-H]- | Model for separation prediction | Yes |
| Fragments for identification | | Additional dimension/techniques | - |

| Fragment name |
|---|
| Fatty acid fragment |

| | | | |
|---|---|---|---|
| Isotope correction at MS1 | No | Lipid Identification Software | MS-DIAL |
| Isotope correction at MS2 | No | Data manipulation | Smoothing, Centroiding |
| MS1 verified by standard | No | Nomenclature for intact lipid molecule | Yes |
| MS2 verified by standard | No | Nomenclature for fragment ions | No |
| Background check at MS1 | Yes | Further identification remarks | - |

#### 5) Acyl diacylglyceryl glucuronide(ADGGA)[M-H]- / Lipid quantification

| | | | |
|---|---|---|---|
| Quantitative | Yes | Limit of quantification | No |
| MS Level for quantification | MS1 | Normalization to reference | No |
| Internal lipid standard(s) MS1 | | Lipid Quantification Software | MS-DIAL |

| Internal standard | Endogenous subclass |
|---|---|
| LPC 18:1(d7) | ADGGA subclass |

| | | | |
|---|---|---|---|
| Type of quantification | Internal standard amount | Batch correction | No |
| Response correction | No | Further quantification remarks | - |
| Type I isotope correction | No | | |

#### 6) Acylcarnitine (CAR)[M+H]+ / Lipid identification

| | | | |
|---|---|---|---|
| Lipid class | Acylcarnitine (CAR) | Background check at MS2 | No |
| Derivatization | - | Did you presume assumptions for identification? | No |
| MS Level for identification | MS1, MS2 | Check isomer overlap | No |
| Identification level | Molecular species level | RT verified by standard | Yes |
| Polarity mode | Positive | Separation of isobaric/isomeric interferece confirmed | Yes |
| Type of positive (precursor)ion | [M+H]+ | Model for separation prediction | Yes |
| Fragments for identification | | Additional dimension/techniques | - |

| Fragment name |
|---|
| Characteristic fragment (C4H5O2+) |

| | | | |
|---|---|---|---|
| Isotope correction at MS1 | No | Lipid Identification Software | MS-DIAL |
| Isotope correction at MS2 | No | Data manipulation | Smoothing, Centroiding |
| MS1 verified by standard | No | Nomenclature for intact lipid molecule | Yes |
| MS2 verified by standard | No | Nomenclature for fragment ions | No |
| Background check at MS1 | Yes | Further identification remarks | - |

#### 6) Acylcarnitine (CAR)[M+H]+ / Lipid quantification

| | | | |
|---|---|---|---|
| Quantitative | Yes | Limit of quantification | No |
| MS Level for quantification | MS1 | Normalization to reference | No |
| Internal lipid standard(s) MS1 | | Lipid Quantification Software | MS-DIAL |

| Internal standard | Endogenous subclass |
|---|---|
| LPC 18:1(d7) | CAR subclass |

| | | | |
|---|---|---|---|
| Type of quantification | Internal standard amount | Batch correction | No |
| Response correction | No | Further quantification remarks | - |
| Type I isotope correction | No | | |

#### 7) Acylhexosyl brassicasterol (AHexBRS)[M+CH3COO]- / Lipid identification

| | | | |
|---|---|---|---|
| Lipid class | Acylhexosyl brassicasterol (AHexBRS) | Background check at MS2 | No |
| Derivatization | - | Did you presume assumptions for identification? | No |
| MS Level for identification | MS1, MS2 | Check isomer overlap | No |
| Identification level | Molecular species level | RT verified by standard | Yes |
| Polarity mode | Negative | Separation of isobaric/isomeric interferece confirmed | Yes |
| Type of negative (precursor)ion | [M+CH3COO]- | Model for separation prediction | Yes |
| Fragments for identification | | Additional dimension/techniques | - |

| Fragment name |
|---|
| Fatty acid fragment |

| | | | |
|---|---|---|---|
| Isotope correction at MS1 | No | Lipid Identification Software | MS-DIAL |
| Isotope correction at MS2 | No | Data manipulation | Smoothing, Centroiding |
| MS1 verified by standard | No | Nomenclature for intact lipid molecule | Yes |
| MS2 verified by standard | No | Nomenclature for fragment ions | No |
| Background check at MS1 | Yes | Further identification remarks | - |

#### 7) Acylhexosyl brassicasterol (AHexBRS)[M+CH3COO]- / Lipid quantification

| | | | |
|---|---|---|---|
| Quantitative | Yes | Limit of quantification | No |
| MS Level for quantification | MS1 | Normalization to reference | No |
| Internal lipid standard(s) MS1 | | Lipid Quantification Software | MS-DIAL |

| Internal standard | Endogenous subclass |
|---|---|
| CE 18:1(d7) | AHexBRS subclass |

| | | | |
|---|---|---|---|
| Type of quantification | Internal standard amount | Batch correction | No |
| Response correction | No | Further quantification remarks | - |
| Type I isotope correction | No | | |

#### 8) Acylhexosyl campesterol (AHexCAS)[M+CH3COO]- / Lipid identification

| | | | |
|---|---|---|---|
| Lipid class | Acylhexosyl campesterol (AHexCAS) | Background check at MS2 | No |
| Derivatization | - | Did you presume assumptions for identification? | No |
| MS Level for identification | MS1, MS2 | Check isomer overlap | No |
| Identification level | Molecular species level | RT verified by standard | Yes |
| Polarity mode | Negative | Separation of isobaric/isomeric interferece confirmed | Yes |
| Type of negative (precursor)ion | [M+CH3COO]- | Model for separation prediction | Yes |
| Fragments for identification | | Additional dimension/techniques | - |

| Fragment name |
|---|
| Fatty acid fragment |

| | | | |
|---|---|---|---|
| Isotope correction at MS1 | No | Lipid Identification Software | MS-DIAL |
| Isotope correction at MS2 | No | Data manipulation | Smoothing, Centroiding |
| MS1 verified by standard | No | Nomenclature for intact lipid molecule | Yes |
| MS2 verified by standard | No | Nomenclature for fragment ions | No |
| Background check at MS1 | Yes | Further identification remarks | - |

#### 8) Acylhexosyl campesterol (AHexCAS)[M+CH3COO]- / Lipid quantification

| | | | |
|---|---|---|---|
| Quantitative | Yes | Limit of quantification | No |
| MS Level for quantification | MS1 | Normalization to reference | No |
| Internal lipid standard(s) MS1 | | Lipid Quantification Software | MS-DIAL |

| Internal standard | Endogenous subclass |
|---|---|
| CE 18:1(d7) | AHexCAS subclass |

| | | | |
|---|---|---|---|
| Type of quantification | Internal standard amount | Batch correction | No |
| Response correction | No | Further quantification remarks | - |
| Type I isotope correction | No | | |

#### 9) Acylhexosyl cholesterol (AHexCS)[M+CH3COO]- / Lipid identification

| | | | |
|---|---|---|---|
| Lipid class | Acylhexosyl cholesterol (AHexCS) | Background check at MS2 | No |
| Derivatization | - | Did you presume assumptions for identification? | No |
| MS Level for identification | MS1, MS2 | Check isomer overlap | No |
| Identification level | Molecular species level | RT verified by standard | Yes |
| Polarity mode | Negative | Separation of isobaric/isomeric interferece confirmed | Yes |
| Type of negative (precursor)ion | [M+CH3COO]- | Model for separation prediction | Yes |
| Fragments for identification | | Additional dimension/techniques | - |

| Fragment name |
|---|
| Fatty acid fragment |

| | | | |
|---|---|---|---|
| Isotope correction at MS1 | No | Lipid Identification Software | MS-DIAL |
| Isotope correction at MS2 | No | Data manipulation | Smoothing, Centroiding |
| MS1 verified by standard | No | Nomenclature for intact lipid molecule | Yes |
| MS2 verified by standard | No | Nomenclature for fragment ions | No |
| Background check at MS1 | Yes | Further identification remarks | - |

#### 9) Acylhexosyl cholesterol (AHexCS)[M+CH3COO]- / Lipid quantification

| | | | |
|---|---|---|---|
| Quantitative | Yes | Limit of quantification | No |
| MS Level for quantification | MS1 | Normalization to reference | No |
| Internal lipid standard(s) MS1 | | Lipid Quantification Software | MS-DIAL |

| Internal standard | Endogenous subclass |
|---|---|
| CE 18:1(d7) | AHexCS subclass |

| | | | |
|---|---|---|---|
| Type of quantification | Internal standard amount | Batch correction | No |
| Response correction | No | Further quantification remarks | - |
| Type I isotope correction | No | | |

#### 10) Acylhexosyl sitosterol (AHexSIS)[M+CH3COO]- / Lipid identification

| | | | |
|---|---|---|---|
| Lipid class | Acylhexosyl sitosterol (AHexSIS) | Background check at MS2 | No |
| Derivatization | - | Did you presume assumptions for identification? | No |
| MS Level for identification | MS1, MS2 | Check isomer overlap | No |
| Identification level | Molecular species level | RT verified by standard | Yes |
| Polarity mode | Negative | Separation of isobaric/isomeric interferece confirmed | Yes |
| Type of negative (precursor)ion | [M+CH3COO]- | Model for separation prediction | Yes |
| Fragments for identification | | Additional dimension/techniques | - |

| Fragment name |
|---|
| Fatty acid fragment |

| | | | |
|---|---|---|---|
| Isotope correction at MS1 | No | Lipid Identification Software | MS-DIAL |
| Isotope correction at MS2 | No | Data manipulation | Smoothing, Centroiding |
| MS1 verified by standard | No | Nomenclature for intact lipid molecule | Yes |
| MS2 verified by standard | No | Nomenclature for fragment ions | No |
| Background check at MS1 | Yes | Further identification remarks | - |

#### 10) Acylhexosyl sitosterol (AHexSIS)[M+CH3COO]- / Lipid quantification

| | | | |
|---|---|---|---|
| Quantitative | Yes | Limit of quantification | No |
| MS Level for quantification | MS1 | Normalization to reference | No |
| Internal lipid standard(s) MS1 | | Lipid Quantification Software | MS-DIAL |

| Internal standard | Endogenous subclass |
|---|---|
| CE 18:1(d7) | AHexSIS subclass |

| | | | |
|---|---|---|---|
| Type of quantification | Internal standard amount | Batch correction | No |
| Response correction | No | Further quantification remarks | - |
| Type I isotope correction | No | | |

#### 11) Acylhexosyl stigmasterol (AHexSTS)[M+CH3COO]- / Lipid identification

| | | | |
|---|---|---|---|
| Lipid class | Acylhexosyl stigmasterol (AHexSTS) | Background check at MS2 | No |
| Derivatization | - | Did you presume assumptions for identification? | No |
| MS Level for identification | MS1, MS2 | Check isomer overlap | No |
| Identification level | Molecular species level | RT verified by standard | Yes |
| Polarity mode | Negative | Separation of isobaric/isomeric interferece confirmed | Yes |
| Type of negative (precursor)ion | [M+CH3COO]- | Model for separation prediction | Yes |
| Fragments for identification | | Additional dimension/techniques | - |

| Fragment name |
|---|
| Fatty acid fragment |

| | | | |
|---|---|---|---|
| Isotope correction at MS1 | No | Lipid Identification Software | MS-DIAL |
| Isotope correction at MS2 | No | Data manipulation | Smoothing, Centroiding |
| MS1 verified by standard | No | Nomenclature for intact lipid molecule | Yes |
| MS2 verified by standard | No | Nomenclature for fragment ions | No |
| Background check at MS1 | Yes | Further identification remarks | - |

#### 11) Acylhexosyl stigmasterol (AHexSTS)[M+CH3COO]- / Lipid quantification

| | | | |
|---|---|---|---|
| Quantitative | Yes | Limit of quantification | No |
| MS Level for quantification | MS1 | Normalization to reference | No |
| Internal lipid standard(s) MS1 | | Lipid Quantification Software | MS-DIAL |

| Internal standard | Endogenous subclass |
|---|---|
| CE 18:1(d7) | AHexSTS subclass |

| | | | |
|---|---|---|---|
| Type of quantification | Internal standard amount | Batch correction | No |
| Response correction | No | Further quantification remarks | - |
| Type I isotope correction | No | | |

#### 12) Acylhexosylceramide (AHexCer)[M+CH3COO]- / Lipid identification

| | | | |
|---|---|---|---|
| Lipid class | Acylhexosylceramide (AHexCer) | Background check at MS2 | No |
| Derivatization | - | Did you presume assumptions for identification? | No |
| MS Level for identification | MS1, MS2 | Check isomer overlap | No |
| Identification level | Molecular species level | RT verified by standard | Yes |
| Polarity mode | Negative | Separation of isobaric/isomeric interferece confirmed | Yes |
| Type of negative (precursor)ion | [M+CH3COO]- | Model for separation prediction | Yes |
| Fragments for identification | | Additional dimension/techniques | - |

| Fragment name |
|---|
| Neutral loss of fatty acid |
| Neutral loss of fatty acid and hexose |
| Fatty acid fragment |
| Neutral loss of N- acyl |
| Neutral loss of fatty acid and H2O |
| Neutral loss of N-acyl and H2O |

| | | | |
|---|---|---|---|
| Isotope correction at MS1 | No | Lipid Identification Software | MS-DIAL |
| Isotope correction at MS2 | No | Data manipulation | Smoothing, Centroiding |
| MS1 verified by standard | No | Nomenclature for intact lipid molecule | Yes |
| MS2 verified by standard | No | Nomenclature for fragment ions | No |
| Background check at MS1 | Yes | Further identification remarks | - |

#### 12) Acylhexosylceramide (AHexCer)[M+CH3COO]- / Lipid quantification

| | | | |
|---|---|---|---|
| Quantitative | Yes | Limit of quantification | No |
| MS Level for quantification | MS1 | Normalization to reference | No |
| Internal lipid standard(s) MS1 | | Lipid Quantification Software | MS-DIAL |

| Internal standard | Endogenous subclass |
|---|---|
| Cer 18:1;2O/15:0(d7) | AHexCer subclass |

| | | | |
|---|---|---|---|
| Type of quantification | Internal standard amount | Batch correction | No |
| Response correction | No | Further quantification remarks | - |
| Type I isotope correction | No | | |

#### 13) Acylsphingomyelin (ASM)[M+CH3COO]- / Lipid identification

| | | | |
|---|---|---|---|
| Lipid class | Acylsphingomyelin (ASM) | Background check at MS2 | No |
| Derivatization | - | Did you presume assumptions for identification? | No |
| MS Level for identification | MS1, MS2 | Check isomer overlap | No |
| Identification level | Molecular species level | RT verified by standard | Yes |
| Polarity mode | Negative | Separation of isobaric/isomeric interferece confirmed | Yes |
| Type of negative (precursor)ion | [M+CH3COO]- | Model for separation prediction | Yes |
| Fragments for identification | | Additional dimension/techniques | - |

| Fragment name |
|---|
| Neutral loss of methyl |
| Neutral loss of methyl and fatty acid and H2O |
| Fatty acid fragment |
| Acyl amide |

| | | | |
|---|---|---|---|
| Isotope correction at MS1 | No | Lipid Identification Software | MS-DIAL |
| Isotope correction at MS2 | No | Data manipulation | Smoothing, Centroiding |
| MS1 verified by standard | No | Nomenclature for intact lipid molecule | Yes |
| MS2 verified by standard | No | Nomenclature for fragment ions | No |
| Background check at MS1 | Yes | Further identification remarks | - |

#### 13) Acylsphingomyelin (ASM)[M+CH3COO]- / Lipid quantification

| | | | |
|---|---|---|---|
| Quantitative | Yes | Limit of quantification | No |
| MS Level for quantification | MS1 | Normalization to reference | No |
| Internal lipid standard(s) MS1 | | Lipid Quantification Software | MS-DIAL |

| Internal standard | Endogenous subclass |
|---|---|
| SM 18:1;2O/18:1(d9) | ASM subclass |

| | | | |
|---|---|---|---|
| Type of quantification | Internal standard amount | Batch correction | No |
| Response correction | No | Further quantification remarks | - |
| Type I isotope correction | No | | |

#### 14) BMP[M+NH4]+ / Lipid identification

| | | | |
|---|---|---|---|
| Lipid class | BMP | Background check at MS2 | No |
| Derivatization | - | Did you presume assumptions for identification? | No |
| MS Level for identification | MS1, MS2 | Check isomer overlap | No |
| Identification level | Molecular species level | RT verified by standard | Yes |
| Polarity mode | Positive | Separation of isobaric/isomeric interferece confirmed | Yes |
| Type of positive (precursor)ion | [M+NH4]+ | Model for separation prediction | Yes |
| Fragments for identification | | Additional dimension/techniques | - |

| Fragment name |
|---|
| Dehydro-monoacyl glycerols |
| Neutral loss of glycerophosphate |

| | | | |
|---|---|---|---|
| Isotope correction at MS1 | No | Lipid Identification Software | MS-DIAL |
| Isotope correction at MS2 | No | Data manipulation | Smoothing, Centroiding |
| MS1 verified by standard | No | Nomenclature for intact lipid molecule | Yes |
| MS2 verified by standard | No | Nomenclature for fragment ions | No |
| Background check at MS1 | Yes | Further identification remarks | - |

#### 14) BMP[M+NH4]+ / Lipid quantification

| | | | |
|---|---|---|---|
| Quantitative | Yes | Limit of quantification | No |
| MS Level for quantification | MS1 | Normalization to reference | No |
| Internal lipid standard(s) MS1 | | Lipid Quantification Software | MS-DIAL |

| Internal standard | Endogenous subclass |
|---|---|
| PG 15:0_18:1(d7) | BMP subclass |

| | | | |
|---|---|---|---|
| Type of quantification | Internal standard amount | Batch correction | No |
| Response correction | No | Further quantification remarks | - |
| Type I isotope correction | No | | |

#### 15) Brassicasterol[M+NH4]+ / Lipid identification

| | | | |
|---|---|---|---|
| Lipid class | Brassicasterol | Check isomer overlap | No |
| Derivatization | - | RT verified by standard | Yes |
| MS Level for identification | MS1 | Separation of isobaric/isomeric interferece confirmed | Yes |
| Identification level | Species level | Model for separation prediction | Yes |
| Polarity mode | Positive | Additional dimension/techniques | - |
| Type of positive (precursor)ion | [M+NH4]+ | Lipid Identification Software | MS-DIAL |
| Isotope correction at MS1 | No | Data manipulation | Smoothing, Centroiding |
| MS1 verified by standard | No | Nomenclature for intact lipid molecule | Yes |
| Background check at MS1 | Yes | Further identification remarks | - |
| Did you presume assumptions for identification? | No | | |

#### 15) Brassicasterol[M+NH4]+ / Lipid quantification

| | | | |
|---|---|---|---|
| Quantitative | Yes | Limit of quantification | No |
| MS Level for quantification | MS1 | Normalization to reference | No |
| Internal lipid standard(s) MS1 | | Lipid Quantification Software | MS-DIAL |

| Internal standard | Endogenous subclass |
|---|---|
| CE 18:1(d7) | Brassicasterol |

| | | | |
|---|---|---|---|
| Type of quantification | Internal standard amount | Batch correction | No |
| Response correction | No | Further quantification remarks | - |
| Type I isotope correction | No | | |

#### 16) Brassicasterol ester (BRSE)[M+NH4]+ / Lipid identification

| | | | |
|---|---|---|---|
| Lipid class | Brassicasterol ester (BRSE) | Background check at MS2 | No |
| Derivatization | - | Did you presume assumptions for identification? | No |
| MS Level for identification | MS1, MS2 | Check isomer overlap | No |
| Identification level | Molecular species level | RT verified by standard | Yes |
| Polarity mode | Positive | Separation of isobaric/isomeric interferece confirmed | Yes |
| Type of positive (precursor)ion | [M+NH4]+ | Model for separation prediction | Yes |
| Fragments for identification | | Additional dimension/techniques | - |

| Fragment name |
|---|
| Neutral loss of fatty acid |

| | | | |
|---|---|---|---|
| Isotope correction at MS1 | No | Lipid Identification Software | MS-DIAL |
| Isotope correction at MS2 | No | Data manipulation | Smoothing, Centroiding |
| MS1 verified by standard | No | Nomenclature for intact lipid molecule | Yes |
| MS2 verified by standard | No | Nomenclature for fragment ions | No |
| Background check at MS1 | Yes | Further identification remarks | - |

#### 16) Brassicasterol ester (BRSE)[M+NH4]+ / Lipid quantification

| | | | |
|---|---|---|---|
| Quantitative | Yes | Limit of quantification | No |
| MS Level for quantification | MS1 | Normalization to reference | No |
| Internal lipid standard(s) MS1 | | Lipid Quantification Software | MS-DIAL |

| Internal standard | Endogenous subclass |
|---|---|
| CE 18:1(d7) | BRSE subclass |

| | | | |
|---|---|---|---|
| Type of quantification | Internal standard amount | Batch correction | No |
| Response correction | No | Further quantification remarks | - |
| Type I isotope correction | No | | |

#### 17) Campesterol ester (CASE)[M+NH4]+ / Lipid identification

| | | | |
|---|---|---|---|
| Lipid class | Campesterol ester (CASE) | Background check at MS2 | No |
| Derivatization | - | Did you presume assumptions for identification? | No |
| MS Level for identification | MS1, MS2 | Check isomer overlap | No |
| Identification level | Molecular species level | RT verified by standard | Yes |
| Polarity mode | Positive | Separation of isobaric/isomeric interferece confirmed | Yes |
| Type of positive (precursor)ion | [M+NH4]+ | Model for separation prediction | Yes |
| Fragments for identification | | Additional dimension/techniques | - |

| Fragment name |
|---|
| Neutral loss of fatty acid |

| | | | |
|---|---|---|---|
| Isotope correction at MS1 | No | Lipid Identification Software | MS-DIAL |
| Isotope correction at MS2 | No | Data manipulation | Smoothing, Centroiding |
| MS1 verified by standard | No | Nomenclature for intact lipid molecule | Yes |
| MS2 verified by standard | No | Nomenclature for fragment ions | No |
| Background check at MS1 | Yes | Further identification remarks | - |

#### 17) Campesterol ester (CASE)[M+NH4]+ / Lipid quantification

| | | | |
|---|---|---|---|
| Quantitative | Yes | Limit of quantification | No |
| MS Level for quantification | MS1 | Normalization to reference | No |
| Internal lipid standard(s) MS1 | | Lipid Quantification Software | MS-DIAL |

| Internal standard | Endogenous subclass |
|---|---|
| CE 18:1(d7) | CASE subclass |

| | | | |
|---|---|---|---|
| Type of quantification | Internal standard amount | Batch correction | No |
| Response correction | No | Further quantification remarks | - |
| Type I isotope correction | No | | |

#### 18) CL[M-H]- / Lipid identification

| | | | |
|---|---|---|---|
| Lipid class | CL | Background check at MS2 | No |
| Derivatization | - | Did you presume assumptions for identification? | No |
| MS Level for identification | MS1, MS2 | Check isomer overlap | No |
| Identification level | Molecular species level | RT verified by standard | Yes |
| Polarity mode | Negative | Separation of isobaric/isomeric interferece confirmed | Yes |
| Type of negative (precursor)ion | [M-H]- | Model for separation prediction | Yes |
| Fragments for identification | | Additional dimension/techniques | - |

| Fragment name |
|---|
| Phosphoglycerol - H2O |
| Fatty acid fragment |
| Phosphatidic acid |

| | | | |
|---|---|---|---|
| Isotope correction at MS1 | No | Lipid Identification Software | MS-DIAL |
| Isotope correction at MS2 | No | Data manipulation | Smoothing, Centroiding |
| MS1 verified by standard | No | Nomenclature for intact lipid molecule | Yes |
| MS2 verified by standard | No | Nomenclature for fragment ions | No |
| Background check at MS1 | Yes | Further identification remarks | - |

#### 18) CL[M-H]- / Lipid quantification

| | | | |
|---|---|---|---|
| Quantitative | Yes | Limit of quantification | No |
| MS Level for quantification | MS1 | Normalization to reference | No |
| Internal lipid standard(s) MS1 | | Lipid Quantification Software | MS-DIAL |

| Internal standard | Endogenous subclass |
|---|---|
| PG 15:0_18:1(d7) | CL subclass |

| | | | |
|---|---|---|---|
| Type of quantification | Internal standard amount | Batch correction | No |
| Response correction | No | Further quantification remarks | - |
| Type I isotope correction | No | | |

#### 19) Ceramide alpha-hydroxy fatty acid-dihydrosphingosine (Cer_ADS)[M+CH3COO]- / Lipid identification

| | | | |
|---|---|---|---|
| Lipid class | Ceramide alpha-hydroxy fatty acid-dihydrosphingosine (Cer_ADS) | Background check at MS2 | No |
| Derivatization | - | Did you presume assumptions for identification? | No |
| MS Level for identification | MS1, MS2 | Check isomer overlap | No |
| Identification level | Molecular species level | RT verified by standard | Yes |
| Polarity mode | Negative | Separation of isobaric/isomeric interferece confirmed | Yes |
| Type of negative (precursor)ion | [M+CH3COO]- | Model for separation prediction | Yes |
| Fragments for identification | | Additional dimension/techniques | - |

| Fragment name |
|---|
| Neutral loss of CH4O2 |
| Neutral loss of H2O |
| Sphinganine |
| Sphinganine -H2O fragment |
| Oxidized acyl -2H fragment |
| Oxidized acyl -CH2O fragment |
| Sphinganine -C2H7NO fragment |

| | | | |
|---|---|---|---|
| Isotope correction at MS1 | No | Lipid Identification Software | MS-DIAL |
| Isotope correction at MS2 | No | Data manipulation | Smoothing, Centroiding |
| MS1 verified by standard | No | Nomenclature for intact lipid molecule | Yes |
| MS2 verified by standard | No | Nomenclature for fragment ions | No |
| Background check at MS1 | Yes | Further identification remarks | - |

#### 19) Ceramide alpha-hydroxy fatty acid-dihydrosphingosine (Cer_ADS)[M+CH3COO]- / Lipid quantification

| | | | |
|---|---|---|---|
| Quantitative | Yes | Limit of quantification | No |
| MS Level for quantification | MS1 | Normalization to reference | No |
| Internal lipid standard(s) MS1 | | Lipid Quantification Software | MS-DIAL |

| Internal standard | Endogenous subclass |
|---|---|
| Cer 18:1;2O/15:0(d7) | Cer_ADS subclass |

| | | | |
|---|---|---|---|
| Type of quantification | Internal standard amount | Batch correction | No |
| Response correction | No | Further quantification remarks | - |
| Type I isotope correction | No | | |

#### 20) Ceramide alpha-hydroxy fatty acid-phytospingosine (Cer_AP)[M+CH3COO]- / Lipid identification

| | | | |
|---|---|---|---|
| Lipid class | Ceramide alpha-hydroxy fatty acid-phytospingosine (Cer_AP) | Background check at MS2 | No |
| Derivatization | - | Did you presume assumptions for identification? | No |
| MS Level for identification | MS1, MS2 | Check isomer overlap | No |
| Identification level | Molecular species level | RT verified by standard | Yes |
| Polarity mode | Negative | Separation of isobaric/isomeric interferece confirmed | Yes |
| Type of negative (precursor)ion | [M+CH3COO]- | Model for separation prediction | Yes |
| Fragments for identification | | Additional dimension/techniques | - |

| Fragment name |
|---|
| Oxidized fatty acyl -CH2O fragment |
| Oxidized fatty acyl +O fragment |
| Oxidized fatty acyl +C3H5NO fragment |

| | | | |
|---|---|---|---|
| Isotope correction at MS1 | No | Lipid Identification Software | MS-DIAL |
| Isotope correction at MS2 | No | Data manipulation | Smoothing, Centroiding |
| MS1 verified by standard | No | Nomenclature for intact lipid molecule | Yes |
| MS2 verified by standard | No | Nomenclature for fragment ions | No |
| Background check at MS1 | Yes | Further identification remarks | - |

#### 20) Ceramide alpha-hydroxy fatty acid-phytospingosine (Cer_AP)[M+CH3COO]- / Lipid quantification

| | | | |
|---|---|---|---|
| Quantitative | Yes | Limit of quantification | No |
| MS Level for quantification | MS1 | Normalization to reference | No |
| Internal lipid standard(s) MS1 | | Lipid Quantification Software | MS-DIAL |

| Internal standard | Endogenous subclass |
|---|---|
| Cer 18:1;2O/15:0(d7) | Cer_AP subclass |

| | | | |
|---|---|---|---|
| Type of quantification | Internal standard amount | Batch correction | No |
| Response correction | No | Further quantification remarks | - |
| Type I isotope correction | No | | |

#### 21) Ceramide alpha-hydroxy fatty acid-sphingosine (Cer_AS)[M+CH3COO]- / Lipid identification

| | | | |
|---|---|---|---|
| Lipid class | Ceramide alpha-hydroxy fatty acid-sphingosine (Cer_AS) | Background check at MS2 | No |
| Derivatization | - | Did you presume assumptions for identification? | No |
| MS Level for identification | MS1, MS2 | Check isomer overlap | No |
| Identification level | Molecular species level | RT verified by standard | Yes |
| Polarity mode | Negative | Separation of isobaric/isomeric interferece confirmed | Yes |
| Type of negative (precursor)ion | [M+CH3COO]- | Model for separation prediction | Yes |
| Fragments for identification | | Additional dimension/techniques | - |

| Fragment name |
|---|
| Neutral loss of CH4O2 |
| Neutral loss of H2O |
| Sphingosine -H2O fragment |
| Sphingosine -C2H7NO fragment |
| Oxidized fatty acyl -CH2O fragment |
| Oxidized fatty acyl -2H fragment |

| | | | |
|---|---|---|---|
| Isotope correction at MS1 | No | Lipid Identification Software | MS-DIAL |
| Isotope correction at MS2 | No | Data manipulation | Smoothing, Centroiding |
| MS1 verified by standard | No | Nomenclature for intact lipid molecule | Yes |
| MS2 verified by standard | No | Nomenclature for fragment ions | No |
| Background check at MS1 | Yes | Further identification remarks | - |

#### 21) Ceramide alpha-hydroxy fatty acid-sphingosine (Cer_AS)[M+CH3COO]- / Lipid quantification

| | | | |
|---|---|---|---|
| Quantitative | Yes | Limit of quantification | No |
| MS Level for quantification | MS1 | Normalization to reference | No |
| Internal lipid standard(s) MS1 | | Lipid Quantification Software | MS-DIAL |

| Internal standard | Endogenous subclass |
|---|---|
| Cer 18:1;2O/15:0(d7) | Cer_AS subclass |

| | | | |
|---|---|---|---|
| Type of quantification | Internal standard amount | Batch correction | No |
| Response correction | No | Further quantification remarks | - |
| Type I isotope correction | No | | |

#### 22) Ceramide beta-hydroxy fatty acid-dihydrosphingosine (Cer_BDS)[M-H]- / Lipid identification

| | | | |
|---|---|---|---|
| Lipid class | Ceramide beta-hydroxy fatty acid-dihydrosphingosine (Cer_BDS) | Background check at MS2 | No |
| Derivatization | - | Did you presume assumptions for identification? | No |
| MS Level for identification | MS1, MS2 | Check isomer overlap | No |
| Identification level | Molecular species level | RT verified by standard | Yes |
| Polarity mode | Negative | Separation of isobaric/isomeric interferece confirmed | Yes |
| Type of negative (precursor)ion | [M-H]- | Model for separation prediction | Yes |
| Fragments for identification | | Additional dimension/techniques | - |

| Fragment name |
|---|
| Sphinganine |
| Sphinganine +C2H2O fragment |
| Sphinganine +C2H2O -CH4O fragment |
| Sphinganine -C2H7NO fragment |

| | | | |
|---|---|---|---|
| Isotope correction at MS1 | No | Lipid Identification Software | MS-DIAL |
| Isotope correction at MS2 | No | Data manipulation | Smoothing, Centroiding |
| MS1 verified by standard | No | Nomenclature for intact lipid molecule | Yes |
| MS2 verified by standard | No | Nomenclature for fragment ions | No |
| Background check at MS1 | Yes | Further identification remarks | - |

#### 22) Ceramide beta-hydroxy fatty acid-dihydrosphingosine (Cer_BDS)[M-H]- / Lipid quantification

| | | | |
|---|---|---|---|
| Quantitative | Yes | Limit of quantification | No |
| MS Level for quantification | MS1 | Normalization to reference | No |
| Internal lipid standard(s) MS1 | | Lipid Quantification Software | MS-DIAL |

| Internal standard | Endogenous subclass |
|---|---|
| Cer 18:1;2O/15:0(d7) | Cer_BDS subclass |

| | | | |
|---|---|---|---|
| Type of quantification | Internal standard amount | Batch correction | No |
| Response correction | No | Further quantification remarks | - |
| Type I isotope correction | No | | |

#### 23) Ceramide beta-hydroxy fatty acid-sphingosine (Cer_BS)[M-H]- / Lipid identification

| | | | |
|---|---|---|---|
| Lipid class | Ceramide beta-hydroxy fatty acid-sphingosine (Cer_BS) | Background check at MS2 | No |
| Derivatization | - | Did you presume assumptions for identification? | No |
| MS Level for identification | MS1, MS2 | Check isomer overlap | No |
| Identification level | Molecular species level | RT verified by standard | Yes |
| Polarity mode | Negative | Separation of isobaric/isomeric interferece confirmed | Yes |
| Type of negative (precursor)ion | [M-H]- | Model for separation prediction | Yes |
| Fragments for identification | | Additional dimension/techniques | - |

| Fragment name |
|---|
| Sphingosine -C2H7NO fragment |
| Sphingosine +C2H2O fragment |
| Sphinganine +C2H2O -CH2O fragment |

| | | | |
|---|---|---|---|
| Isotope correction at MS1 | No | Lipid Identification Software | MS-DIAL |
| Isotope correction at MS2 | No | Data manipulation | Smoothing, Centroiding |
| MS1 verified by standard | No | Nomenclature for intact lipid molecule | Yes |
| MS2 verified by standard | No | Nomenclature for fragment ions | No |
| Background check at MS1 | Yes | Further identification remarks | - |

#### 23) Ceramide beta-hydroxy fatty acid-sphingosine (Cer_BS)[M-H]- / Lipid quantification

| | | | |
|---|---|---|---|
| Quantitative | Yes | Limit of quantification | No |
| MS Level for quantification | MS1 | Normalization to reference | No |
| Internal lipid standard(s) MS1 | | Lipid Quantification Software | MS-DIAL |

| Internal standard | Endogenous subclass |
|---|---|
| Cer 18:1;2O/15:0(d7) | Cer_BS subclass |

| | | | |
|---|---|---|---|
| Type of quantification | Internal standard amount | Batch correction | No |
| Response correction | No | Further quantification remarks | - |
| Type I isotope correction | No | | |

#### 24) Ceramide Esterified beta-hydroxy fatty acid-dihydrosphingosine (Cer_EBDS)[M+CH3COO]- / Lipid identification

| | | | |
|---|---|---|---|
| Lipid class | Ceramide Esterified beta-hydroxy fatty acid-dihydrosphingosine (Cer_EBDS) | Background check at MS2 | No |
| Derivatization | - | Did you presume assumptions for identification? | No |
| MS Level for identification | MS1, MS2 | Check isomer overlap | No |
| Identification level | Molecular species level | RT verified by standard | Yes |
| Polarity mode | Negative | Separation of isobaric/isomeric interferece confirmed | Yes |
| Type of negative (precursor)ion | [M+CH3COO]- | Model for separation prediction | Yes |
| Fragments for identification | | Additional dimension/techniques | - |

| Fragment name |
|---|
| Sphingosine +C2H2O fragment |
| Fatty acid fragment |
| Neutral loss of fatty acyl and H2O |

| | | | |
|---|---|---|---|
| Isotope correction at MS1 | No | Lipid Identification Software | MS-DIAL |
| Isotope correction at MS2 | No | Data manipulation | Smoothing, Centroiding |
| MS1 verified by standard | No | Nomenclature for intact lipid molecule | Yes |
| MS2 verified by standard | No | Nomenclature for fragment ions | No |
| Background check at MS1 | Yes | Further identification remarks | - |

#### 24) Ceramide Esterified beta-hydroxy fatty acid-dihydrosphingosine (Cer_EBDS)[M+CH3COO]- / Lipid quantification

| | | | |
|---|---|---|---|
| Quantitative | Yes | Limit of quantification | No |
| MS Level for quantification | MS1 | Normalization to reference | No |
| Internal lipid standard(s) MS1 | | Lipid Quantification Software | MS-DIAL |

| Internal standard | Endogenous subclass |
|---|---|
| Cer 18:1;2O/15:0(d7) | Cer_EBDS subclass |

| | | | |
|---|---|---|---|
| Type of quantification | Internal standard amount | Batch correction | No |
| Response correction | No | Further quantification remarks | - |
| Type I isotope correction | No | | |

#### 25) Ceramide Esterified omega-hydroxy fatty acid-dihydrosphingosine (Cer_EODS)[M-H]- / Lipid identification

| | | | |
|---|---|---|---|
| Lipid class | Ceramide Esterified omega-hydroxy fatty acid-dihydrosphingosine (Cer_EODS) | Background check at MS2 | No |
| Derivatization | - | Did you presume assumptions for identification? | No |
| MS Level for identification | MS1, MS2 | Check isomer overlap | No |
| Identification level | Molecular species level | RT verified by standard | Yes |
| Polarity mode | Negative | Separation of isobaric/isomeric interferece confirmed | Yes |
| Type of negative (precursor)ion | [M-H]- | Model for separation prediction | Yes |
| Fragments for identification | | Additional dimension/techniques | - |

| Fragment name |
|---|
| Fatty acid fragment |
| Neutral loss of fatty acyl |
| Acyl amide |

| | | | |
|---|---|---|---|
| Isotope correction at MS1 | No | Lipid Identification Software | MS-DIAL |
| Isotope correction at MS2 | No | Data manipulation | Smoothing, Centroiding |
| MS1 verified by standard | No | Nomenclature for intact lipid molecule | Yes |
| MS2 verified by standard | No | Nomenclature for fragment ions | No |
| Background check at MS1 | Yes | Further identification remarks | - |

#### 25) Ceramide Esterified omega-hydroxy fatty acid-dihydrosphingosine (Cer_EODS)[M-H]- / Lipid quantification

| | | | |
|---|---|---|---|
| Quantitative | Yes | Limit of quantification | No |
| MS Level for quantification | MS1 | Normalization to reference | No |
| Internal lipid standard(s) MS1 | | Lipid Quantification Software | MS-DIAL |

| Internal standard | Endogenous subclass |
|---|---|
| Cer 18:1;2O/15:0(d7) | Cer_EODS subclass |

| | | | |
|---|---|---|---|
| Type of quantification | Internal standard amount | Batch correction | No |
| Response correction | No | Further quantification remarks | - |
| Type I isotope correction | No | | |

#### 26) Ceramide Esterified omega-hydroxy fatty acid-sphingosine (Cer_EOS)[M+CH3COO]- / Lipid identification

| | | | |
|---|---|---|---|
| Lipid class | Ceramide Esterified omega-hydroxy fatty acid-sphingosine (Cer_EOS) | Background check at MS2 | No |
| Derivatization | - | Did you presume assumptions for identification? | No |
| MS Level for identification | MS1, MS2 | Check isomer overlap | No |
| Identification level | Molecular species level | RT verified by standard | Yes |
| Polarity mode | Negative | Separation of isobaric/isomeric interferece confirmed | Yes |
| Type of negative (precursor)ion | [M+CH3COO]- | Model for separation prediction | Yes |
| Fragments for identification | | Additional dimension/techniques | - |

| Fragment name |
|---|
| Acyl amide |
| Fatty acid fragment |
| Neutral loss of fatty acyl |

| | | | |
|---|---|---|---|
| Isotope correction at MS1 | No | Lipid Identification Software | MS-DIAL |
| Isotope correction at MS2 | No | Data manipulation | Smoothing, Centroiding |
| MS1 verified by standard | No | Nomenclature for intact lipid molecule | Yes |
| MS2 verified by standard | No | Nomenclature for fragment ions | No |
| Background check at MS1 | Yes | Further identification remarks | - |

#### 26) Ceramide Esterified omega-hydroxy fatty acid-sphingosine (Cer_EOS)[M+CH3COO]- / Lipid quantification

| | | | |
|---|---|---|---|
| Quantitative | Yes | Limit of quantification | No |
| MS Level for quantification | MS1 | Normalization to reference | No |
| Internal lipid standard(s) MS1 | | Lipid Quantification Software | MS-DIAL |

| Internal standard | Endogenous subclass |
|---|---|
| Cer 18:1;2O/15:0(d7) | Cer_EOS subclass |

| | | | |
|---|---|---|---|
| Type of quantification | Internal standard amount | Batch correction | No |
| Response correction | No | Further quantification remarks | - |
| Type I isotope correction | No | | |

#### 27) Ceramide non-hydroxyfatty acid-dihydrosphingosine (Cer_NDS)[M+CH3COO]- / Lipid identification

| | | | |
|---|---|---|---|
| Lipid class | Ceramide non-hydroxyfatty acid-dihydrosphingosine (Cer_NDS) | Background check at MS2 | No |
| Derivatization | - | Did you presume assumptions for identification? | No |
| MS Level for identification | MS1, MS2 | Check isomer overlap | No |
| Identification level | Molecular species level | RT verified by standard | Yes |
| Polarity mode | Negative | Separation of isobaric/isomeric interferece confirmed | Yes |
| Type of negative (precursor)ion | [M+CH3COO]- | Model for separation prediction | Yes |
| Fragments for identification | | Additional dimension/techniques | - |

| Fragment name |
|---|
| Neutral loss of CH4O |
| Neutral loss of CH4O2 |
| Sphinganine -C2H7NO fragment |
| Fatty acyl +C2H3N fragment |
| Fatty acyl -2H fragment |

| | | | |
|---|---|---|---|
| Isotope correction at MS1 | No | Lipid Identification Software | MS-DIAL |
| Isotope correction at MS2 | No | Data manipulation | Smoothing, Centroiding |
| MS1 verified by standard | No | Nomenclature for intact lipid molecule | Yes |
| MS2 verified by standard | No | Nomenclature for fragment ions | No |
| Background check at MS1 | Yes | Further identification remarks | - |

#### 27) Ceramide non-hydroxyfatty acid-dihydrosphingosine (Cer_NDS)[M+CH3COO]- / Lipid quantification

| | | | |
|---|---|---|---|
| Quantitative | Yes | Limit of quantification | No |
| MS Level for quantification | MS1 | Normalization to reference | No |
| Internal lipid standard(s) MS1 | | Lipid Quantification Software | MS-DIAL |

| Internal standard | Endogenous subclass |
|---|---|
| Cer 18:1;2O/15:0(d7) | Cer_NDS subclass |

| | | | |
|---|---|---|---|
| Type of quantification | Internal standard amount | Batch correction | No |
| Response correction | No | Further quantification remarks | - |
| Type I isotope correction | No | | |

#### 28) Ceramide non-hydroxyfatty acid-phytospingosine (Cer_NP)[M+CH3COO]- / Lipid identification

| | | | |
|---|---|---|---|
| Lipid class | Ceramide non-hydroxyfatty acid-phytospingosine (Cer_NP) | Background check at MS2 | No |
| Derivatization | - | Did you presume assumptions for identification? | No |
| MS Level for identification | MS1, MS2 | Check isomer overlap | No |
| Identification level | Molecular species level | RT verified by standard | Yes |
| Polarity mode | Negative | Separation of isobaric/isomeric interferece confirmed | Yes |
| Type of negative (precursor)ion | [M+CH3COO]- | Model for separation prediction | Yes |
| Fragments for identification | | Additional dimension/techniques | - |

| Fragment name |
|---|
| Neutral loss of H2O |
| Neutral loss of 2H2O |
| Sphinganine -CH7NO fragment |
| Fatty acyl +C3H5NO fragment |
| Acyl amide |

| | | | |
|---|---|---|---|
| Isotope correction at MS1 | No | Lipid Identification Software | MS-DIAL |
| Isotope correction at MS2 | No | Data manipulation | Smoothing, Centroiding |
| MS1 verified by standard | No | Nomenclature for intact lipid molecule | Yes |
| MS2 verified by standard | No | Nomenclature for fragment ions | No |
| Background check at MS1 | Yes | Further identification remarks | - |

#### 28) Ceramide non-hydroxyfatty acid-phytospingosine (Cer_NP)[M+CH3COO]- / Lipid quantification

| | | | |
|---|---|---|---|
| Quantitative | Yes | Limit of quantification | No |
| MS Level for quantification | MS1 | Normalization to reference | No |
| Internal lipid standard(s) MS1 | | Lipid Quantification Software | MS-DIAL |

| Internal standard | Endogenous subclass |
|---|---|
| Cer 18:1;2O/15:0(d7) | Cer_NP subclass |

| | | | |
|---|---|---|---|
| Type of quantification | Internal standard amount | Batch correction | No |
| Response correction | No | Further quantification remarks | - |
| Type I isotope correction | No | | |

#### 29) Ceramide non-hydroxyfatty acid-sphingosine (Cer_NS)[M+CH3COO]- / Lipid identification

| | | | |
|---|---|---|---|
| Lipid class | Ceramide non-hydroxyfatty acid-sphingosine (Cer_NS) | Background check at MS2 | No |
| Derivatization | - | Did you presume assumptions for identification? | No |
| MS Level for identification | MS1, MS2 | Check isomer overlap | No |
| Identification level | Molecular species level | RT verified by standard | Yes |
| Polarity mode | Negative | Separation of isobaric/isomeric interferece confirmed | Yes |
| Type of negative (precursor)ion | [M+CH3COO]- | Model for separation prediction | Yes |
| Fragments for identification | | Additional dimension/techniques | - |

| Fragment name |
|---|
| Neutral loss of H2O |
| Neutral loss of CH2O |
| Sphingosine -C2H7NO fragment |
| Fatty acyl +C2H3N fragment |
| Fatty acyl -2H fragment |

| | | | |
|---|---|---|---|
| Isotope correction at MS1 | No | Lipid Identification Software | MS-DIAL |
| Isotope correction at MS2 | No | Data manipulation | Smoothing, Centroiding |
| MS1 verified by standard | Yes | Nomenclature for intact lipid molecule | Yes |
| MS2 verified by standard | Yes | Nomenclature for fragment ions | No |
| Background check at MS1 | Yes | Further identification remarks | - |

#### 29) Ceramide non-hydroxyfatty acid-sphingosine (Cer_NS)[M+CH3COO]- / Lipid quantification

| | | | |
|---|---|---|---|
| Quantitative | Yes | Limit of quantification | No |
| MS Level for quantification | MS1 | Normalization to reference | No |
| Internal lipid standard(s) MS1 | | Lipid Quantification Software | MS-DIAL |

| Internal standard | Endogenous subclass |
|---|---|
| Cer 18:1;2O/15:0(d7) | Cer_NS subclass |

| | | | |
|---|---|---|---|
| Type of quantification | Internal standard amount | Batch correction | No |
| Response correction | No | Further quantification remarks | - |
| Type I isotope correction | No | | |

#### 30) Ceramide phosphoethanolamine (PE_Cer)[M-H]- / Lipid identification

| | | | |
|---|---|---|---|
| Lipid class | Ceramide phosphoethanolamine (PE_Cer) | Background check at MS2 | No |
| Derivatization | - | Did you presume assumptions for identification? | No |
| MS Level for identification | MS1, MS2 | Check isomer overlap | No |
| Identification level | Molecular species level | RT verified by standard | Yes |
| Polarity mode | Negative | Separation of isobaric/isomeric interferece confirmed | Yes |
| Type of negative (precursor)ion | [M-H]- | Model for separation prediction | Yes |
| Fragments for identification | | Additional dimension/techniques | - |

| Fragment name |
|---|
| Characteristic fragment (C2H7NO4P-) |
| Neutral loss of fatty acyl |

| | | | |
|---|---|---|---|
| Isotope correction at MS1 | No | Lipid Identification Software | MS-DIAL |
| Isotope correction at MS2 | No | Data manipulation | Smoothing, Centroiding |
| MS1 verified by standard | No | Nomenclature for intact lipid molecule | Yes |
| MS2 verified by standard | No | Nomenclature for fragment ions | No |
| Background check at MS1 | Yes | Further identification remarks | - |

#### 30) Ceramide phosphoethanolamine (PE_Cer)[M-H]- / Lipid quantification

| | | | |
|---|---|---|---|
| Quantitative | Yes | Limit of quantification | No |
| MS Level for quantification | MS1 | Normalization to reference | No |
| Internal lipid standard(s) MS1 | | Lipid Quantification Software | MS-DIAL |

| Internal standard | Endogenous subclass |
|---|---|
| Cer 18:1;2O/15:0(d7) | PE_Cer subclass |

| | | | |
|---|---|---|---|
| Type of quantification | Internal standard amount | Batch correction | No |
| Response correction | No | Further quantification remarks | - |
| Type I isotope correction | No | | |

#### 31) Ceramide phosphoinositol (PI_Cer)[M-H]- / Lipid identification

| | | | |
|---|---|---|---|
| Lipid class | Ceramide phosphoinositol (PI_Cer) | Background check at MS2 | No |
| Derivatization | - | Did you presume assumptions for identification? | No |
| MS Level for identification | MS1, MS2 | Check isomer overlap | No |
| Identification level | Molecular species level | RT verified by standard | Yes |
| Polarity mode | Negative | Separation of isobaric/isomeric interferece confirmed | Yes |
| Type of negative (precursor)ion | [M-H]- | Model for separation prediction | Yes |
| Fragments for identification | | Additional dimension/techniques | - |

| Fragment name |
|---|
| Phosphoinositol - H2O |
| Neutral loss of inositol |
| Neutral loss of fatty acyl |

| | | | |
|---|---|---|---|
| Isotope correction at MS1 | No | Lipid Identification Software | MS-DIAL |
| Isotope correction at MS2 | No | Data manipulation | Smoothing, Centroiding |
| MS1 verified by standard | No | Nomenclature for intact lipid molecule | Yes |
| MS2 verified by standard | No | Nomenclature for fragment ions | No |
| Background check at MS1 | Yes | Further identification remarks | - |

#### 31) Ceramide phosphoinositol (PI_Cer)[M-H]- / Lipid quantification

| | | | |
|---|---|---|---|
| Quantitative | Yes | Limit of quantification | No |
| MS Level for quantification | MS1 | Normalization to reference | No |
| Internal lipid standard(s) MS1 | | Lipid Quantification Software | MS-DIAL |

| Internal standard | Endogenous subclass |
|---|---|
| Cer 18:1;2O/15:0(d7) | PI_Cer subclass |

| | | | |
|---|---|---|---|
| Type of quantification | Internal standard amount | Batch correction | No |
| Response correction | No | Further quantification remarks | - |
| Type I isotope correction | No | | |

#### 32) FA[M+NH4]+ / Lipid identification

| | | | |
|---|---|---|---|
| Lipid class | FA | Background check at MS2 | No |
| Derivatization | - | Did you presume assumptions for identification? | No |
| MS Level for identification | MS1, MS2 | Check isomer overlap | No |
| Identification level | Molecular species level | RT verified by standard | Yes |
| Polarity mode | Positive | Separation of isobaric/isomeric interferece confirmed | Yes |
| Type of positive (precursor)ion | [M+NH4]+ | Model for separation prediction | Yes |
| Fragments for identification | | Additional dimension/techniques | - |

| Fragment name |
|---|
| Neutral loss of fatty acyl |

| | | | |
|---|---|---|---|
| Isotope correction at MS1 | No | Lipid Identification Software | MS-DIAL |
| Isotope correction at MS2 | No | Data manipulation | Smoothing, Centroiding |
| MS1 verified by standard | Yes | Nomenclature for intact lipid molecule | Yes |
| MS2 verified by standard | Yes | Nomenclature for fragment ions | No |
| Background check at MS1 | Yes | Further identification remarks | - |

#### 32) FA[M+NH4]+ / Lipid quantification

| | | | |
|---|---|---|---|
| Quantitative | Yes | Limit of quantification | No |
| MS Level for quantification | MS1 | Normalization to reference | No |
| Internal lipid standard(s) MS1 | | Lipid Quantification Software | MS-DIAL |

| Internal standard | Endogenous subclass |
|---|---|
| CE 18:1(d7) | CE subclass |

| | | | |
|---|---|---|---|
| Type of quantification | Internal standard amount | Batch correction | No |
| Response correction | No | Further quantification remarks | - |
| Type I isotope correction | No | | |

#### 33) Cholic acid (BileAcid)[M-H]- / Lipid identification

| | | | |
|---|---|---|---|
| Lipid class | Cholic acid (BileAcid) | Check isomer overlap | No |
| Derivatization | - | RT verified by standard | Yes |
| MS Level for identification | MS1 | Separation of isobaric/isomeric interferece confirmed | Yes |
| Identification level | Species level | Model for separation prediction | Yes |
| Polarity mode | Negative | Additional dimension/techniques | - |
| Type of negative (precursor)ion | [M-H]- | Lipid Identification Software | MS-DIAL |
| Isotope correction at MS1 | No | Data manipulation | Smoothing, Centroiding |
| MS1 verified by standard | No | Nomenclature for intact lipid molecule | Yes |
| Background check at MS1 | Yes | Further identification remarks | - |
| Did you presume assumptions for identification? | No | | |

#### 33) Cholic acid (BileAcid)[M-H]- / Lipid quantification

| | | | |
|---|---|---|---|
| Quantitative | Yes | Limit of quantification | No |
| MS Level for quantification | MS1 | Normalization to reference | No |
| Internal lipid standard(s) MS1 | | Lipid Quantification Software | MS-DIAL |

| Internal standard | Endogenous subclass |
|---|---|
| LPC 18:1(d7) | BileAcid subclass |

| | | | |
|---|---|---|---|
| Type of quantification | Internal standard amount | Batch correction | No |
| Response correction | No | Further quantification remarks | - |
| Type I isotope correction | No | | |

#### 34) Cholic acid sulfate (BASulfate)[M-H]- / Lipid identification

| | | | |
|---|---|---|---|
| Lipid class | Cholic acid sulfate (BASulfate) | Background check at MS2 | No |
| Derivatization | - | Did you presume assumptions for identification? | No |
| MS Level for identification | MS1, MS2 | Check isomer overlap | No |
| Identification level | Molecular species level | RT verified by standard | Yes |
| Polarity mode | Negative | Separation of isobaric/isomeric interferece confirmed | Yes |
| Type of negative (precursor)ion | [M-H]- | Model for separation prediction | Yes |
| Fragments for identification | | Additional dimension/techniques | - |

| Fragment name |
|---|
| Hydrogensulfate |

| | | | |
|---|---|---|---|
| Isotope correction at MS1 | No | Lipid Identification Software | MS-DIAL |
| Isotope correction at MS2 | No | Data manipulation | Smoothing, Centroiding |
| MS1 verified by standard | No | Nomenclature for intact lipid molecule | Yes |
| MS2 verified by standard | No | Nomenclature for fragment ions | No |
| Background check at MS1 | Yes | Further identification remarks | - |

#### 34) Cholic acid sulfate (BASulfate)[M-H]- / Lipid quantification

| | | | |
|---|---|---|---|
| Quantitative | Yes | Limit of quantification | No |
| MS Level for quantification | MS1 | Normalization to reference | No |
| Internal lipid standard(s) MS1 | | Lipid Quantification Software | MS-DIAL |

| Internal standard | Endogenous subclass |
|---|---|
| LPC 18:1(d7) | BASulfate subclass |

| | | | |
|---|---|---|---|
| Type of quantification | Internal standard amount | Batch correction | No |
| Response correction | No | Further quantification remarks | - |
| Type I isotope correction | No | | |

#### 35) Coenzyme Q (CoQ)[M+H]+ / Lipid identification

| | | | |
|---|---|---|---|
| Lipid class | Coenzyme Q (CoQ) | Background check at MS2 | No |
| Derivatization | - | Did you presume assumptions for identification? | No |
| MS Level for identification | MS1, MS2 | Check isomer overlap | No |
| Identification level | Molecular species level | RT verified by standard | Yes |
| Polarity mode | Positive | Separation of isobaric/isomeric interferece confirmed | Yes |
| Type of positive (precursor)ion | [M+H]+ | Model for separation prediction | Yes |
| Fragments for identification | | Additional dimension/techniques | - |

| Fragment name |
|---|
| Characteristic fragment (C10H13O4+) |

| | | | |
|---|---|---|---|
| Isotope correction at MS1 | No | Lipid Identification Software | MS-DIAL |
| Isotope correction at MS2 | No | Data manipulation | Smoothing, Centroiding |
| MS1 verified by standard | No | Nomenclature for intact lipid molecule | Yes |
| MS2 verified by standard | No | Nomenclature for fragment ions | No |
| Background check at MS1 | Yes | Further identification remarks | - |

#### 35) Coenzyme Q (CoQ)[M+H]+ / Lipid quantification

| | | | |
|---|---|---|---|
| Quantitative | Yes | Limit of quantification | No |
| MS Level for quantification | MS1 | Normalization to reference | No |
| Internal lipid standard(s) MS1 | | Lipid Quantification Software | MS-DIAL |

| Internal standard | Endogenous subclass |
|---|---|
| LPC 18:1(d7) | CoQ subclass |

| | | | |
|---|---|---|---|
| Type of quantification | Internal standard amount | Batch correction | No |
| Response correction | No | Further quantification remarks | - |
| Type I isotope correction | No | | |

#### 36) Dehydroergosterol ester (DEGSE)[M+NH4]+ / Lipid identification

| | | | |
|---|---|---|---|
| Lipid class | Dehydroergosterol ester (DEGSE) | Background check at MS2 | No |
| Derivatization | - | Did you presume assumptions for identification? | No |
| MS Level for identification | MS1, MS2 | Check isomer overlap | No |
| Identification level | Molecular species level | RT verified by standard | Yes |
| Polarity mode | Positive | Separation of isobaric/isomeric interferece confirmed | Yes |
| Type of positive (precursor)ion | [M+NH4]+ | Model for separation prediction | Yes |
| Fragments for identification | | Additional dimension/techniques | - |

| Fragment name |
|---|
| Neutral loss of fatty acyl |

| | | | |
|---|---|---|---|
| Isotope correction at MS1 | No | Lipid Identification Software | MS-DIAL |
| Isotope correction at MS2 | No | Data manipulation | Smoothing, Centroiding |
| MS1 verified by standard | No | Nomenclature for intact lipid molecule | Yes |
| MS2 verified by standard | No | Nomenclature for fragment ions | No |
| Background check at MS1 | Yes | Further identification remarks | - |

#### 36) Dehydroergosterol ester (DEGSE)[M+NH4]+ / Lipid quantification

| | | | |
|---|---|---|---|
| Quantitative | Yes | Limit of quantification | No |
| MS Level for quantification | MS1 | Normalization to reference | No |
| Internal lipid standard(s) MS1 | | Lipid Quantification Software | MS-DIAL |

| Internal standard | Endogenous subclass |
|---|---|
| CE 18:1(d7) | DEGSE subclass |

| | | | |
|---|---|---|---|
| Type of quantification | Internal standard amount | Batch correction | No |
| Response correction | No | Further quantification remarks | - |
| Type I isotope correction | No | | |

#### 37) Desmosterol ester (DSMSE)[M+NH4]+ / Lipid identification

| | | | |
|---|---|---|---|
| Lipid class | Desmosterol ester (DSMSE) | Background check at MS2 | No |
| Derivatization | - | Did you presume assumptions for identification? | No |
| MS Level for identification | MS1, MS2 | Check isomer overlap | No |
| Identification level | Molecular species level | RT verified by standard | Yes |
| Polarity mode | Positive | Separation of isobaric/isomeric interferece confirmed | Yes |
| Type of positive (precursor)ion | [M+NH4]+ | Model for separation prediction | Yes |
| Fragments for identification | | Additional dimension/techniques | - |

| Fragment name |
|---|
| Neutral loss of fatty acyl |

| | | | |
|---|---|---|---|
| Isotope correction at MS1 | No | Lipid Identification Software | MS-DIAL |
| Isotope correction at MS2 | No | Data manipulation | Smoothing, Centroiding |
| MS1 verified by standard | No | Nomenclature for intact lipid molecule | Yes |
| MS2 verified by standard | No | Nomenclature for fragment ions | No |
| Background check at MS1 | Yes | Further identification remarks | - |

#### 37) Desmosterol ester (DSMSE)[M+NH4]+ / Lipid quantification

| | | | |
|---|---|---|---|
| Quantitative | Yes | Limit of quantification | No |
| MS Level for quantification | MS1 | Normalization to reference | No |
| Internal lipid standard(s) MS1 | | Lipid Quantification Software | MS-DIAL |

| Internal standard | Endogenous subclass |
|---|---|
| CE 18:1(d7) | DSMSE subclass |

| | | | |
|---|---|---|---|
| Type of quantification | Internal standard amount | Batch correction | No |
| Response correction | No | Further quantification remarks | - |
| Type I isotope correction | No | | |

#### 38) Diacylglyceryl glucuronide (DGGA)[M-H]- / Lipid identification

| | | | |
|---|---|---|---|
| Lipid class | Diacylglyceryl glucuronide (DGGA) | Background check at MS2 | No |
| Derivatization | - | Did you presume assumptions for identification? | No |
| MS Level for identification | MS1, MS2 | Check isomer overlap | No |
| Identification level | Molecular species level | RT verified by standard | Yes |
| Polarity mode | Negative | Separation of isobaric/isomeric interferece confirmed | Yes |
| Type of negative (precursor)ion | [M-H]- | Model for separation prediction | Yes |
| Fragments for identification | | Additional dimension/techniques | - |

| Fragment name |
|---|
| Fatty acid fragment |

| | | | |
|---|---|---|---|
| Isotope correction at MS1 | No | Lipid Identification Software | MS-DIAL |
| Isotope correction at MS2 | No | Data manipulation | Smoothing, Centroiding |
| MS1 verified by standard | No | Nomenclature for intact lipid molecule | Yes |
| MS2 verified by standard | No | Nomenclature for fragment ions | No |
| Background check at MS1 | Yes | Further identification remarks | - |

#### 38) Diacylglyceryl glucuronide (DGGA)[M-H]- / Lipid quantification

| | | | |
|---|---|---|---|
| Quantitative | Yes | Limit of quantification | No |
| MS Level for quantification | MS1 | Normalization to reference | No |
| Internal lipid standard(s) MS1 | | Lipid Quantification Software | MS-DIAL |

| Internal standard | Endogenous subclass |
|---|---|
| LPC 18:1(d7) | DGGA subclass |

| | | | |
|---|---|---|---|
| Type of quantification | Internal standard amount | Batch correction | No |
| Response correction | No | Further quantification remarks | - |
| Type I isotope correction | No | | |

#### 39) Diacylglyceryl trimethylhomoserine (DGTS)[M+H]+ / Lipid identification

| | | | |
|---|---|---|---|
| Lipid class | Diacylglyceryl trimethylhomoserine (DGTS) | Background check at MS2 | No |
| Derivatization | - | Did you presume assumptions for identification? | No |
| MS Level for identification | MS1, MS2 | Check isomer overlap | No |
| Identification level | Molecular species level | RT verified by standard | Yes |
| Polarity mode | Positive | Separation of isobaric/isomeric interferece confirmed | Yes |
| Type of positive (precursor)ion | [M+H]+ | Model for separation prediction | Yes |
| Fragments for identification | | Additional dimension/techniques | - |

| Fragment name |
|---|
| Characteristic fragment (C10H22NO5+) |
| Characteristic fragment (C7H14NO2+) |
| Neutral loss of fatty acyl |
| Neutral loss of fatty acyl and H2O |

| | | | |
|---|---|---|---|
| Isotope correction at MS1 | No | Lipid Identification Software | MS-DIAL |
| Isotope correction at MS2 | No | Data manipulation | Smoothing, Centroiding |
| MS1 verified by standard | No | Nomenclature for intact lipid molecule | Yes |
| MS2 verified by standard | No | Nomenclature for fragment ions | No |
| Background check at MS1 | Yes | Further identification remarks | - |

#### 39) Diacylglyceryl trimethylhomoserine (DGTS)[M+H]+ / Lipid quantification

| | | | |
|---|---|---|---|
| Quantitative | Yes | Limit of quantification | No |
| MS Level for quantification | MS1 | Normalization to reference | No |
| Internal lipid standard(s) MS1 | | Lipid Quantification Software | MS-DIAL |

| Internal standard | Endogenous subclass |
|---|---|
| LPC 18:1(d7) | DGTS subclass |

| | | | |
|---|---|---|---|
| Type of quantification | Internal standard amount | Batch correction | No |
| Response correction | No | Further quantification remarks | - |
| Type I isotope correction | No | | |

#### 40) Diacylglyceryl-3-O-carboxyhydroxymethylcholine (DGCC)[M+H]+ / Lipid identification

| | | | |
|---|---|---|---|
| Lipid class | Diacylglyceryl-3-O-carboxyhydroxymethylcholine (DGCC) | Background check at MS2 | No |
| Derivatization | - | Did you presume assumptions for identification? | No |
| MS Level for identification | MS1, MS2 | Check isomer overlap | No |
| Identification level | Molecular species level | RT verified by standard | Yes |
| Polarity mode | Positive | Separation of isobaric/isomeric interferece confirmed | Yes |
| Type of positive (precursor)ion | [M+H]+ | Model for separation prediction | Yes |
| Fragments for identification | | Additional dimension/techniques | - |

| Fragment name |
|---|
| Characteristic fragment (C6H14NO2+) |
| Neutral loss of fatty acyl |
| Neutral loss of fatty acyl and H2O |

| | | | |
|---|---|---|---|
| Isotope correction at MS1 | No | Lipid Identification Software | MS-DIAL |
| Isotope correction at MS2 | No | Data manipulation | Smoothing, Centroiding |
| MS1 verified by standard | No | Nomenclature for intact lipid molecule | Yes |
| MS2 verified by standard | No | Nomenclature for fragment ions | No |
| Background check at MS1 | Yes | Further identification remarks | - |

#### 40) Diacylglyceryl-3-O-carboxyhydroxymethylcholine (DGCC)[M+H]+ / Lipid quantification

| | | | |
|---|---|---|---|
| Quantitative | Yes | Limit of quantification | No |
| MS Level for quantification | MS1 | Normalization to reference | No |
| Internal lipid standard(s) MS1 | | Lipid Quantification Software | MS-DIAL |

| Internal standard | Endogenous subclass |
|---|---|
| LPC 18:1(d7) | DGCC subclass |

| | | | |
|---|---|---|---|
| Type of quantification | Internal standard amount | Batch correction | No |
| Response correction | No | Further quantification remarks | - |
| Type I isotope correction | No | | |

#### 41) Digalactosylmonoacylglycerol (DGMG)[M+CH3COO]- / Lipid identification

| | | | |
|---|---|---|---|
| Lipid class | Digalactosylmonoacylglycerol (DGMG) | Background check at MS2 | No |
| Derivatization | - | Did you presume assumptions for identification? | No |
| MS Level for identification | MS1, MS2 | Check isomer overlap | No |
| Identification level | Molecular species level | RT verified by standard | Yes |
| Polarity mode | Negative | Separation of isobaric/isomeric interferece confirmed | Yes |
| Type of negative (precursor)ion | [M+CH3COO]- | Model for separation prediction | Yes |
| Fragments for identification | | Additional dimension/techniques | - |

| Fragment name |
|---|
| Fatty acid fragment |

| | | | |
|---|---|---|---|
| Isotope correction at MS1 | No | Lipid Identification Software | MS-DIAL |
| Isotope correction at MS2 | No | Data manipulation | Smoothing, Centroiding |
| MS1 verified by standard | No | Nomenclature for intact lipid molecule | Yes |
| MS2 verified by standard | No | Nomenclature for fragment ions | No |
| Background check at MS1 | Yes | Further identification remarks | - |

#### 41) Digalactosylmonoacylglycerol (DGMG)[M+CH3COO]- / Lipid quantification

| | | | |
|---|---|---|---|
| Quantitative | Yes | Limit of quantification | No |
| MS Level for quantification | MS1 | Normalization to reference | No |
| Internal lipid standard(s) MS1 | | Lipid Quantification Software | MS-DIAL |

| Internal standard | Endogenous subclass |
|---|---|
| LPC 18:1(d7) | DGMG subclass |

| | | | |
|---|---|---|---|
| Type of quantification | Internal standard amount | Batch correction | No |
| Response correction | No | Further quantification remarks | - |
| Type I isotope correction | No | | |

#### 42) Hex2Cer[M+H]+ / Lipid identification

| | | | |
|---|---|---|---|
| Lipid class | Hex2Cer | Background check at MS2 | No |
| Derivatization | - | Did you presume assumptions for identification? | No |
| MS Level for identification | MS1, MS2 | Check isomer overlap | No |
| Identification level | Molecular species level | RT verified by standard | Yes |
| Polarity mode | Positive | Separation of isobaric/isomeric interferece confirmed | Yes |
| Type of positive (precursor)ion | [M+H]+ | Model for separation prediction | Yes |
| Fragments for identification | | Additional dimension/techniques | - |

| Fragment name |
|---|
| Neutral loss of hexose |
| Neutral loss of 2hexose |
| Sphingosine -H2O fragment |
| Sphingosine -2H2O fragment |
| Sphingosine -CH4O2 fragment |

| | | | |
|---|---|---|---|
| Isotope correction at MS1 | No | Lipid Identification Software | MS-DIAL |
| Isotope correction at MS2 | No | Data manipulation | Smoothing, Centroiding |
| MS1 verified by standard | No | Nomenclature for intact lipid molecule | Yes |
| MS2 verified by standard | No | Nomenclature for fragment ions | No |
| Background check at MS1 | Yes | Further identification remarks | - |

#### 42) Hex2Cer[M+H]+ / Lipid quantification

| | | | |
|---|---|---|---|
| Quantitative | Yes | Limit of quantification | No |
| MS Level for quantification | MS1 | Normalization to reference | No |
| Internal lipid standard(s) MS1 | | Lipid Quantification Software | MS-DIAL |

| Internal standard | Endogenous subclass |
|---|---|
| Cer 18:1;2O/15:0(d7) | Hex2Cer subclass |

| | | | |
|---|---|---|---|
| Type of quantification | Internal standard amount | Batch correction | No |
| Response correction | No | Further quantification remarks | - |
| Type I isotope correction | No | | |

#### 43) Dilysocardiolipin (DLCL)[M-H]- / Lipid identification

| | | | |
|---|---|---|---|
| Lipid class | Dilysocardiolipin (DLCL) | Background check at MS2 | No |
| Derivatization | - | Did you presume assumptions for identification? | No |
| MS Level for identification | MS1, MS2 | Check isomer overlap | No |
| Identification level | Molecular species level | RT verified by standard | Yes |
| Polarity mode | Negative | Separation of isobaric/isomeric interferece confirmed | Yes |
| Type of negative (precursor)ion | [M-H]- | Model for separation prediction | Yes |
| Fragments for identification | | Additional dimension/techniques | - |

| Fragment name |
|---|
| Phosphoglycerol -H2O fragment |
| lysophosphatidic acid |

| | | | |
|---|---|---|---|
| Isotope correction at MS1 | No | Lipid Identification Software | MS-DIAL |
| Isotope correction at MS2 | No | Data manipulation | Smoothing, Centroiding |
| MS1 verified by standard | No | Nomenclature for intact lipid molecule | Yes |
| MS2 verified by standard | No | Nomenclature for fragment ions | No |
| Background check at MS1 | Yes | Further identification remarks | - |

#### 43) Dilysocardiolipin (DLCL)[M-H]- / Lipid quantification

| | | | |
|---|---|---|---|
| Quantitative | Yes | Limit of quantification | No |
| MS Level for quantification | MS1 | Normalization to reference | No |
| Internal lipid standard(s) MS1 | | Lipid Quantification Software | MS-DIAL |

| Internal standard | Endogenous subclass |
|---|---|
| PG 15:0_18:1(d7) | DLCL subclass |

| | | | |
|---|---|---|---|
| Type of quantification | Internal standard amount | Batch correction | No |
| Response correction | No | Further quantification remarks | - |
| Type I isotope correction | No | | |

#### 44) Ergosterol ester (EGSE)[M+NH4]+ / Lipid identification

| | | | |
|---|---|---|---|
| Lipid class | Ergosterol ester (EGSE) | Background check at MS2 | No |
| Derivatization | - | Did you presume assumptions for identification? | No |
| MS Level for identification | MS1, MS2 | Check isomer overlap | No |
| Identification level | Molecular species level | RT verified by standard | Yes |
| Polarity mode | Positive | Separation of isobaric/isomeric interferece confirmed | Yes |
| Type of positive (precursor)ion | [M+NH4]+ | Model for separation prediction | Yes |
| Fragments for identification | | Additional dimension/techniques | - |

| Fragment name |
|---|
| Neutral loss of fatty acyl |

| | | | |
|---|---|---|---|
| Isotope correction at MS1 | No | Lipid Identification Software | MS-DIAL |
| Isotope correction at MS2 | No | Data manipulation | Smoothing, Centroiding |
| MS1 verified by standard | No | Nomenclature for intact lipid molecule | Yes |
| MS2 verified by standard | No | Nomenclature for fragment ions | No |
| Background check at MS1 | Yes | Further identification remarks | - |

#### 44) Ergosterol ester (EGSE)[M+NH4]+ / Lipid quantification

| | | | |
|---|---|---|---|
| Quantitative | Yes | Limit of quantification | No |
| MS Level for quantification | MS1 | Normalization to reference | No |
| Internal lipid standard(s) MS1 | | Lipid Quantification Software | MS-DIAL |

| Internal standard | Endogenous subclass |
|---|---|
| CE 18:1(d7) | EGSE subclass |

| | | | |
|---|---|---|---|
| Type of quantification | Internal standard amount | Batch correction | No |
| Response correction | No | Further quantification remarks | - |
| Type I isotope correction | No | | |

#### 45) Esterified ketodeoxycholic acid (KDCAE)[M+NH4]+ / Lipid identification

| | | | |
|---|---|---|---|
| Lipid class | Esterified ketodeoxycholic acid (KDCAE) | Background check at MS2 | No |
| Derivatization | - | Did you presume assumptions for identification? | No |
| MS Level for identification | MS1, MS2 | Check isomer overlap | No |
| Identification level | Molecular species level | RT verified by standard | Yes |
| Polarity mode | Positive | Separation of isobaric/isomeric interferece confirmed | Yes |
| Type of positive (precursor)ion | [M+NH4]+ | Model for separation prediction | Yes |
| Fragments for identification | | Additional dimension/techniques | - |

| Fragment name |
|---|
| Neutral loss of fatty acyl |

| | | | |
|---|---|---|---|
| Isotope correction at MS1 | No | Lipid Identification Software | MS-DIAL |
| Isotope correction at MS2 | No | Data manipulation | Smoothing, Centroiding |
| MS1 verified by standard | No | Nomenclature for intact lipid molecule | Yes |
| MS2 verified by standard | No | Nomenclature for fragment ions | No |
| Background check at MS1 | Yes | Further identification remarks | - |

#### 45) Esterified ketodeoxycholic acid (KDCAE)[M+NH4]+ / Lipid quantification

| | | | |
|---|---|---|---|
| Quantitative | Yes | Limit of quantification | No |
| MS Level for quantification | MS1 | Normalization to reference | No |
| Internal lipid standard(s) MS1 | | Lipid Quantification Software | MS-DIAL |

| Internal standard | Endogenous subclass |
|---|---|
| CE 18:1(d7) | KDCAE subclass |

| | | | |
|---|---|---|---|
| Type of quantification | Internal standard amount | Batch correction | No |
| Response correction | No | Further quantification remarks | - |
| Type I isotope correction | No | | |

#### 46) Esterified taurodeoxycholic Acid (TDCAE)[M+NH4]+ / Lipid identification

| | | | |
|---|---|---|---|
| Lipid class | Esterified taurodeoxycholic Acid (TDCAE) | Background check at MS2 | No |
| Derivatization | - | Did you presume assumptions for identification? | No |
| MS Level for identification | MS1, MS2 | Check isomer overlap | No |
| Identification level | Molecular species level | RT verified by standard | Yes |
| Polarity mode | Positive | Separation of isobaric/isomeric interferece confirmed | Yes |
| Type of positive (precursor)ion | [M+NH4]+ | Model for separation prediction | Yes |
| Fragments for identification | | Additional dimension/techniques | - |

| Fragment name |
|---|
| Neutral loss of fatty acyl |

| | | | |
|---|---|---|---|
| Isotope correction at MS1 | No | Lipid Identification Software | MS-DIAL |
| Isotope correction at MS2 | No | Data manipulation | Smoothing, Centroiding |
| MS1 verified by standard | No | Nomenclature for intact lipid molecule | Yes |
| MS2 verified by standard | No | Nomenclature for fragment ions | No |
| Background check at MS1 | Yes | Further identification remarks | - |

#### 46) Esterified taurodeoxycholic Acid (TDCAE)[M+NH4]+ / Lipid quantification

| | | | |
|---|---|---|---|
| Quantitative | Yes | Limit of quantification | No |
| MS Level for quantification | MS1 | Normalization to reference | No |
| Internal lipid standard(s) MS1 | | Lipid Quantification Software | MS-DIAL |

| Internal standard | Endogenous subclass |
|---|---|
| CE 18:1(d7) | TDCAE subclass |

| | | | |
|---|---|---|---|
| Type of quantification | Internal standard amount | Batch correction | No |
| Response correction | No | Further quantification remarks | - |
| Type I isotope correction | No | | |

#### 47) Esterified deoxycholic acid (DCAE)[M+NH4]+ / Lipid identification

| | | | |
|---|---|---|---|
| Lipid class | Esterified deoxycholic acid (DCAE) | Background check at MS2 | No |
| Derivatization | - | Did you presume assumptions for identification? | No |
| MS Level for identification | MS1, MS2 | Check isomer overlap | No |
| Identification level | Molecular species level | RT verified by standard | Yes |
| Polarity mode | Positive | Separation of isobaric/isomeric interferece confirmed | Yes |
| Type of positive (precursor)ion | [M+NH4]+ | Model for separation prediction | Yes |
| Fragments for identification | | Additional dimension/techniques | - |

| Fragment name |
|---|
| Neutral loss of fatty acyl |

| | | | |
|---|---|---|---|
| Isotope correction at MS1 | No | Lipid Identification Software | MS-DIAL |
| Isotope correction at MS2 | No | Data manipulation | Smoothing, Centroiding |
| MS1 verified by standard | No | Nomenclature for intact lipid molecule | Yes |
| MS2 verified by standard | No | Nomenclature for fragment ions | No |
| Background check at MS1 | Yes | Further identification remarks | - |

#### 47) Esterified deoxycholic acid (DCAE)[M+NH4]+ / Lipid quantification

| | | | |
|---|---|---|---|
| Quantitative | Yes | Limit of quantification | No |
| MS Level for quantification | MS1 | Normalization to reference | No |
| Internal lipid standard(s) MS1 | | Lipid Quantification Software | MS-DIAL |

| Internal standard | Endogenous subclass |
|---|---|
| CE 18:1(d7) | DCAE subclass |

| | | | |
|---|---|---|---|
| Type of quantification | Internal standard amount | Batch correction | No |
| Response correction | No | Further quantification remarks | - |
| Type I isotope correction | No | | |

#### 48) Esterified ketolithocholic acid (KLCAE)[M+NH4]+ / Lipid identification

| | | | |
|---|---|---|---|
| Lipid class | Esterified ketolithocholic acid (KLCAE) | Background check at MS2 | No |
| Derivatization | - | Did you presume assumptions for identification? | No |
| MS Level for identification | MS1, MS2 | Check isomer overlap | No |
| Identification level | Molecular species level | RT verified by standard | Yes |
| Polarity mode | Positive | Separation of isobaric/isomeric interferece confirmed | Yes |
| Type of positive (precursor)ion | [M+NH4]+ | Model for separation prediction | Yes |
| Fragments for identification | | Additional dimension/techniques | - |

| Fragment name |
|---|
| Neutral loss of fatty acyl |

| | | | |
|---|---|---|---|
| Isotope correction at MS1 | No | Lipid Identification Software | MS-DIAL |
| Isotope correction at MS2 | No | Data manipulation | Smoothing, Centroiding |
| MS1 verified by standard | No | Nomenclature for intact lipid molecule | Yes |
| MS2 verified by standard | No | Nomenclature for fragment ions | No |
| Background check at MS1 | Yes | Further identification remarks | - |

#### 48) Esterified ketolithocholic acid (KLCAE)[M+NH4]+ / Lipid quantification

| | | | |
|---|---|---|---|
| Quantitative | Yes | Limit of quantification | No |
| MS Level for quantification | MS1 | Normalization to reference | No |
| Internal lipid standard(s) MS1 | | Lipid Quantification Software | MS-DIAL |

| Internal standard | Endogenous subclass |
|---|---|
| CE 18:1(d7) | KLCAE subclass |

| | | | |
|---|---|---|---|
| Type of quantification | Internal standard amount | Batch correction | No |
| Response correction | No | Further quantification remarks | - |
| Type I isotope correction | No | | |

#### 49) Esterified lithocholic acid (LCAE)[M+NH4]+ / Lipid identification

| | | | |
|---|---|---|---|
| Lipid class | Esterified lithocholic acid (LCAE) | Background check at MS2 | No |
| Derivatization | - | Did you presume assumptions for identification? | No |
| MS Level for identification | MS1, MS2 | Check isomer overlap | No |
| Identification level | Molecular species level | RT verified by standard | Yes |
| Polarity mode | Positive | Separation of isobaric/isomeric interferece confirmed | Yes |
| Type of positive (precursor)ion | [M+NH4]+ | Model for separation prediction | Yes |
| Fragments for identification | | Additional dimension/techniques | - |

| Fragment name |
|---|
| Neutral loss of fatty acyl |

| | | | |
|---|---|---|---|
| Isotope correction at MS1 | No | Lipid Identification Software | MS-DIAL |
| Isotope correction at MS2 | No | Data manipulation | Smoothing, Centroiding |
| MS1 verified by standard | No | Nomenclature for intact lipid molecule | Yes |
| MS2 verified by standard | No | Nomenclature for fragment ions | No |
| Background check at MS1 | Yes | Further identification remarks | - |

#### 49) Esterified lithocholic acid (LCAE)[M+NH4]+ / Lipid quantification

| | | | |
|---|---|---|---|
| Quantitative | Yes | Limit of quantification | No |
| MS Level for quantification | MS1 | Normalization to reference | No |
| Internal lipid standard(s) MS1 | | Lipid Quantification Software | MS-DIAL |

| Internal standard | Endogenous subclass |
|---|---|
| CE 18:1(d7) | LCAE subclass |

| | | | |
|---|---|---|---|
| Type of quantification | Internal standard amount | Batch correction | No |
| Response correction | No | Further quantification remarks | - |
| Type I isotope correction | No | | |

#### 50) Ether-linked digalactosyldiacylglycerol (EtherDGDG)[M+CH3COO]- / Lipid identification

| | | | |
|---|---|---|---|
| Lipid class | Ether-linked digalactosyldiacylglycerol (EtherDGDG) | Background check at MS2 | No |
| Derivatization | - | Did you presume assumptions for identification? | No |
| MS Level for identification | MS1, MS2 | Check isomer overlap | No |
| Identification level | Molecular species level | RT verified by standard | Yes |
| Polarity mode | Negative | Separation of isobaric/isomeric interferece confirmed | Yes |
| Type of negative (precursor)ion | [M+CH3COO]- | Model for separation prediction | Yes |
| Fragments for identification | | Additional dimension/techniques | - |

| Fragment name |
|---|
| Neutral loss of fatty acyl |
| Fatty acid fragment |

| | | | |
|---|---|---|---|
| Isotope correction at MS1 | No | Lipid Identification Software | MS-DIAL |
| Isotope correction at MS2 | No | Data manipulation | Smoothing, Centroiding |
| MS1 verified by standard | No | Nomenclature for intact lipid molecule | Yes |
| MS2 verified by standard | No | Nomenclature for fragment ions | No |
| Background check at MS1 | Yes | Further identification remarks | - |

#### 50) Ether-linked digalactosyldiacylglycerol (EtherDGDG)[M+CH3COO]- / Lipid quantification

| | | | |
|---|---|---|---|
| Quantitative | Yes | Limit of quantification | No |
| MS Level for quantification | MS1 | Normalization to reference | No |
| Internal lipid standard(s) MS1 | | Lipid Quantification Software | MS-DIAL |

| Internal standard | Endogenous subclass |
|---|---|
| LPC 18:1(d7) | EtherDGDG subclass |

| | | | |
|---|---|---|---|
| Type of quantification | Internal standard amount | Batch correction | No |
| Response correction | No | Further quantification remarks | - |
| Type I isotope correction | No | | |

#### 51) LPC O[M+H]+ / Lipid identification

| | | | |
|---|---|---|---|
| Lipid class | LPC O | Background check at MS2 | No |
| Derivatization | - | Did you presume assumptions for identification? | No |
| MS Level for identification | MS1, MS2 | Check isomer overlap | No |
| Identification level | Molecular species level | RT verified by standard | Yes |
| Polarity mode | Positive | Separation of isobaric/isomeric interferece confirmed | Yes |
| Type of positive (precursor)ion | [M+H]+ | Model for separation prediction | Yes |
| Fragments for identification | | Additional dimension/techniques | - |

| Fragment name |
|---|
| Characteristic fragments (C5H14NO+) |
| Characteristic fragments (C2H6O4P+) |
| Characteristic fragments (C5H15NO4P+) |

| | | | |
|---|---|---|---|
| Isotope correction at MS1 | No | Lipid Identification Software | MS-DIAL |
| Isotope correction at MS2 | No | Data manipulation | Smoothing, Centroiding |
| MS1 verified by standard | No | Nomenclature for intact lipid molecule | Yes |
| MS2 verified by standard | No | Nomenclature for fragment ions | No |
| Background check at MS1 | Yes | Further identification remarks | - |

#### 51) LPC O[M+H]+ / Lipid quantification

| | | | |
|---|---|---|---|
| Quantitative | Yes | Limit of quantification | No |
| MS Level for quantification | MS1 | Normalization to reference | No |
| Internal lipid standard(s) MS1 | | Lipid Quantification Software | MS-DIAL |

| Internal standard | Endogenous subclass |
|---|---|
| LPC 18:1(d7) | EtherLPC subclass |

| | | | |
|---|---|---|---|
| Type of quantification | Internal standard amount | Batch correction | No |
| Response correction | No | Further quantification remarks | - |
| Type I isotope correction | No | | |

#### 52) LPE O[M+H]+ / Lipid identification

| | | | |
|---|---|---|---|
| Lipid class | LPE O | Background check at MS2 | No |
| Derivatization | - | Did you presume assumptions for identification? | No |
| MS Level for identification | MS1, MS2 | Check isomer overlap | No |
| Identification level | Molecular species level | RT verified by standard | Yes |
| Polarity mode | Positive | Separation of isobaric/isomeric interferece confirmed | Yes |
| Type of positive (precursor)ion | [M+H]+ | Model for separation prediction | Yes |
| Fragments for identification | | Additional dimension/techniques | - |

| Fragment name |
|---|
| Neutral loss of C3H8NO4P |
| Neutral loss of C3H10NO5P |

| | | | |
|---|---|---|---|
| Isotope correction at MS1 | No | Lipid Identification Software | MS-DIAL |
| Isotope correction at MS2 | No | Data manipulation | Smoothing, Centroiding |
| MS1 verified by standard | No | Nomenclature for intact lipid molecule | Yes |
| MS2 verified by standard | No | Nomenclature for fragment ions | No |
| Background check at MS1 | Yes | Further identification remarks | - |

#### 52) LPE O[M+H]+ / Lipid quantification

| | | | |
|---|---|---|---|
| Quantitative | Yes | Limit of quantification | No |
| MS Level for quantification | MS1 | Normalization to reference | No |
| Internal lipid standard(s) MS1 | | Lipid Quantification Software | MS-DIAL |

| Internal standard | Endogenous subclass |
|---|---|
| LPE 18:1(d7) | EtherLPE subclass |

| | | | |
|---|---|---|---|
| Type of quantification | Internal standard amount | Batch correction | No |
| Response correction | No | Further quantification remarks | - |
| Type I isotope correction | No | | |

#### 53) Ether-linked lysophosphatidylglycerol (EtherLPG)[M-H]- / Lipid identification

| | | | |
|---|---|---|---|
| Lipid class | Ether-linked lysophosphatidylglycerol (EtherLPG) | Background check at MS2 | No |
| Derivatization | - | Did you presume assumptions for identification? | No |
| MS Level for identification | MS1, MS2 | Check isomer overlap | No |
| Identification level | Molecular species level | RT verified by standard | Yes |
| Polarity mode | Negative | Separation of isobaric/isomeric interferece confirmed | Yes |
| Type of negative (precursor)ion | [M-H]- | Model for separation prediction | Yes |
| Fragments for identification | | Additional dimension/techniques | - |

| Fragment name |
|---|
| Phosphoglycerol -H2O fragment |
| Alkyl Ether fragment |

| | | | |
|---|---|---|---|
| Isotope correction at MS1 | No | Lipid Identification Software | MS-DIAL |
| Isotope correction at MS2 | No | Data manipulation | Smoothing, Centroiding |
| MS1 verified by standard | No | Nomenclature for intact lipid molecule | Yes |
| MS2 verified by standard | No | Nomenclature for fragment ions | No |
| Background check at MS1 | Yes | Further identification remarks | - |

#### 53) Ether-linked lysophosphatidylglycerol (EtherLPG)[M-H]- / Lipid quantification

| | | | |
|---|---|---|---|
| Quantitative | Yes | Limit of quantification | No |
| MS Level for quantification | MS1 | Normalization to reference | No |
| Internal lipid standard(s) MS1 | | Lipid Quantification Software | MS-DIAL |

| Internal standard | Endogenous subclass |
|---|---|
| PG 15:0_18:1(d7) | EtherLPG subclass |

| | | | |
|---|---|---|---|
| Type of quantification | Internal standard amount | Batch correction | No |
| Response correction | No | Further quantification remarks | - |
| Type I isotope correction | No | | |

#### 54) Ether-linked monogalactosyldiacylglycerol (EtherMGDG)[M+CH3COO]- / Lipid identification

| | | | |
|---|---|---|---|
| Lipid class | Ether-linked monogalactosyldiacylglycerol (EtherMGDG) | Background check at MS2 | No |
| Derivatization | - | Did you presume assumptions for identification? | No |
| MS Level for identification | MS1, MS2 | Check isomer overlap | No |
| Identification level | Molecular species level | RT verified by standard | Yes |
| Polarity mode | Negative | Separation of isobaric/isomeric interferece confirmed | Yes |
| Type of negative (precursor)ion | [M+CH3COO]- | Model for separation prediction | Yes |
| Fragments for identification | | Additional dimension/techniques | - |
| Isotope correction at MS1 | No | Lipid Identification Software | MS-DIAL |
| Isotope correction at MS2 | No | Data manipulation | Smoothing, Centroiding |
| MS1 verified by standard | No | Nomenclature for intact lipid molecule | Yes |
| MS2 verified by standard | No | Nomenclature for fragment ions | No |
| Background check at MS1 | Yes | Further identification remarks | - |

Fragments for identification:

| Fragment name |
|---|
| Neutral loss of fatty acyl |
| Fatty acid fragment |

#### 54) Ether-linked monogalactosyldiacylglycerol (EtherMGDG)[M+CH3COO]- / Lipid quantification

| | | | |
|---|---|---|---|
| Quantitative | Yes | Limit of quantification | No |
| MS Level for quantification | MS1 | Normalization to reference | No |
| Internal lipid standard(s) MS1 | | Lipid Quantification Software | MS-DIAL |
| Type of quantification | Internal standard amount | Batch correction | No |
| Response correction | No | Further quantification remarks | - |
| Type I isotope correction | No | | |

Internal lipid standard(s) MS1:

| Internal standard | Endogenous subclass |
|---|---|
| LPC 18:1(d7) | EtherMGDG subclass |

#### 55) Ether-linked oxidized phosphatidylcholine (EtherOxPC)[M+CH3COO]- / Lipid identification

| | | | |
|---|---|---|---|
| Lipid class | Ether-linked oxidized phosphatidylcholine (EtherOxPC) | Background check at MS2 | No |
| Derivatization | - | Did you presume assumptions for identification? | No |
| MS Level for identification | MS1, MS2 | Check isomer overlap | No |
| Identification level | Molecular species level | RT verified by standard | Yes |
| Polarity mode | Negative | Separation of isobaric/isomeric interferece confirmed | Yes |
| Type of negative (precursor)ion | [M+CH3COO]- | Model for separation prediction | Yes |
| Fragments for identification | | Additional dimension/techniques | - |

| Fragment name |
|---|
| Oxidized fatty acid fragment |
| Oxidized fatty acid -H2O fragment |
| Neutral loss of methyl moiety |

| | | | |
|---|---|---|---|
| Isotope correction at MS1 | No | Lipid Identification Software | MS-DIAL |
| Isotope correction at MS2 | No | Data manipulation | Smoothing, Centroiding |
| MS1 verified by standard | No | Nomenclature for intact lipid molecule | Yes |
| MS2 verified by standard | No | Nomenclature for fragment ions | No |
| Background check at MS1 | Yes | Further identification remarks | - |

#### 55) Ether-linked oxidized phosphatidylcholine (EtherOxPC)[M+CH3COO]- / Lipid quantification

| | | | |
|---|---|---|---|
| Quantitative | Yes | Limit of quantification | No |
| MS Level for quantification | MS1 | Normalization to reference | No |
| Internal lipid standard(s) MS1 | | Lipid Quantification Software | MS-DIAL |

| Internal standard | Endogenous subclass |
|---|---|
| PC 15:0_18:1(d7) | EtherOxPC subclass |

| | | | |
|---|---|---|---|
| Type of quantification | Internal standard amount | Batch correction | No |
| Response correction | No | Further quantification remarks | - |
| Type I isotope correction | No | | |

#### 56) Ether-linked oxidized phosphatidylethanolamine (EtherOxPE)[M-H]- / Lipid identification

| | | | |
|---|---|---|---|
| Lipid class | Ether-linked oxidized phosphatidylethanolamine (EtherOxPE) | Background check at MS2 | No |
| Derivatization | - | Did you presume assumptions for identification? | No |
| MS Level for identification | MS1, MS2 | Check isomer overlap | No |
| Identification level | Molecular species level | RT verified by standard | Yes |
| Polarity mode | Negative | Separation of isobaric/isomeric interferece confirmed | Yes |
| Type of negative (precursor)ion | [M-H]- | Model for separation prediction | Yes |
| Fragments for identification | | Additional dimension/techniques | - |

| Fragment name |
|---|
| Oxidized fatty acid fragment |
| Oxidized fatty acid -H2O fragment |

| | | | |
|---|---|---|---|
| Isotope correction at MS1 | No | Lipid Identification Software | MS-DIAL |
| Isotope correction at MS2 | No | Data manipulation | Smoothing, Centroiding |
| MS1 verified by standard | No | Nomenclature for intact lipid molecule | Yes |
| MS2 verified by standard | No | Nomenclature for fragment ions | No |
| Background check at MS1 | Yes | Further identification remarks | - |

#### 56) Ether-linked oxidized phosphatidylethanolamine (EtherOxPE)[M-H]- / Lipid quantification

| | | | |
|---|---|---|---|
| Quantitative | Yes | Limit of quantification | No |
| MS Level for quantification | MS1 | Normalization to reference | No |
| Internal lipid standard(s) MS1 | | Lipid Quantification Software | MS-DIAL |

| Internal standard | Endogenous subclass |
|---|---|
| PE 15:0_18:1(d7) | EtherOxPE subclass |

| | | | |
|---|---|---|---|
| Type of quantification | Internal standard amount | Batch correction | No |
| Response correction | No | Further quantification remarks | - |
| Type I isotope correction | No | | |

#### 57) PC O[M+CH3COO]- / Lipid identification

| | | | |
|---|---|---|---|
| Lipid class | PC O | Background check at MS2 | No |
| Derivatization | - | Did you presume assumptions for identification? | No |
| MS Level for identification | MS1, MS2 | Check isomer overlap | No |
| Identification level | Molecular species level | RT verified by standard | Yes |
| Polarity mode | Negative | Separation of isobaric/isomeric interferece confirmed | Yes |
| Type of negative (precursor)ion | [M+CH3COO]- | Model for separation prediction | Yes |
| Fragments for identification | | Additional dimension/techniques | - |

| Fragment name |
|---|
| Neutral loss of methyl moiety |
| Fatty acid fragment |

| | | | |
|---|---|---|---|
| Isotope correction at MS1 | No | Lipid Identification Software | MS-DIAL |
| Isotope correction at MS2 | No | Data manipulation | Smoothing, Centroiding |
| MS1 verified by standard | No | Nomenclature for intact lipid molecule | Yes |
| MS2 verified by standard | No | Nomenclature for fragment ions | No |
| Background check at MS1 | Yes | Further identification remarks | - |

#### 57) PC O[M+CH3COO]- / Lipid quantification

| | | | |
|---|---|---|---|
| Quantitative | Yes | Limit of quantification | No |
| MS Level for quantification | MS1 | Normalization to reference | No |
| Internal lipid standard(s) MS1 | | Lipid Quantification Software | MS-DIAL |

| Internal standard | Endogenous subclass |
|---|---|
| PC 15:0_18:1(d7) | EtherPC subclass |

| | | | |
|---|---|---|---|
| Type of quantification | Internal standard amount | Batch correction | No |
| Response correction | No | Further quantification remarks | - |
| Type I isotope correction | No | | |

#### 58) PE O[M-H]- / Lipid identification

| | | | |
|---|---|---|---|
| Lipid class | PE O | Background check at MS2 | No |
| Derivatization | - | Did you presume assumptions for identification? | No |
| MS Level for identification | MS1, MS2 | Check isomer overlap | No |
| Identification level | Molecular species level | RT verified by standard | Yes |
| Polarity mode | Negative | Separation of isobaric/isomeric interferece confirmed | Yes |
| Type of negative (precursor)ion | [M-H]- | Model for separation prediction | Yes |
| Fragments for identification | | Additional dimension/techniques | - |

| Fragment name |
|---|
| Neutral loss of fatty acyl |
| Fatty acid fragment |

| | | | |
|---|---|---|---|
| Isotope correction at MS1 | No | Lipid Identification Software | MS-DIAL |
| Isotope correction at MS2 | No | Data manipulation | Smoothing, Centroiding |
| MS1 verified by standard | No | Nomenclature for intact lipid molecule | Yes |
| MS2 verified by standard | No | Nomenclature for fragment ions | No |
| Background check at MS1 | Yes | Further identification remarks | - |

#### 58) PE O[M-H]- / Lipid quantification

| | | | |
|---|---|---|---|
| Quantitative | Yes | Limit of quantification | No |
| MS Level for quantification | MS1 | Normalization to reference | No |
| Internal lipid standard(s) MS1 | | Lipid Quantification Software | MS-DIAL |

| Internal standard | Endogenous subclass |
|---|---|
| PE 15:0_18:1(d7) | EtherPE subclass |

| | | | |
|---|---|---|---|
| Type of quantification | Internal standard amount | Batch correction | No |
| Response correction | No | Further quantification remarks | - |
| Type I isotope correction | No | | |

#### 59) Ether-linked phosphatidylglycerol (EtherPG)[M-H]- / Lipid identification

| | | | |
|---|---|---|---|
| Lipid class | Ether-linked phosphatidylglycerol (EtherPG) | Background check at MS2 | No |
| Derivatization | - | Did you presume assumptions for identification? | No |
| MS Level for identification | MS1, MS2 | Check isomer overlap | No |
| Identification level | Molecular species level | RT verified by standard | Yes |
| Polarity mode | Negative | Separation of isobaric/isomeric interferece confirmed | Yes |
| Type of negative (precursor)ion | [M-H]- | Model for separation prediction | Yes |
| Fragments for identification | | Additional dimension/techniques | - |

| Fragment name |
|---|
| Phosphoglycerol -H2O |
| Fatty acid fragment |
| Alkyl ether +O fragment |

| | | | |
|---|---|---|---|
| Isotope correction at MS1 | No | Lipid Identification Software | MS-DIAL |
| Isotope correction at MS2 | No | Data manipulation | Smoothing, Centroiding |
| MS1 verified by standard | No | Nomenclature for intact lipid molecule | Yes |
| MS2 verified by standard | No | Nomenclature for fragment ions | No |
| Background check at MS1 | Yes | Further identification remarks | - |

#### 59) Ether-linked phosphatidylglycerol (EtherPG)[M-H]- / Lipid quantification

| | | | |
|---|---|---|---|
| Quantitative | Yes | Limit of quantification | No |
| MS Level for quantification | MS1 | Normalization to reference | No |
| Internal lipid standard(s) MS1 | | Lipid Quantification Software | MS-DIAL |

| Internal standard | Endogenous subclass |
|---|---|
| PG 15:0_18:1(d7) | EtherPG subclass |

| | | | |
|---|---|---|---|
| Type of quantification | Internal standard amount | Batch correction | No |
| Response correction | No | Further quantification remarks | - |
| Type I isotope correction | No | | |

#### 60) Ether-linked phosphatidylinositol (EtherPI)[M-H]- / Lipid identification

| | | | |
|---|---|---|---|
| Lipid class | Ether-linked phosphatidylinositol (EtherPI) | Background check at MS2 | No |
| Derivatization | - | Did you presume assumptions for identification? | No |
| MS Level for identification | MS1, MS2 | Check isomer overlap | No |
| Identification level | Molecular species level | RT verified by standard | Yes |
| Polarity mode | Negative | Separation of isobaric/isomeric interferece confirmed | Yes |
| Type of negative (precursor)ion | [M-H]- | Model for separation prediction | Yes |
| Fragments for identification | | Additional dimension/techniques | - |

| Fragment name |
|---|
| Phosphoinositol -H2O |
| Fatty acid fragment |
| Alkyl ether + C3H5O4P fragment |

| | | | |
|---|---|---|---|
| Isotope correction at MS1 | No | Lipid Identification Software | MS-DIAL |
| Isotope correction at MS2 | No | Data manipulation | Smoothing, Centroiding |
| MS1 verified by standard | No | Nomenclature for intact lipid molecule | Yes |
| MS2 verified by standard | No | Nomenclature for fragment ions | No |
| Background check at MS1 | Yes | Further identification remarks | - |

#### 60) Ether-linked phosphatidylinositol (EtherPI)[M-H]- / Lipid quantification

| | | | |
|---|---|---|---|
| Quantitative | Yes | Limit of quantification | No |
| MS Level for quantification | MS1 | Normalization to reference | No |
| Internal lipid standard(s) MS1 | | Lipid Quantification Software | MS-DIAL |

| Internal standard | Endogenous subclass |
|---|---|
| PI 15:0_18:1(d7) | EtherPI subclass |

| | | | |
|---|---|---|---|
| Type of quantification | Internal standard amount | Batch correction | No |
| Response correction | No | Further quantification remarks | - |
| Type I isotope correction | No | | |

#### 61) Ether-linked phosphatidylserine (EtherPS)[M-H]- / Lipid identification

| | | | |
|---|---|---|---|
| Lipid class | Ether-linked phosphatidylserine (EtherPS) | Background check at MS2 | No |
| Derivatization | - | Did you presume assumptions for identification? | No |
| MS Level for identification | MS1, MS2 | Check isomer overlap | No |
| Identification level | Molecular species level | RT verified by standard | Yes |
| Polarity mode | Negative | Separation of isobaric/isomeric interferece confirmed | Yes |
| Type of negative (precursor)ion | [M-H]- | Model for separation prediction | Yes |
| Fragments for identification | | Additional dimension/techniques | - |

| Fragment name |
|---|
| Neutral loss of C3H5NO2 |
| Neutral loss of 22:6 Acyl and H2O and C3H5NO2 |
| Fatty acid fragment |

| | | | |
|---|---|---|---|
| Isotope correction at MS1 | No | Lipid Identification Software | MS-DIAL |
| Isotope correction at MS2 | No | Data manipulation | Smoothing, Centroiding |
| MS1 verified by standard | No | Nomenclature for intact lipid molecule | Yes |
| MS2 verified by standard | No | Nomenclature for fragment ions | No |
| Background check at MS1 | Yes | Further identification remarks | - |

#### 61) Ether-linked phosphatidylserine (EtherPS)[M-H]- / Lipid quantification

| | | | |
|---|---|---|---|
| Quantitative | Yes | Limit of quantification | No |
| MS Level for quantification | MS1 | Normalization to reference | No |
| Internal lipid standard(s) MS1 | | Lipid Quantification Software | MS-DIAL |

| Internal standard | Endogenous subclass |
|---|---|
| PS 15:0_18:1(d7) | EtherPS subclass |

| | | | |
|---|---|---|---|
| Type of quantification | Internal standard amount | Batch correction | No |
| Response correction | No | Further quantification remarks | - |
| Type I isotope correction | No | | |

#### 62) Ether-linked triacylglycerol (EtherTG)[M+NH4]+ / Lipid identification

| | | | |
|---|---|---|---|
| Lipid class | Ether-linked triacylglycerol (EtherTG) | Background check at MS2 | No |
| Derivatization | - | Did you presume assumptions for identification? | No |
| MS Level for identification | MS1, MS2 | Check isomer overlap | No |
| Identification level | Molecular species level | RT verified by standard | Yes |
| Polarity mode | Positive | Separation of isobaric/isomeric interferece confirmed | Yes |
| Type of positive (precursor)ion | [M+NH4]+ | Model for separation prediction | Yes |
| Fragments for identification | | Additional dimension/techniques | - |

| Fragment name |
|---|
| Neutral loss of fatty acyl and H2O |
| Neutral loss of alkyl ether |

| | | | |
|---|---|---|---|
| Isotope correction at MS1 | No | Lipid Identification Software | MS-DIAL |
| Isotope correction at MS2 | No | Data manipulation | Smoothing, Centroiding |
| MS1 verified by standard | No | Nomenclature for intact lipid molecule | Yes |
| MS2 verified by standard | No | Nomenclature for fragment ions | No |
| Background check at MS1 | Yes | Further identification remarks | - |

#### 62) Ether-linked triacylglycerol (EtherTG)[M+NH4]+ / Lipid quantification

| | | | |
|---|---|---|---|
| Quantitative | Yes | Limit of quantification | No |
| MS Level for quantification | MS1 | Normalization to reference | No |
| Internal lipid standard(s) MS1 | | Lipid Quantification Software | MS-DIAL |

| Internal standard | Endogenous subclass |
|---|---|
| TG 15:0_18:1(d7)_15:0 | EtherTG subclass |

| | | | |
|---|---|---|---|
| Type of quantification | Internal standard amount | Batch correction | No |
| Response correction | No | Further quantification remarks | - |
| Type I isotope correction | No | | |

#### 63) Fatty acid ester of hydroxyl fatty acid (FAHFA)[M-H]- / Lipid identification

| | | | |
|---|---|---|---|
| Lipid class | Fatty acid ester of hydroxyl fatty acid (FAHFA) | Background check at MS2 | No |
| Derivatization | - | Did you presume assumptions for identification? | No |
| MS Level for identification | MS1, MS2 | Check isomer overlap | No |
| Identification level | Molecular species level | RT verified by standard | Yes |
| Polarity mode | Negative | Separation of isobaric/isomeric interferece confirmed | Yes |
| Type of negative (precursor)ion | [M-H]- | Model for separation prediction | Yes |
| Fragments for identification | | Additional dimension/techniques | - |

| Fragment name |
|---|
| Neutral loss of fatty acyl and H2O |
| Fatty acid fragment |

| | | | |
|---|---|---|---|
| Isotope correction at MS1 | No | Lipid Identification Software | MS-DIAL |
| Isotope correction at MS2 | No | Data manipulation | Smoothing, Centroiding |
| MS1 verified by standard | No | Nomenclature for intact lipid molecule | Yes |
| MS2 verified by standard | No | Nomenclature for fragment ions | No |
| Background check at MS1 | Yes | Further identification remarks | - |

#### 63) Fatty acid ester of hydroxyl fatty acid (FAHFA)[M-H]- / Lipid quantification

| | | | |
|---|---|---|---|
| Quantitative | Yes | Limit of quantification | No |
| MS Level for quantification | MS1 | Normalization to reference | No |
| Internal lipid standard(s) MS1 | | Lipid Quantification Software | MS-DIAL |

| Internal standard | Endogenous subclass |
|---|---|
| FA 18:0(d3) | FAHFA subclass |

| | | | |
|---|---|---|---|
| Type of quantification | Internal standard amount | Batch correction | No |
| Response correction | No | Further quantification remarks | - |
| Type I isotope correction | No | | |

#### 64) Ganglioside GD1a (GD1a)[M-H]- / Lipid identification

| | | | |
|---|---|---|---|
| Lipid class | Ganglioside GD1a (GD1a) | Background check at MS2 | No |
| Derivatization | - | Did you presume assumptions for identification? | No |
| MS Level for identification | MS1, MS2 | Check isomer overlap | No |
| Identification level | Species level | RT verified by standard | Yes |
| Polarity mode | Negative | Separation of isobaric/isomeric interferece confirmed | Yes |
| Type of negative (precursor)ion | [M-H]- | Model for separation prediction | Yes |
| Fragments for identification | | Additional dimension/techniques | - |

| Fragment name |
|---|
| Characteristic fragment (C11H16NO8-) |

| | | | |
|---|---|---|---|
| Isotope correction at MS1 | No | Lipid Identification Software | MS-DIAL |
| Isotope correction at MS2 | No | Data manipulation | Smoothing, Centroiding |
| MS1 verified by standard | No | Nomenclature for intact lipid molecule | Yes |
| MS2 verified by standard | No | Nomenclature for fragment ions | No |
| Background check at MS1 | Yes | Further identification remarks | - |

#### 64) Ganglioside GD1a (GD1a)[M-H]- / Lipid quantification

| | | | |
|---|---|---|---|
| Quantitative | Yes | Limit of quantification | No |
| MS Level for quantification | MS1 | Normalization to reference | No |
| Internal lipid standard(s) MS1 | | Lipid Quantification Software | MS-DIAL |

| Internal standard | Endogenous subclass |
|---|---|
| LPC 18:1(d7) | GD1a subclass |

| | | | |
|---|---|---|---|
| Type of quantification | Internal standard amount | Batch correction | No |
| Response correction | No | Further quantification remarks | - |
| Type I isotope correction | No | | |

#### 65) Ganglioside GD1b (GD1b)[M-H]- / Lipid identification

| | | | |
|---|---|---|---|
| Lipid class | Ganglioside GD1b (GD1b) | Background check at MS2 | No |
| Derivatization | - | Did you presume assumptions for identification? | No |
| MS Level for identification | MS1, MS2 | Check isomer overlap | No |
| Identification level | Species level | RT verified by standard | Yes |
| Polarity mode | Negative | Separation of isobaric/isomeric interferece confirmed | Yes |
| Type of negative (precursor)ion | [M-H]- | Model for separation prediction | Yes |
| Fragments for identification | | Additional dimension/techniques | - |

| Fragment name |
|---|
| Characteristic fragment (C11H16NO8-) |

| | | | |
|---|---|---|---|
| Isotope correction at MS1 | No | Lipid Identification Software | MS-DIAL |
| Isotope correction at MS2 | No | Data manipulation | Smoothing, Centroiding |
| MS1 verified by standard | No | Nomenclature for intact lipid molecule | Yes |
| MS2 verified by standard | No | Nomenclature for fragment ions | No |
| Background check at MS1 | Yes | Further identification remarks | - |

#### 65) Ganglioside GD1b (GD1b)[M-H]- / Lipid quantification

| | | | |
|---|---|---|---|
| Quantitative | Yes | Limit of quantification | No |
| MS Level for quantification | MS1 | Normalization to reference | No |
| Internal lipid standard(s) MS1 | | Lipid Quantification Software | MS-DIAL |

| Internal standard | Endogenous subclass |
|---|---|
| LPC 18:1(d7) | GD1b subclass |

| | | | |
|---|---|---|---|
| Type of quantification | Internal standard amount | Batch correction | No |
| Response correction | No | Further quantification remarks | - |
| Type I isotope correction | No | | |

#### 66) Ganglioside GD2 (GD2)[M-H]- / Lipid identification

| | | | |
|---|---|---|---|
| Lipid class | Ganglioside GD2 (GD2) | Background check at MS2 | No |
| Derivatization | - | Did you presume assumptions for identification? | No |
| MS Level for identification | MS1, MS2 | Check isomer overlap | No |
| Identification level | Species level | RT verified by standard | Yes |
| Polarity mode | Negative | Separation of isobaric/isomeric interferece confirmed | Yes |
| Type of negative (precursor)ion | [M-H]- | Model for separation prediction | Yes |
| Fragments for identification | | Additional dimension/techniques | - |

| Fragment name |
|---|
| Characteristic fragment (C11H16NO8-) |

| | | | |
|---|---|---|---|
| Isotope correction at MS1 | No | Lipid Identification Software | MS-DIAL |
| Isotope correction at MS2 | No | Data manipulation | Smoothing, Centroiding |
| MS1 verified by standard | No | Nomenclature for intact lipid molecule | Yes |
| MS2 verified by standard | No | Nomenclature for fragment ions | No |
| Background check at MS1 | Yes | Further identification remarks | - |

#### 66) Ganglioside GD2 (GD2)[M-H]- / Lipid quantification

| | | | |
|---|---|---|---|
| Quantitative | Yes | Limit of quantification | No |
| MS Level for quantification | MS1 | Normalization to reference | No |
| Internal lipid standard(s) MS1 | | Lipid Quantification Software | MS-DIAL |

| Internal standard | Endogenous subclass |
|---|---|
| LPC 18:1(d7) | GD2 subclass |

| | | | |
|---|---|---|---|
| Type of quantification | Internal standard amount | Batch correction | No |
| Response correction | No | Further quantification remarks | - |
| Type I isotope correction | No | | |

#### 67) GD3[M-H]- / Lipid identification

| | | | |
|---|---|---|---|
| Lipid class | GD3 | Background check at MS2 | No |
| Derivatization | - | Did you presume assumptions for identification? | No |
| MS Level for identification | MS1, MS2 | Check isomer overlap | No |
| Identification level | Species level | RT verified by standard | Yes |
| Polarity mode | Negative | Separation of isobaric/isomeric interferece confirmed | Yes |
| Type of negative (precursor)ion | [M-H]- | Model for separation prediction | Yes |
| Fragments for identification | | Additional dimension/techniques | - |

| Fragment name |
|---|
| Characteristic fragment (C11H16NO8-) |

| | | | |
|---|---|---|---|
| Isotope correction at MS1 | No | Lipid Identification Software | MS-DIAL |
| Isotope correction at MS2 | No | Data manipulation | Smoothing, Centroiding |
| MS1 verified by standard | No | Nomenclature for intact lipid molecule | Yes |
| MS2 verified by standard | No | Nomenclature for fragment ions | No |
| Background check at MS1 | Yes | Further identification remarks | - |

#### 67) GD3[M-H]- / Lipid quantification

| | | | |
|---|---|---|---|
| Quantitative | Yes | Limit of quantification | No |
| MS Level for quantification | MS1 | Normalization to reference | No |
| Internal lipid standard(s) MS1 | | Lipid Quantification Software | MS-DIAL |

| Internal standard | Endogenous subclass |
|---|---|
| LPC 18:1(d7) | GD3 subclass |

| | | | |
|---|---|---|---|
| Type of quantification | Internal standard amount | Batch correction | No |
| Response correction | No | Further quantification remarks | - |
| Type I isotope correction | No | | |

#### 68) GM1[M-H]- / Lipid identification

| | | | |
|---|---|---|---|
| Lipid class | GM1 | Background check at MS2 | No |
| Derivatization | - | Did you presume assumptions for identification? | No |
| MS Level for identification | MS1, MS2 | Check isomer overlap | No |
| Identification level | Species level | RT verified by standard | Yes |
| Polarity mode | Negative | Separation of isobaric/isomeric interferece confirmed | Yes |
| Type of negative (precursor)ion | [M-H]- | Model for separation prediction | Yes |
| Fragments for identification | | Additional dimension/techniques | - |

| Fragment name |
|---|
| Characteristic fragment (C11H16NO8-) |

| | | | |
|---|---|---|---|
| Isotope correction at MS1 | No | Lipid Identification Software | MS-DIAL |
| Isotope correction at MS2 | No | Data manipulation | Smoothing, Centroiding |
| MS1 verified by standard | No | Nomenclature for intact lipid molecule | Yes |
| MS2 verified by standard | No | Nomenclature for fragment ions | No |
| Background check at MS1 | Yes | Further identification remarks | - |

#### 68) GM1[M-H]- / Lipid quantification

| | | | |
|---|---|---|---|
| Quantitative | Yes | Limit of quantification | No |
| MS Level for quantification | MS1 | Normalization to reference | No |
| Internal lipid standard(s) MS1 | | Lipid Quantification Software | MS-DIAL |

| Internal standard | Endogenous subclass |
|---|---|
| LPC 18:1(d7) | GM1 subclass |

| | | | |
|---|---|---|---|
| Type of quantification | Internal standard amount | Batch correction | No |
| Response correction | No | Further quantification remarks | - |
| Type I isotope correction | No | | |

#### 69) GM3[M+NH4]+ / Lipid identification

| | | | |
|---|---|---|---|
| Lipid class | GM3 | Background check at MS2 | No |
| Derivatization | - | Did you presume assumptions for identification? | No |
| MS Level for identification | MS1, MS2 | Check isomer overlap | No |
| Identification level | Species level | RT verified by standard | Yes |
| Polarity mode | Positive | Separation of isobaric/isomeric interferece confirmed | Yes |
| Type of positive (precursor)ion | [M+NH4]+ | Model for separation prediction | Yes |
| Fragments for identification | | Additional dimension/techniques | - |

| Fragment name |
|---|
| Neutral loss of H2O |
| Neutral loss of H2O and C23H37NO18 |
| Sphingosine -H2O fragment |
| Sphingosine -2H2O fragment |

| | | | |
|---|---|---|---|
| Isotope correction at MS1 | No | Lipid Identification Software | MS-DIAL |
| Isotope correction at MS2 | No | Data manipulation | Smoothing, Centroiding |
| MS1 verified by standard | No | Nomenclature for intact lipid molecule | Yes |
| MS2 verified by standard | No | Nomenclature for fragment ions | No |
| Background check at MS1 | Yes | Further identification remarks | - |

#### 69) GM3[M+NH4]+ / Lipid quantification

| | | | |
|---|---|---|---|
| Quantitative | Yes | Limit of quantification | No |
| MS Level for quantification | MS1 | Normalization to reference | No |
| Internal lipid standard(s) MS1 | | Lipid Quantification Software | MS-DIAL |

| Internal standard | Endogenous subclass |
|---|---|
| LPC 18:1(d7) | GM3 subclass |

| | | | |
|---|---|---|---|
| Type of quantification | Internal standard amount | Batch correction | No |
| Response correction | No | Further quantification remarks | - |
| Type I isotope correction | No | | |

#### 70) Ganglioside GQ1b (GQ1b)[M-2H]2- / Lipid identification

| | | | |
|---|---|---|---|
| Lipid class | Ganglioside GQ1b (GQ1b) | Background check at MS2 | No |
| Derivatization | - | Did you presume assumptions for identification? | No |
| MS Level for identification | MS1, MS2 | Check isomer overlap | No |
| Identification level | Species level | RT verified by standard | Yes |
| Polarity mode | Negative | Separation of isobaric/isomeric interferece confirmed | Yes |
| Type of negative (precursor)ion | [M-2H]2- | Model for separation prediction | Yes |
| Fragments for identification | | Additional dimension/techniques | - |

| Fragment name |
|---|
| Characteristic fragment (C11H16NO8-) |
| Characteristic fragment (C22H33N2O16-) |

| | | | |
|---|---|---|---|
| Isotope correction at MS1 | No | Lipid Identification Software | MS-DIAL |
| Isotope correction at MS2 | No | Data manipulation | Smoothing, Centroiding |
| MS1 verified by standard | No | Nomenclature for intact lipid molecule | Yes |
| MS2 verified by standard | No | Nomenclature for fragment ions | No |
| Background check at MS1 | Yes | Further identification remarks | - |

#### 70) Ganglioside GQ1b (GQ1b)[M-2H]2- / Lipid quantification

| | | | |
|---|---|---|---|
| Quantitative | Yes | Limit of quantification | No |
| MS Level for quantification | MS1 | Normalization to reference | No |
| Internal lipid standard(s) MS1 | | Lipid Quantification Software | MS-DIAL |

| Internal standard | Endogenous subclass |
|---|---|
| LPC 18:1(d7) | GQ1b subclass |

| | | | |
|---|---|---|---|
| Type of quantification | Internal standard amount | Batch correction | No |
| Response correction | No | Further quantification remarks | - |
| Type I isotope correction | No | | |

#### 71) Ganglioside GT1b (GT1b)[M-2H]2- / Lipid identification

| | | | |
|---|---|---|---|
| Lipid class | Ganglioside GT1b (GT1b) | Background check at MS2 | No |
| Derivatization | - | Did you presume assumptions for identification? | No |
| MS Level for identification | MS1, MS2 | Check isomer overlap | No |
| Identification level | Species level | RT verified by standard | Yes |
| Polarity mode | Negative | Separation of isobaric/isomeric interferece confirmed | Yes |
| Type of negative (precursor)ion | [M-2H]2- | Model for separation prediction | Yes |
| Fragments for identification | | Additional dimension/techniques | - |

| Fragment name |
|---|
| Characteristic fragment (C11H16NO8-) |
| Characteristic fragment (C22H33N2O16-) |

| | | | |
|---|---|---|---|
| Isotope correction at MS1 | No | Lipid Identification Software | MS-DIAL |
| Isotope correction at MS2 | No | Data manipulation | Smoothing, Centroiding |
| MS1 verified by standard | No | Nomenclature for intact lipid molecule | Yes |
| MS2 verified by standard | No | Nomenclature for fragment ions | No |
| Background check at MS1 | Yes | Further identification remarks | - |

#### 71) Ganglioside GT1b (GT1b)[M-2H]2- / Lipid quantification

| | | | |
|---|---|---|---|
| Quantitative | Yes | Limit of quantification | No |
| MS Level for quantification | MS1 | Normalization to reference | No |
| Internal lipid standard(s) MS1 | | Lipid Quantification Software | MS-DIAL |

| Internal standard | Endogenous subclass |
|---|---|
| LPC 18:1(d7) | GT1b subclass |

| | | | |
|---|---|---|---|
| Type of quantification | Internal standard amount | Batch correction | No |
| Response correction | No | Further quantification remarks | - |
| Type I isotope correction | No | | |

#### 72) Glycerophospho N-acyl ethanolamine (GPNAE)[M-H]- / Lipid identification

| | | | |
|---|---|---|---|
| Lipid class | Glycerophospho N-acyl ethanolamine (GPNAE) | Background check at MS2 | No |
| Derivatization | - | Did you presume assumptions for identification? | No |
| MS Level for identification | MS1, MS2 | Check isomer overlap | No |
| Identification level | Molecular species level | RT verified by standard | Yes |
| Polarity mode | Negative | Separation of isobaric/isomeric interferece confirmed | Yes |
| Type of negative (precursor)ion | [M-H]- | Model for separation prediction | Yes |
| Fragments for identification | | Additional dimension/techniques | - |

| Fragment name |
|---|
| Characteristic fragment (C3H8PO6-) |
| Phosphite |

| | | | |
|---|---|---|---|
| Isotope correction at MS1 | No | Lipid Identification Software | MS-DIAL |
| Isotope correction at MS2 | No | Data manipulation | Smoothing, Centroiding |
| MS1 verified by standard | No | Nomenclature for intact lipid molecule | Yes |
| MS2 verified by standard | No | Nomenclature for fragment ions | No |
| Background check at MS1 | Yes | Further identification remarks | - |

#### 72) Glycerophospho N-acyl ethanolamine (GPNAE)[M-H]- / Lipid quantification

| | | | |
|---|---|---|---|
| Quantitative | Yes | Limit of quantification | No |
| MS Level for quantification | MS1 | Normalization to reference | No |
| Internal lipid standard(s) MS1 | | Lipid Quantification Software | MS-DIAL |

| Internal standard | Endogenous subclass |
|---|---|
| LPC 18:1(d7) | GPNAE subclass |

| | | | |
|---|---|---|---|
| Type of quantification | Internal standard amount | Batch correction | No |
| Response correction | No | Further quantification remarks | - |
| Type I isotope correction | No | | |

#### 73) Hemibismonoacylglycerophosphate (HBMP)[M+NH4]+ / Lipid identification

| | | | |
|---|---|---|---|
| Lipid class | Hemibismonoacylglycerophosphate (HBMP) | Background check at MS2 | No |
| Derivatization | - | Did you presume assumptions for identification? | No |
| MS Level for identification | MS1, MS2 | Check isomer overlap | No |
| Identification level | Molecular species level | RT verified by standard | Yes |
| Polarity mode | Positive | Separation of isobaric/isomeric interferece confirmed | Yes |
| Type of positive (precursor)ion | [M+NH4]+ | Model for separation prediction | Yes |
| Fragments for identification | | Additional dimension/techniques | - |
| Isotope correction at MS1 | No | Lipid Identification Software | MS-DIAL |
| Isotope correction at MS2 | No | Data manipulation | Smoothing, Centroiding |
| MS1 verified by standard | No | Nomenclature for intact lipid molecule | Yes |
| MS2 verified by standard | No | Nomenclature for fragment ions | No |
| Background check at MS1 | Yes | Further identification remarks | - |

Fragments for identification:

| Fragment name |
|---|
| Dehydro-monoacyl glycerols |
| Dehydro-diacyl glycerol |

#### 73) Hemibismonoacylglycerophosphate (HBMP)[M+NH4]+ / Lipid quantification

| | | | |
|---|---|---|---|
| Quantitative | Yes | Limit of quantification | No |
| MS Level for quantification | MS1 | Normalization to reference | No |
| Internal lipid standard(s) MS1 | | Lipid Quantification Software | MS-DIAL |
| Type of quantification | Internal standard amount | Batch correction | No |
| Response correction | No | Further quantification remarks | - |
| Type I isotope correction | No | | |

Internal lipid standard(s) MS1:

| Internal standard | Endogenous subclass |
|---|---|
| PG 15:0_18:1(d7) | HBMP subclass |

#### 74) Hexosylceramide alpha-hydroxy fatty acid-phytospingosine (HexCer_AP)[M+H]+ / Lipid identification

| | | | |
|---|---|---|---|
| Lipid class | Hexosylceramide alpha-hydroxy fatty acid-phytospingosine (HexCer_AP) | Background check at MS2 | No |
| Derivatization | - | Did you presume assumptions for identification? | No |
| MS Level for identification | MS1, MS2 | Check isomer overlap | No |
| Identification level | Molecular species level | RT verified by standard | Yes |
| Polarity mode | Positive | Separation of isobaric/isomeric interferece confirmed | Yes |
| Type of positive (precursor)ion | [M+H]+ | Model for separation prediction | Yes |
| Fragments for identification | | Additional dimension/techniques | - |

| Fragment name |
|---|
| Neutral loss of hexose |
| Neutral loss of hexose and H2O |
| Phytosphingosine |
| Phytosphingosine -H2O fragment |
| Phytosphingosine -3H2O fragment |

| | | | |
|---|---|---|---|
| Isotope correction at MS1 | No | Lipid Identification Software | MS-DIAL |
| Isotope correction at MS2 | No | Data manipulation | Smoothing, Centroiding |
| MS1 verified by standard | No | Nomenclature for intact lipid molecule | Yes |
| MS2 verified by standard | No | Nomenclature for fragment ions | No |
| Background check at MS1 | Yes | Further identification remarks | - |

#### 74) Hexosylceramide alpha-hydroxy fatty acid-phytospingosine (HexCer_AP)[M+H]+ / Lipid quantification

| | | | |
|---|---|---|---|
| Quantitative | Yes | Limit of quantification | No |
| MS Level for quantification | MS1 | Normalization to reference | No |
| Internal lipid standard(s) MS1 | | Lipid Quantification Software | MS-DIAL |

| Internal standard | Endogenous subclass |
|---|---|
| Cer 18:1;2O/15:0(d7) | HexCer_AP subclass |

| | | | |
|---|---|---|---|
| Type of quantification | Internal standard amount | Batch correction | No |
| Response correction | No | Further quantification remarks | - |
| Type I isotope correction | No | | |

#### 75) Hexosylceramide Esterified omega-hydroxy fatty acid-sphingosine (HexCer_EOS)[M+H]+ / Lipid identification

| | | | |
|---|---|---|---|
| Lipid class | Hexosylceramide Esterified omega-hydroxy fatty acid-sphingosine (HexCer_EOS) | Background check at MS2 | No |
| Derivatization | - | Did you presume assumptions for identification? | No |
| MS Level for identification | MS1, MS2 | Check isomer overlap | No |
| Identification level | Molecular species level | RT verified by standard | Yes |
| Polarity mode | Positive | Separation of isobaric/isomeric interferece confirmed | Yes |
| Type of positive (precursor)ion | [M+H]+ | Model for separation prediction | Yes |
| Fragments for identification | | Additional dimension/techniques | - |

| Fragment name |
|---|
| Neutral loss of hexose |
| Neutral loss of hexose and H2O |
| Sphingosine -H2O fragment |
| Sphingosine -2H2O fragment |
| Sphingosine -CH4O2 fragment |

| | | | |
|---|---|---|---|
| Isotope correction at MS1 | No | Lipid Identification Software | MS-DIAL |
| Isotope correction at MS2 | No | Data manipulation | Smoothing, Centroiding |
| MS1 verified by standard | No | Nomenclature for intact lipid molecule | Yes |
| MS2 verified by standard | No | Nomenclature for fragment ions | No |
| Background check at MS1 | Yes | Further identification remarks | - |

#### 75) Hexosylceramide Esterified omega-hydroxy fatty acid-sphingosine (HexCer_EOS)[M+H]+ / Lipid quantification

| | | | |
|---|---|---|---|
| Quantitative | Yes | Limit of quantification | No |
| MS Level for quantification | MS1 | Normalization to reference | No |
| Internal lipid standard(s) MS1 | | Lipid Quantification Software | MS-DIAL |

| Internal standard | Endogenous subclass |
|---|---|
| Cer 18:1;2O/15:0(d7) | HexCer_EOS subclass |

| | | | |
|---|---|---|---|
| Type of quantification | Internal standard amount | Batch correction | No |
| Response correction | No | Further quantification remarks | - |
| Type I isotope correction | No | | |

#### 76) Hexosylceramide hydroxyfatty acid-dihydrosphingosine (HexCer_HDS)[M+H]+ / Lipid identification

| | | | |
|---|---|---|---|
| Lipid class | Hexosylceramide hydroxyfatty acid-dihydrosphingosine (HexCer_HDS) | Background check at MS2 | No |
| Derivatization | - | Did you presume assumptions for identification? | No |
| MS Level for identification | MS1, MS2 | Check isomer overlap | No |
| Identification level | Molecular species level | RT verified by standard | Yes |
| Polarity mode | Positive | Separation of isobaric/isomeric interferece confirmed | Yes |
| Type of positive (precursor)ion | [M+H]+ | Model for separation prediction | Yes |
| Fragments for identification | | Additional dimension/techniques | - |

| Fragment name |
|---|
| Neutral loss of hexose |
| Neutral loss of hexose and H2O |
| Sphinganine -H2O fragment |
| Sphinganine -2H2O fragment |
| Sphinganine -CH4O2 fragment |

| | | | |
|---|---|---|---|
| Isotope correction at MS1 | No | Lipid Identification Software | MS-DIAL |
| Isotope correction at MS2 | No | Data manipulation | Smoothing, Centroiding |
| MS1 verified by standard | No | Nomenclature for intact lipid molecule | Yes |
| MS2 verified by standard | No | Nomenclature for fragment ions | No |
| Background check at MS1 | Yes | Further identification remarks | - |

#### 76) Hexosylceramide hydroxyfatty acid-dihydrosphingosine (HexCer_HDS)[M+H]+ / Lipid quantification

| | | | |
|---|---|---|---|
| Quantitative | Yes | Limit of quantification | No |
| MS Level for quantification | MS1 | Normalization to reference | No |
| Internal lipid standard(s) MS1 | | Lipid Quantification Software | MS-DIAL |

| Internal standard | Endogenous subclass |
|---|---|
| Cer 18:1;2O/15:0(d7) | HexCer_HDS subclass |

| | | | |
|---|---|---|---|
| Type of quantification | Internal standard amount | Batch correction | No |
| Response correction | No | Further quantification remarks | - |
| Type I isotope correction | No | | |

#### 77) Hexosylceramide hydroxyfatty acid-sphingosine (HexCer_HS)[M+H]+ / Lipid identification

| | | | |
|---|---|---|---|
| Lipid class | Hexosylceramide hydroxyfatty acid-sphingosine (HexCer_HS) | Background check at MS2 | No |
| Derivatization | - | Did you presume assumptions for identification? | No |
| MS Level for identification | MS1, MS2 | Check isomer overlap | No |
| Identification level | Molecular species level | RT verified by standard | Yes |
| Polarity mode | Positive | Separation of isobaric/isomeric interferece confirmed | Yes |
| Type of positive (precursor)ion | [M+H]+ | Model for separation prediction | Yes |
| Fragments for identification | | Additional dimension/techniques | - |

| Fragment name |
|---|
| Neutral loss of hexose |
| Neutral loss of hexose and H2O |
| Sphingosine -H2O fragment |
| Sphingosine -2H2O fragment |
| Sphingosine -CH4O2 fragment |

| | | | |
|---|---|---|---|
| Isotope correction at MS1 | No | Lipid Identification Software | MS-DIAL |
| Isotope correction at MS2 | No | Data manipulation | Smoothing, Centroiding |
| MS1 verified by standard | No | Nomenclature for intact lipid molecule | Yes |
| MS2 verified by standard | No | Nomenclature for fragment ions | No |
| Background check at MS1 | Yes | Further identification remarks | - |

#### 77) Hexosylceramide hydroxyfatty acid-sphingosine (HexCer_HS)[M+H]+ / Lipid quantification

| | | | |
|---|---|---|---|
| Quantitative | Yes | Limit of quantification | No |
| MS Level for quantification | MS1 | Normalization to reference | No |
| Internal lipid standard(s) MS1 | | Lipid Quantification Software | MS-DIAL |

| Internal standard | Endogenous subclass |
|---|---|
| Cer 18:1;2O/15:0(d7) | HexCer_HS subclass |

| | | | |
|---|---|---|---|
| Type of quantification | Internal standard amount | Batch correction | No |
| Response correction | No | Further quantification remarks | - |
| Type I isotope correction | No | | |

#### 78) Hexosylceramide non-hydroxyfatty acid-dihydrosphingosine (HexCer_NDS)[M+H]+ / Lipid identification

| | | | |
|---|---|---|---|
| Lipid class | Hexosylceramide non-hydroxyfatty acid-dihydrosphingosine (HexCer_NDS) | Background check at MS2 | No |
| Derivatization | - | Did you presume assumptions for identification? | No |
| MS Level for identification | MS1, MS2 | Check isomer overlap | No |
| Identification level | Molecular species level | RT verified by standard | Yes |
| Polarity mode | Positive | Separation of isobaric/isomeric interferece confirmed | Yes |
| Type of positive (precursor)ion | [M+H]+ | Model for separation prediction | Yes |
| Fragments for identification | | Additional dimension/techniques | - |

| Fragment name |
|---|
| Neutral loss of hexose and H2O |
| Sphinganine -H2O fragment |
| Sphinganine -2H2O fragment |

| | | | |
|---|---|---|---|
| Isotope correction at MS1 | No | Lipid Identification Software | MS-DIAL |
| Isotope correction at MS2 | No | Data manipulation | Smoothing, Centroiding |
| MS1 verified by standard | No | Nomenclature for intact lipid molecule | Yes |
| MS2 verified by standard | No | Nomenclature for fragment ions | No |
| Background check at MS1 | Yes | Further identification remarks | - |

#### 78) Hexosylceramide non-hydroxyfatty acid-dihydrosphingosine (HexCer_NDS)[M+H]+ / Lipid quantification

| | | | |
|---|---|---|---|
| Quantitative | Yes | Limit of quantification | No |
| MS Level for quantification | MS1 | Normalization to reference | No |
| Internal lipid standard(s) MS1 | | Lipid Quantification Software | MS-DIAL |

| Internal standard | Endogenous subclass |
|---|---|
| Cer 18:1;2O/15:0(d7) | HexCer_NDS subclass |

| | | | |
|---|---|---|---|
| Type of quantification | Internal standard amount | Batch correction | No |
| Response correction | No | Further quantification remarks | - |
| Type I isotope correction | No | | |

#### 79) Hexosylceramide non-hydroxyfatty acid-sphingosine (HexCer_NS)[M+H]+ / Lipid identification

| | | | |
|---|---|---|---|
| Lipid class | Hexosylceramide non-hydroxyfatty acid-sphingosine (HexCer_NS) | Background check at MS2 | No |
| Derivatization | - | Did you presume assumptions for identification? | No |
| MS Level for identification | MS1, MS2 | Check isomer overlap | No |
| Identification level | Molecular species level | RT verified by standard | Yes |
| Polarity mode | Positive | Separation of isobaric/isomeric interferece confirmed | Yes |
| Type of positive (precursor)ion | [M+H]+ | Model for separation prediction | Yes |
| Fragments for identification | | Additional dimension/techniques | - |

| Fragment name |
|---|
| Neutral loss of H2O |
| Neutral loss of hexose and H2O |
| Sphingosine -H2O fragment |
| Sphingosine -2H2O fragment |
| Sphingosine -CH4O2 fragment |

| | | | |
|---|---|---|---|
| Isotope correction at MS1 | No | Lipid Identification Software | MS-DIAL |
| Isotope correction at MS2 | No | Data manipulation | Smoothing, Centroiding |
| MS1 verified by standard | No | Nomenclature for intact lipid molecule | Yes |
| MS2 verified by standard | No | Nomenclature for fragment ions | No |
| Background check at MS1 | Yes | Further identification remarks | - |

#### 79) Hexosylceramide non-hydroxyfatty acid-sphingosine (HexCer_NS)[M+H]+ / Lipid quantification

| | | | |
|---|---|---|---|
| Quantitative | Yes | Limit of quantification | No |
| MS Level for quantification | MS1 | Normalization to reference | No |
| Internal lipid standard(s) MS1 | | Lipid Quantification Software | MS-DIAL |

| Internal standard | Endogenous subclass |
|---|---|
| Cer 18:1;2O/15:0(d7) | HexCer_NS subclass |

| | | | |
|---|---|---|---|
| Type of quantification | Internal standard amount | Batch correction | No |
| Response correction | No | Further quantification remarks | - |
| Type I isotope correction | No | | |

#### 80) Lysocardiolipin (MLCL)[M-H]- / Lipid identification

| | | | |
|---|---|---|---|
| Lipid class | Lysocardiolipin (MLCL) | Background check at MS2 | No |
| Derivatization | - | Did you presume assumptions for identification? | No |
| MS Level for identification | MS1, MS2 | Check isomer overlap | No |
| Identification level | Molecular species level | RT verified by standard | Yes |
| Polarity mode | Negative | Separation of isobaric/isomeric interferece confirmed | Yes |
| Type of negative (precursor)ion | [M-H]- | Model for separation prediction | Yes |
| Fragments for identification | | Additional dimension/techniques | - |

| Fragment name |
|---|
| Phosphoglycerol -H2O fragment |
| Phosphatidic acid |
| Lysophosphatidic acid |
| Fatty acid fragment |

| | | | |
|---|---|---|---|
| Isotope correction at MS1 | No | Lipid Identification Software | MS-DIAL |
| Isotope correction at MS2 | No | Data manipulation | Smoothing, Centroiding |
| MS1 verified by standard | No | Nomenclature for intact lipid molecule | Yes |
| MS2 verified by standard | No | Nomenclature for fragment ions | No |
| Background check at MS1 | Yes | Further identification remarks | - |

#### 80) Lysocardiolipin (MLCL)[M-H]- / Lipid quantification

| | | | |
|---|---|---|---|
| Quantitative | Yes | Limit of quantification | No |
| MS Level for quantification | MS1 | Normalization to reference | No |
| Internal lipid standard(s) MS1 | | Lipid Quantification Software | MS-DIAL |

| Internal standard | Endogenous subclass |
|---|---|
| PG 15:0_18:1(d7) | MLCL subclass |

| | | | |
|---|---|---|---|
| Type of quantification | Internal standard amount | Batch correction | No |
| Response correction | No | Further quantification remarks | - |
| Type I isotope correction | No | | |

#### 81) Lysodiacylglyceryl-3-O-carboxyhydroxymethylcholine (LDGCC)[M+H]+ / Lipid identification

| | | | |
|---|---|---|---|
| Lipid class | Lysodiacylglyceryl-3-O-carboxyhydroxymethylcholine (LDGCC) | Background check at MS2 | No |
| Derivatization | - | Did you presume assumptions for identification? | No |
| MS Level for identification | MS1, MS2 | Check isomer overlap | No |
| Identification level | Molecular species level | RT verified by standard | Yes |
| Polarity mode | Positive | Separation of isobaric/isomeric interferece confirmed | Yes |
| Type of positive (precursor)ion | [M+H]+ | Model for separation prediction | Yes |
| Fragments for identification | | Additional dimension/techniques | - |
| Isotope correction at MS1 | No | Lipid Identification Software | MS-DIAL |
| Isotope correction at MS2 | No | Data manipulation | Smoothing, Centroiding |
| MS1 verified by standard | No | Nomenclature for intact lipid molecule | Yes |
| MS2 verified by standard | No | Nomenclature for fragment ions | No |
| Background check at MS1 | Yes | Further identification remarks | - |

Fragments for identification:

| Fragment name |
|---|
| Characteristic fragment (C6H14NO2+) |

#### 81) Lysodiacylglyceryl-3-O-carboxyhydroxymethylcholine (LDGCC)[M+H]+ / Lipid quantification

| | | | |
|---|---|---|---|
| Quantitative | Yes | Limit of quantification | No |
| MS Level for quantification | MS1 | Normalization to reference | No |
| Internal lipid standard(s) MS1 | | Lipid Quantification Software | MS-DIAL |
| Type of quantification | Internal standard amount | Batch correction | No |
| Response correction | No | Further quantification remarks | - |
| Type I isotope correction | No | | |

Internal lipid standard(s) MS1:

| Internal standard | Endogenous subclass |
|---|---|
| LPC 18:1(d7) | LDGCC subclass |

#### 82) Lysogiacylglyceryl trimethylhomoserine (LDGTS)[M+H]+ / Lipid identification

| | | | |
|---|---|---|---|
| Lipid class | Lysogiacylglyceryl trimethylhomoserine (LDGTS) | Background check at MS2 | No |
| Derivatization | - | Did you presume assumptions for identification? | No |
| MS Level for identification | MS1, MS2 | Check isomer overlap | No |
| Identification level | Molecular species level | RT verified by standard | Yes |
| Polarity mode | Positive | Separation of isobaric/isomeric interferece confirmed | Yes |
| Type of positive (precursor)ion | [M+H]+ | Model for separation prediction | Yes |
| Fragments for identification | | Additional dimension/techniques | - |

| Fragment name |
|---|
| Characteristic fragments (C7H14NO2+) |
| Characteristic fragments (C10H22NO5+) |

| | | | |
|---|---|---|---|
| Isotope correction at MS1 | No | Lipid Identification Software | MS-DIAL |
| Isotope correction at MS2 | No | Data manipulation | Smoothing, Centroiding |
| MS1 verified by standard | No | Nomenclature for intact lipid molecule | Yes |
| MS2 verified by standard | No | Nomenclature for fragment ions | No |
| Background check at MS1 | Yes | Further identification remarks | - |

#### 82) Lysogiacylglyceryl trimethylhomoserine (LDGTS)[M+H]+ / Lipid quantification

| | | | |
|---|---|---|---|
| Quantitative | Yes | Limit of quantification | No |
| MS Level for quantification | MS1 | Normalization to reference | No |
| Internal lipid standard(s) MS1 | | Lipid Quantification Software | MS-DIAL |

| Internal standard | Endogenous subclass |
|---|---|
| LPC 18:1(d7) | LDGTS subclass |

| | | | |
|---|---|---|---|
| Type of quantification | Internal standard amount | Batch correction | No |
| Response correction | No | Further quantification remarks | - |
| Type I isotope correction | No | | |

#### 83) LPC[M+H]+ / Lipid identification

| | | | |
|---|---|---|---|
| Lipid class | LPC | Background check at MS2 | No |
| Derivatization | - | Did you presume assumptions for identification? | No |
| MS Level for identification | MS1, MS2 | Check isomer overlap | No |
| Identification level | Molecular species level | RT verified by standard | Yes |
| Polarity mode | Positive | Separation of isobaric/isomeric interferece confirmed | Yes |
| Type of positive (precursor)ion | [M+H]+ | Model for separation prediction | Yes |
| Fragments for identification | | Additional dimension/techniques | - |

| Fragment name |
|---|
| Characteristic fragment (C5H15NO4P+) |

| | | | |
|---|---|---|---|
| Isotope correction at MS1 | No | Lipid Identification Software | MS-DIAL |
| Isotope correction at MS2 | No | Data manipulation | Smoothing, Centroiding |
| MS1 verified by standard | Yes | Nomenclature for intact lipid molecule | Yes |
| MS2 verified by standard | Yes | Nomenclature for fragment ions | No |
| Background check at MS1 | Yes | Further identification remarks | - |

#### 83) LPC[M+H]+ / Lipid quantification

| | | | |
|---|---|---|---|
| Quantitative | Yes | Limit of quantification | No |
| MS Level for quantification | MS1 | Normalization to reference | No |
| Internal lipid standard(s) MS1 | | Lipid Quantification Software | MS-DIAL |

| Internal standard | Endogenous subclass |
|---|---|
| LPC 18:1(d7) | LPC subclass |

| | | | |
|---|---|---|---|
| Type of quantification | Internal standard amount | Batch correction | No |
| Response correction | No | Further quantification remarks | - |
| Type I isotope correction | No | | |

#### 84) LPA[M-H]- / Lipid identification

| | | | |
|---|---|---|---|
| Lipid class | LPA | Background check at MS2 | No |
| Derivatization | - | Did you presume assumptions for identification? | No |
| MS Level for identification | MS1, MS2 | Check isomer overlap | No |
| Identification level | Molecular species level | RT verified by standard | Yes |
| Polarity mode | Negative | Separation of isobaric/isomeric interferece confirmed | Yes |
| Type of negative (precursor)ion | [M-H]- | Model for separation prediction | Yes |
| Fragments for identification | | Additional dimension/techniques | - |

| Fragment name |
|---|
| Phosphoglycerol -H2O fragment |

| | | | |
|---|---|---|---|
| Isotope correction at MS1 | No | Lipid Identification Software | MS-DIAL |
| Isotope correction at MS2 | No | Data manipulation | Smoothing, Centroiding |
| MS1 verified by standard | No | Nomenclature for intact lipid molecule | Yes |
| MS2 verified by standard | No | Nomenclature for fragment ions | No |
| Background check at MS1 | Yes | Further identification remarks | - |

#### 84) LPA[M-H]- / Lipid quantification

| | | | |
|---|---|---|---|
| Quantitative | Yes | Limit of quantification | No |
| MS Level for quantification | MS1 | Normalization to reference | No |
| Internal lipid standard(s) MS1 | | Lipid Quantification Software | MS-DIAL |

| Internal standard | Endogenous subclass |
|---|---|
| LPC 18:1(d7) | LPA subclass |

| | | | |
|---|---|---|---|
| Type of quantification | Internal standard amount | Batch correction | No |
| Response correction | No | Further quantification remarks | - |
| Type I isotope correction | No | | |

#### 85) LPE[M+H]+ / Lipid identification

| | | | |
|---|---|---|---|
| Lipid class | LPE | Background check at MS2 | No |
| Derivatization | - | Did you presume assumptions for identification? | No |
| MS Level for identification | MS1, MS2 | Check isomer overlap | No |
| Identification level | Molecular species level | RT verified by standard | Yes |
| Polarity mode | Positive | Separation of isobaric/isomeric interferece confirmed | Yes |
| Type of positive (precursor)ion | [M+H]+ | Model for separation prediction | Yes |
| Fragments for identification | | Additional dimension/techniques | - |

| Fragment name |
|---|
| Neutral loss of C2H8NO4P |

| | | | |
|---|---|---|---|
| Isotope correction at MS1 | No | Lipid Identification Software | MS-DIAL |
| Isotope correction at MS2 | No | Data manipulation | Smoothing, Centroiding |
| MS1 verified by standard | Yes | Nomenclature for intact lipid molecule | Yes |
| MS2 verified by standard | Yes | Nomenclature for fragment ions | No |
| Background check at MS1 | Yes | Further identification remarks | - |

#### 85) LPE[M+H]+ / Lipid quantification

| | | | |
|---|---|---|---|
| Quantitative | Yes | Limit of quantification | No |
| MS Level for quantification | MS1 | Normalization to reference | No |
| Internal lipid standard(s) MS1 | | Lipid Quantification Software | MS-DIAL |

| Internal standard | Endogenous subclass |
|---|---|
| LPE 18:1(d7) | LPE subclass |

| | | | |
|---|---|---|---|
| Type of quantification | Internal standard amount | Batch correction | No |
| Response correction | No | Further quantification remarks | - |
| Type I isotope correction | No | | |

#### 86) LPG[M-H]- / Lipid identification

| | | | |
|---|---|---|---|
| Lipid class | LPG | Background check at MS2 | No |
| Derivatization | - | Did you presume assumptions for identification? | No |
| MS Level for identification | MS1, MS2 | Check isomer overlap | No |
| Identification level | Molecular species level | RT verified by standard | Yes |
| Polarity mode | Negative | Separation of isobaric/isomeric interferece confirmed | Yes |
| Type of negative (precursor)ion | [M-H]- | Model for separation prediction | Yes |
| Fragments for identification | | Additional dimension/techniques | - |

| Fragment name |
|---|
| Phosphoglycerol -H2O fragment |

| | | | |
|---|---|---|---|
| Isotope correction at MS1 | No | Lipid Identification Software | MS-DIAL |
| Isotope correction at MS2 | No | Data manipulation | Smoothing, Centroiding |
| MS1 verified by standard | No | Nomenclature for intact lipid molecule | Yes |
| MS2 verified by standard | No | Nomenclature for fragment ions | No |
| Background check at MS1 | Yes | Further identification remarks | - |

#### 86) LPG[M-H]- / Lipid quantification

| | | | |
|---|---|---|---|
| Quantitative | Yes | Limit of quantification | No |
| MS Level for quantification | MS1 | Normalization to reference | No |
| Internal lipid standard(s) MS1 | | Lipid Quantification Software | MS-DIAL |

| Internal standard | Endogenous subclass |
|---|---|
| PG 15:0_18:1(d7) | LPG subclass |

| | | | |
|---|---|---|---|
| Type of quantification | Internal standard amount | Batch correction | No |
| Response correction | No | Further quantification remarks | - |
| Type I isotope correction | No | | |

#### 87) LPI[M-H]- / Lipid identification

| | | | |
|---|---|---|---|
| Lipid class | LPI | Background check at MS2 | No |
| Derivatization | - | Did you presume assumptions for identification? | No |
| MS Level for identification | MS1, MS2 | Check isomer overlap | No |
| Identification level | Molecular species level | RT verified by standard | Yes |
| Polarity mode | Negative | Separation of isobaric/isomeric interferece confirmed | Yes |
| Type of negative (precursor)ion | [M-H]- | Model for separation prediction | Yes |
| Fragments for identification | | Additional dimension/techniques | - |

| Fragment name |
|---|
| Phosphoinositol -H2O fragment |
| Characteristic fragment (C9H16O10P-) |
| Fatty acid fragment |

| | | | |
|---|---|---|---|
| Isotope correction at MS1 | No | Lipid Identification Software | MS-DIAL |
| Isotope correction at MS2 | No | Data manipulation | Smoothing, Centroiding |
| MS1 verified by standard | No | Nomenclature for intact lipid molecule | Yes |
| MS2 verified by standard | No | Nomenclature for fragment ions | No |
| Background check at MS1 | Yes | Further identification remarks | - |

#### 87) LPI[M-H]- / Lipid quantification

| | | | |
|---|---|---|---|
| Quantitative | Yes | Limit of quantification | No |
| MS Level for quantification | MS1 | Normalization to reference | No |
| Internal lipid standard(s) MS1 | | Lipid Quantification Software | MS-DIAL |

| Internal standard | Endogenous subclass |
|---|---|
| PI 15:0_18:1(d7) | LPI subclass |

| | | | |
|---|---|---|---|
| Type of quantification | Internal standard amount | Batch correction | No |
| Response correction | No | Further quantification remarks | - |
| Type I isotope correction | No | | |

#### 88) LPS[M-H]- / Lipid identification

| | | | |
|---|---|---|---|
| Lipid class | LPS | Background check at MS2 | No |
| Derivatization | - | Did you presume assumptions for identification? | No |
| MS Level for identification | MS1, MS2 | Check isomer overlap | No |
| Identification level | Molecular species level | RT verified by standard | Yes |
| Polarity mode | Negative | Separation of isobaric/isomeric interferece confirmed | Yes |
| Type of negative (precursor)ion | [M-H]- | Model for separation prediction | Yes |
| Fragments for identification | | Additional dimension/techniques | - |

| Fragment name |
|---|
| Neutral loss of C3H6NO2 |
| Phosphoglycerol -H2O fragment |

| | | | |
|---|---|---|---|
| Isotope correction at MS1 | No | Lipid Identification Software | MS-DIAL |
| Isotope correction at MS2 | No | Data manipulation | Smoothing, Centroiding |
| MS1 verified by standard | No | Nomenclature for intact lipid molecule | Yes |
| MS2 verified by standard | No | Nomenclature for fragment ions | No |
| Background check at MS1 | Yes | Further identification remarks | - |

#### 88) LPS[M-H]- / Lipid quantification

| | | | |
|---|---|---|---|
| Quantitative | Yes | Limit of quantification | No |
| MS Level for quantification | MS1 | Normalization to reference | No |
| Internal lipid standard(s) MS1 | | Lipid Quantification Software | MS-DIAL |

| Internal standard | Endogenous subclass |
|---|---|
| PS 15:0_18:1(d7) | LPS subclass |

| | | | |
|---|---|---|---|
| Type of quantification | Internal standard amount | Batch correction | No |
| Response correction | No | Further quantification remarks | - |
| Type I isotope correction | No | | |

#### 89) MIPC[M-H]- / Lipid identification

| | | | |
|---|---|---|---|
| Lipid class | MIPC | Background check at MS2 | No |
| Derivatization | - | Did you presume assumptions for identification? | No |
| MS Level for identification | MS1, MS2 | Check isomer overlap | No |
| Identification level | Molecular species level | RT verified by standard | Yes |
| Polarity mode | Negative | Separation of isobaric/isomeric interferece confirmed | Yes |
| Type of negative (precursor)ion | [M-H]- | Model for separation prediction | Yes |
| Fragments for identification | | Additional dimension/techniques | - |

| Fragment name |
|---|
| Characteristic fragment (C12H22O14P-) |
| Phytosphingosine -C2H7NO fragment |

| | | | |
|---|---|---|---|
| Isotope correction at MS1 | No | Lipid Identification Software | MS-DIAL |
| Isotope correction at MS2 | No | Data manipulation | Smoothing, Centroiding |
| MS1 verified by standard | No | Nomenclature for intact lipid molecule | Yes |
| MS2 verified by standard | No | Nomenclature for fragment ions | No |
| Background check at MS1 | Yes | Further identification remarks | - |

#### 89) MIPC[M-H]- / Lipid quantification

| | | | |
|---|---|---|---|
| Quantitative | Yes | Limit of quantification | No |
| MS Level for quantification | MS1 | Normalization to reference | No |
| Internal lipid standard(s) MS1 | | Lipid Quantification Software | MS-DIAL |

| Internal standard | Endogenous subclass |
|---|---|
| PI 15:0_18:1(d7) | MIPC subclass |

| | | | |
|---|---|---|---|
| Type of quantification | Internal standard amount | Batch correction | No |
| Response correction | No | Further quantification remarks | - |
| Type I isotope correction | No | | |

#### 90) MG[M+NH4]+ / Lipid identification

| | | | |
|---|---|---|---|
| Lipid class | MG | Background check at MS2 | No |
| Derivatization | - | Did you presume assumptions for identification? | No |
| MS Level for identification | MS1, MS2 | Check isomer overlap | No |
| Identification level | Molecular species level | RT verified by standard | Yes |
| Polarity mode | Positive | Separation of isobaric/isomeric interferece confirmed | Yes |
| Type of positive (precursor)ion | [M+NH4]+ | Model for separation prediction | Yes |
| Fragments for identification | | Additional dimension/techniques | - |

| Fragment name |
|---|
| Neutral loss of H2O |

| | | | |
|---|---|---|---|
| Isotope correction at MS1 | No | Lipid Identification Software | MS-DIAL |
| Isotope correction at MS2 | No | Data manipulation | Smoothing, Centroiding |
| MS1 verified by standard | Yes | Nomenclature for intact lipid molecule | Yes |
| MS2 verified by standard | Yes | Nomenclature for fragment ions | No |
| Background check at MS1 | Yes | Further identification remarks | - |

#### 90) MG[M+NH4]+ / Lipid quantification

| | | | |
|---|---|---|---|
| Quantitative | Yes | Limit of quantification | No |
| MS Level for quantification | MS1 | Normalization to reference | No |
| Internal lipid standard(s) MS1 | | Lipid Quantification Software | MS-DIAL |

| Internal standard | Endogenous subclass |
|---|---|
| MG 18:1(d7) | MG subclass |

| | | | |
|---|---|---|---|
| Type of quantification | Internal standard amount | Batch correction | No |
| Response correction | No | Further quantification remarks | - |
| Type I isotope correction | No | | |

#### 91) MGDG[M+CH3COO]- / Lipid identification

| | | | |
|---|---|---|---|
| Lipid class | MGDG | Background check at MS2 | No |
| Derivatization | - | Did you presume assumptions for identification? | No |
| MS Level for identification | MS1, MS2 | Check isomer overlap | No |
| Identification level | Molecular species level | RT verified by standard | Yes |
| Polarity mode | Negative | Separation of isobaric/isomeric interferece confirmed | Yes |
| Type of negative (precursor)ion | [M+CH3COO]- | Model for separation prediction | Yes |
| Fragments for identification | | Additional dimension/techniques | - |

| Fragment name |
|---|
| Fatty acid fragment |

| | | | |
|---|---|---|---|
| Isotope correction at MS1 | No | Lipid Identification Software | MS-DIAL |
| Isotope correction at MS2 | No | Data manipulation | Smoothing, Centroiding |
| MS1 verified by standard | No | Nomenclature for intact lipid molecule | Yes |
| MS2 verified by standard | No | Nomenclature for fragment ions | No |
| Background check at MS1 | Yes | Further identification remarks | - |

#### 91) MGDG[M+CH3COO]- / Lipid quantification

| | | | |
|---|---|---|---|
| Quantitative | Yes | Limit of quantification | No |
| MS Level for quantification | MS1 | Normalization to reference | No |
| Internal lipid standard(s) MS1 | | Lipid Quantification Software | MS-DIAL |

| Internal standard | Endogenous subclass |
|---|---|
| LPC 18:1(d7) | MGDG subclass |

| | | | |
|---|---|---|---|
| Type of quantification | Internal standard amount | Batch correction | No |
| Response correction | No | Further quantification remarks | - |
| Type I isotope correction | No | | |

#### 92) Monogalactosylmonoacylglycerol (MGMG)[M+CH3COO]- / Lipid identification

| | | | |
|---|---|---|---|
| Lipid class | Monogalactosylmonoacylglycerol (MGMG) | Background check at MS2 | No |
| Derivatization | - | Did you presume assumptions for identification? | No |
| MS Level for identification | MS1, MS2 | Check isomer overlap | No |
| Identification level | Molecular species level | RT verified by standard | Yes |
| Polarity mode | Negative | Separation of isobaric/isomeric interferece confirmed | Yes |
| Type of negative (precursor)ion | [M+CH3COO]- | Model for separation prediction | Yes |
| Fragments for identification | | Additional dimension/techniques | - |
| Isotope correction at MS1 | No | Lipid Identification Software | MS-DIAL |
| Isotope correction at MS2 | No | Data manipulation | Smoothing, Centroiding |
| MS1 verified by standard | No | Nomenclature for intact lipid molecule | Yes |
| MS2 verified by standard | No | Nomenclature for fragment ions | No |
| Background check at MS1 | Yes | Further identification remarks | - |

Fragments for identification:

| Fragment name |
|---|
| Fatty acid fragment |

#### 92) Monogalactosylmonoacylglycerol (MGMG)[M+CH3COO]- / Lipid quantification

| | | | |
|---|---|---|---|
| Quantitative | Yes | Limit of quantification | No |
| MS Level for quantification | MS1 | Normalization to reference | No |
| Internal lipid standard(s) MS1 | | Lipid Quantification Software | MS-DIAL |
| Type of quantification | Internal standard amount | Batch correction | No |
| Response correction | No | Further quantification remarks | - |
| Type I isotope correction | No | | |

Internal lipid standard(s) MS1:

| Internal standard | Endogenous subclass |
|---|---|
| LPC 18:1(d7) | MGMG subclass |

#### 93) N-acyl ethanolamines (NAE)[M+CH3COO]- / Lipid identification

| | | | |
|---|---|---|---|
| Lipid class | N-acyl ethanolamines (NAE) | Background check at MS2 | No |
| Derivatization | - | Did you presume assumptions for identification? | No |
| MS Level for identification | MS1, MS2 | Check isomer overlap | No |
| Identification level | Molecular species level | RT verified by standard | Yes |
| Polarity mode | Negative | Separation of isobaric/isomeric interferece confirmed | Yes |
| Type of negative (precursor)ion | [M+CH3COO]- | Model for separation prediction | Yes |
| Fragments for identification | | Additional dimension/techniques | - |

| Fragment name |
|---|
| Neutral loss of 2H |

| | | | |
|---|---|---|---|
| Isotope correction at MS1 | No | Lipid Identification Software | MS-DIAL |
| Isotope correction at MS2 | No | Data manipulation | Smoothing, Centroiding |
| MS1 verified by standard | No | Nomenclature for intact lipid molecule | Yes |
| MS2 verified by standard | No | Nomenclature for fragment ions | No |
| Background check at MS1 | Yes | Further identification remarks | - |

#### 93) N-acyl ethanolamines (NAE)[M+CH3COO]- / Lipid quantification

| | | | |
|---|---|---|---|
| Quantitative | Yes | Limit of quantification | No |
| MS Level for quantification | MS1 | Normalization to reference | No |
| Internal lipid standard(s) MS1 | | Lipid Quantification Software | MS-DIAL |

| Internal standard | Endogenous subclass |
|---|---|
| LPC 18:1(d7) | NAE subclass |

| | | | |
|---|---|---|---|
| Type of quantification | Internal standard amount | Batch correction | No |
| Response correction | No | Further quantification remarks | - |
| Type I isotope correction | No | | |

#### 94) N-acyl glycine (NAGly)[M+NH4]+ / Lipid identification

| | | | |
|---|---|---|---|
| Lipid class | N-acyl glycine (NAGly) | Background check at MS2 | No |
| Derivatization | - | Did you presume assumptions for identification? | No |
| MS Level for identification | MS1, MS2 | Check isomer overlap | No |
| Identification level | Molecular species level | RT verified by standard | Yes |
| Polarity mode | Positive | Separation of isobaric/isomeric interferece confirmed | Yes |
| Type of positive (precursor)ion | [M+NH4]+ | Model for separation prediction | Yes |
| Fragments for identification | | Additional dimension/techniques | - |

| Fragment name |
|---|
| Glycine |
| Fatty acyl fragment |
| Fatty acyl -H2O fragment |
| Neutral loss of Acyl and H2O |

| | | | |
|---|---|---|---|
| Isotope correction at MS1 | No | Lipid Identification Software | MS-DIAL |
| Isotope correction at MS2 | No | Data manipulation | Smoothing, Centroiding |
| MS1 verified by standard | No | Nomenclature for intact lipid molecule | Yes |
| MS2 verified by standard | No | Nomenclature for fragment ions | No |
| Background check at MS1 | Yes | Further identification remarks | - |

#### 94) N-acyl glycine (NAGly)[M+NH4]+ / Lipid quantification

| | | | |
|---|---|---|---|
| Quantitative | Yes | Limit of quantification | No |
| MS Level for quantification | MS1 | Normalization to reference | No |
| Internal lipid standard(s) MS1 | | Lipid Quantification Software | MS-DIAL |

| Internal standard | Endogenous subclass |
|---|---|
| LPC 18:1(d7) | NAGly subclass |

| | | | |
|---|---|---|---|
| Type of quantification | Internal standard amount | Batch correction | No |
| Response correction | No | Further quantification remarks | - |
| Type I isotope correction | No | | |

#### 95) N-acyl glycyl serine (NAGlySer)[M+NH4]+ / Lipid identification

| | | | |
|---|---|---|---|
| Lipid class | N-acyl glycyl serine (NAGlySer) | Background check at MS2 | No |
| Derivatization | - | Did you presume assumptions for identification? | No |
| MS Level for identification | MS1, MS2 | Check isomer overlap | No |
| Identification level | Molecular species level | RT verified by standard | Yes |
| Polarity mode | Positive | Separation of isobaric/isomeric interferece confirmed | Yes |
| Type of positive (precursor)ion | [M+NH4]+ | Model for separation prediction | Yes |
| Fragments for identification | | Additional dimension/techniques | - |

| Fragment name |
|---|
| Glycylserine |
| Fatty acyl fragment |
| Serine |
| Neutral loss of Acyl and H2O |
| Acyl Glycine fragment |

| | | | |
|---|---|---|---|
| Isotope correction at MS1 | No | Lipid Identification Software | MS-DIAL |
| Isotope correction at MS2 | No | Data manipulation | Smoothing, Centroiding |
| MS1 verified by standard | No | Nomenclature for intact lipid molecule | Yes |
| MS2 verified by standard | No | Nomenclature for fragment ions | No |
| Background check at MS1 | Yes | Further identification remarks | - |

#### 95) N-acyl glycyl serine (NAGlySer)[M+NH4]+ / Lipid quantification

| | | | |
|---|---|---|---|
| Quantitative | Yes | Limit of quantification | No |
| MS Level for quantification | MS1 | Normalization to reference | No |
| Internal lipid standard(s) MS1 | | Lipid Quantification Software | MS-DIAL |

| Internal standard | Endogenous subclass |
|---|---|
| LPC 18:1(d7) | NAGlySer subclass |

| | | | |
|---|---|---|---|
| Type of quantification | Internal standard amount | Batch correction | No |
| Response correction | No | Further quantification remarks | - |
| Type I isotope correction | No | | |

#### 96) N-acyl ornithine (NAOrn)[M+H]+ / Lipid identification

| | | | |
|---|---|---|---|
| Lipid class | N-acyl ornithine (NAOrn) | Background check at MS2 | No |
| Derivatization | - | Did you presume assumptions for identification? | No |
| MS Level for identification | MS1, MS2 | Check isomer overlap | No |
| Identification level | Molecular species level | RT verified by standard | Yes |
| Polarity mode | Positive | Separation of isobaric/isomeric interferece confirmed | Yes |
| Type of positive (precursor)ion | [M+H]+ | Model for separation prediction | Yes |
| Fragments for identification | | Additional dimension/techniques | - |

| Fragment name |
|---|
| Ornithine -H2O |
| Characteristic fragment (C4H8N+) |
| Neutral loss of Acyl and H2O |
| Neutral loss of Acyl and 2H2O |
| Fatty acyl -H2O fragment |

| | | | |
|---|---|---|---|
| Isotope correction at MS1 | No | Lipid Identification Software | MS-DIAL |
| Isotope correction at MS2 | No | Data manipulation | Smoothing, Centroiding |
| MS1 verified by standard | No | Nomenclature for intact lipid molecule | Yes |
| MS2 verified by standard | No | Nomenclature for fragment ions | No |
| Background check at MS1 | Yes | Further identification remarks | - |

#### 96) N-acyl ornithine (NAOrn)[M+H]+ / Lipid quantification

| | | | |
|---|---|---|---|
| Quantitative | Yes | Limit of quantification | No |
| MS Level for quantification | MS1 | Normalization to reference | No |
| Internal lipid standard(s) MS1 | | Lipid Quantification Software | MS-DIAL |

| Internal standard | Endogenous subclass |
|---|---|
| LPC 18:1(d7) | NAOrn subclass |

| | | | |
|---|---|---|---|
| Type of quantification | Internal standard amount | Batch correction | No |
| Response correction | No | Further quantification remarks | - |
| Type I isotope correction | No | | |

#### 97) N-acyl-lysophosphatidylethanolamine (LNAPE)[M-H]- / Lipid identification

| | | | |
|---|---|---|---|
| Lipid class | N-acyl-lysophosphatidylethanolamine (LNAPE) | Background check at MS2 | No |
| Derivatization | - | Did you presume assumptions for identification? | No |
| MS Level for identification | MS1, MS2 | Check isomer overlap | No |
| Identification level | Molecular species level | RT verified by standard | Yes |
| Polarity mode | Negative | Separation of isobaric/isomeric interferece confirmed | Yes |
| Type of negative (precursor)ion | [M-H]- | Model for separation prediction | Yes |
| Fragments for identification | | Additional dimension/techniques | - |

| Fragment name |
|---|
| Phosphoglycerol -H2O fragment |
| Fatty acid fragment |
| Neutral loss of Acyl |
| Neutral loss of Acyl and H2O |

| | | | |
|---|---|---|---|
| Isotope correction at MS1 | No | Lipid Identification Software | MS-DIAL |
| Isotope correction at MS2 | No | Data manipulation | Smoothing, Centroiding |
| MS1 verified by standard | No | Nomenclature for intact lipid molecule | Yes |
| MS2 verified by standard | No | Nomenclature for fragment ions | No |
| Background check at MS1 | Yes | Further identification remarks | - |

#### 97) N-acyl-lysophosphatidylethanolamine (LNAPE)[M-H]- / Lipid quantification

| | | | |
|---|---|---|---|
| Quantitative | Yes | Limit of quantification | No |
| MS Level for quantification | MS1 | Normalization to reference | No |
| Internal lipid standard(s) MS1 | | Lipid Quantification Software | MS-DIAL |

| Internal standard | Endogenous subclass |
|---|---|
| PE 15:0_18:1(d7) | LNAPE subclass |

| | | | |
|---|---|---|---|
| Type of quantification | Internal standard amount | Batch correction | No |
| Response correction | No | Further quantification remarks | - |
| Type I isotope correction | No | | |

#### 98) N-acyl-lysophosphatidylserine (LNAPS)[M-H]- / Lipid identification

| | | | |
|---|---|---|---|
| Lipid class | N-acyl-lysophosphatidylserine (LNAPS) | Background check at MS2 | No |
| Derivatization | - | Did you presume assumptions for identification? | No |
| MS Level for identification | MS1, MS2 | Check isomer overlap | No |
| Identification level | Molecular species level | RT verified by standard | Yes |
| Polarity mode | Negative | Separation of isobaric/isomeric interferece confirmed | Yes |
| Type of negative (precursor)ion | [M-H]- | Model for separation prediction | Yes |
| Fragments for identification | | Additional dimension/techniques | - |

| Fragment name |
|---|
| Phosphoglycerol -H2O fragment |
| Neutral loss of Acyl and C3H5NO2 |

| | | | |
|---|---|---|---|
| Isotope correction at MS1 | No | Lipid Identification Software | MS-DIAL |
| Isotope correction at MS2 | No | Data manipulation | Smoothing, Centroiding |
| MS1 verified by standard | No | Nomenclature for intact lipid molecule | Yes |
| MS2 verified by standard | No | Nomenclature for fragment ions | No |
| Background check at MS1 | Yes | Further identification remarks | - |

#### 98) N-acyl-lysophosphatidylserine (LNAPS)[M-H]- / Lipid quantification

| | | | |
|---|---|---|---|
| Quantitative | Yes | Limit of quantification | No |
| MS Level for quantification | MS1 | Normalization to reference | No |
| Internal lipid standard(s) MS1 | | Lipid Quantification Software | MS-DIAL |

| Internal standard | Endogenous subclass |
|---|---|
| PS 15:0_18:1(d7) | LNAPS subclass |

| | | | |
|---|---|---|---|
| Type of quantification | Internal standard amount | Batch correction | No |
| Response correction | No | Further quantification remarks | - |
| Type I isotope correction | No | | |

#### 99) DMPE[M-H]- / Lipid identification

| | | | |
|---|---|---|---|
| Lipid class | DMPE | Background check at MS2 | No |
| Derivatization | - | Did you presume assumptions for identification? | No |
| MS Level for identification | MS1, MS2 | Check isomer overlap | No |
| Identification level | Molecular species level | RT verified by standard | Yes |
| Polarity mode | Negative | Separation of isobaric/isomeric interferece confirmed | Yes |
| Type of negative (precursor)ion | [M-H]- | Model for separation prediction | Yes |
| Fragments for identification | | Additional dimension/techniques | - |

| Fragment name |
|---|
| Fatty acid fragment |

| | | | |
|---|---|---|---|
| Isotope correction at MS1 | No | Lipid Identification Software | MS-DIAL |
| Isotope correction at MS2 | No | Data manipulation | Smoothing, Centroiding |
| MS1 verified by standard | No | Nomenclature for intact lipid molecule | Yes |
| MS2 verified by standard | No | Nomenclature for fragment ions | No |
| Background check at MS1 | Yes | Further identification remarks | - |

#### 99) DMPE[M-H]- / Lipid quantification

| | | | |
|---|---|---|---|
| Quantitative | Yes | Limit of quantification | No |
| MS Level for quantification | MS1 | Normalization to reference | No |
| Internal lipid standard(s) MS1 | | Lipid Quantification Software | MS-DIAL |

| Internal standard | Endogenous subclass |
|---|---|
| PE 15:0_18:1(d7) | DMPE subclass |

| | | | |
|---|---|---|---|
| Type of quantification | Internal standard amount | Batch correction | No |
| Response correction | No | Further quantification remarks | - |
| Type I isotope correction | No | | |

#### 100) NGcGM3 (NGcGM3)[M-H]- / Lipid identification

| | | | |
|---|---|---|---|
| Lipid class | NGcGM3 (NGcGM3) | Background check at MS2 | No |
| Derivatization | - | Did you presume assumptions for identification? | No |
| MS Level for identification | MS1, MS2 | Check isomer overlap | No |
| Identification level | Molecular species level | RT verified by standard | Yes |
| Polarity mode | Negative | Separation of isobaric/isomeric interferece confirmed | Yes |
| Type of negative (precursor)ion | [M-H]- | Model for separation prediction | Yes |
| Fragments for identification | | Additional dimension/techniques | - |

| Fragment name |
|---|
| Characteristic fragment (C11H16NO9-) |

| | | | |
|---|---|---|---|
| Isotope correction at MS1 | No | Lipid Identification Software | MS-DIAL |
| Isotope correction at MS2 | No | Data manipulation | Smoothing, Centroiding |
| MS1 verified by standard | No | Nomenclature for intact lipid molecule | Yes |
| MS2 verified by standard | No | Nomenclature for fragment ions | No |
| Background check at MS1 | Yes | Further identification remarks | - |

#### 100) NGcGM3 (NGcGM3)[M-H]- / Lipid quantification

| | | | |
|---|---|---|---|
| Quantitative | Yes | Limit of quantification | No |
| MS Level for quantification | MS1 | Normalization to reference | No |
| Internal lipid standard(s) MS1 | | Lipid Quantification Software | MS-DIAL |

| Internal standard | Endogenous subclass |
|---|---|
| LPC 18:1(d7) | NGcGM3 subclass |

| | | | |
|---|---|---|---|
| Type of quantification | Internal standard amount | Batch correction | No |
| Response correction | No | Further quantification remarks | - |
| Type I isotope correction | No | | |

#### 101) MMPE[M-H]- / Lipid identification

| | | | |
|---|---|---|---|
| Lipid class | MMPE | Background check at MS2 | No |
| Derivatization | - | Did you presume assumptions for identification? | No |
| MS Level for identification | MS1, MS2 | Check isomer overlap | No |
| Identification level | Molecular species level | RT verified by standard | Yes |
| Polarity mode | Negative | Separation of isobaric/isomeric interferece confirmed | Yes |
| Type of negative (precursor)ion | [M-H]- | Model for separation prediction | Yes |
| Fragments for identification | | Additional dimension/techniques | - |

| Fragment name |
|---|
| Fatty acid fragment |

| | | | |
|---|---|---|---|
| Isotope correction at MS1 | No | Lipid Identification Software | MS-DIAL |
| Isotope correction at MS2 | No | Data manipulation | Smoothing, Centroiding |
| MS1 verified by standard | No | Nomenclature for intact lipid molecule | Yes |
| MS2 verified by standard | No | Nomenclature for fragment ions | No |
| Background check at MS1 | Yes | Further identification remarks | - |

#### 101) MMPE[M-H]- / Lipid quantification

| | | | |
|---|---|---|---|
| Quantitative | Yes | Limit of quantification | No |
| MS Level for quantification | MS1 | Normalization to reference | No |
| Internal lipid standard(s) MS1 | | Lipid Quantification Software | MS-DIAL |

| Internal standard | Endogenous subclass |
|---|---|
| PE 15:0_18:1(d7) | MMPE subclass |

| | | | |
|---|---|---|---|
| Type of quantification | Internal standard amount | Batch correction | No |
| Response correction | No | Further quantification remarks | - |
| Type I isotope correction | No | | |

#### 102) Oxidized fatty acid (OxFA)[M-H]- / Lipid identification

| | | | |
|---|---|---|---|
| Lipid class | Oxidized fatty acid (OxFA) | Background check at MS2 | No |
| Derivatization | - | Did you presume assumptions for identification? | No |
| MS Level for identification | MS1, MS2 | Check isomer overlap | No |
| Identification level | Species level | RT verified by standard | Yes |
| Polarity mode | Negative | Separation of isobaric/isomeric interferece confirmed | Yes |
| Type of negative (precursor)ion | [M-H]- | Model for separation prediction | Yes |
| Fragments for identification | | Additional dimension/techniques | - |

| Fragment name |
|---|
| Neutral loss of H2O |

| | | | |
|---|---|---|---|
| Isotope correction at MS1 | No | Lipid Identification Software | MS-DIAL |
| Isotope correction at MS2 | No | Data manipulation | Smoothing, Centroiding |
| MS1 verified by standard | No | Nomenclature for intact lipid molecule | Yes |
| MS2 verified by standard | No | Nomenclature for fragment ions | No |
| Background check at MS1 | Yes | Further identification remarks | - |

#### 102) Oxidized fatty acid (OxFA)[M-H]- / Lipid quantification

| | | | |
|---|---|---|---|
| Quantitative | Yes | Limit of quantification | No |
| MS Level for quantification | MS1 | Normalization to reference | No |
| Internal lipid standard(s) MS1 | | Lipid Quantification Software | MS-DIAL |

| Internal standard | Endogenous subclass |
|---|---|
| FA 18:0(d3) | OxFA subclass |

| | | | |
|---|---|---|---|
| Type of quantification | Internal standard amount | Batch correction | No |
| Response correction | No | Further quantification remarks | - |
| Type I isotope correction | No | | |

#### 103) Oxidized phosphatidylcholine (OxPC)[M+CH3COO]- / Lipid identification

| | | | |
|---|---|---|---|
| Lipid class | Oxidized phosphatidylcholine (OxPC) | Background check at MS2 | No |
| Derivatization | - | Did you presume assumptions for identification? | No |
| MS Level for identification | MS1, MS2 | Check isomer overlap | No |
| Identification level | Molecular species level | RT verified by standard | Yes |
| Polarity mode | Negative | Separation of isobaric/isomeric interferece confirmed | Yes |
| Type of negative (precursor)ion | [M+CH3COO]- | Model for separation prediction | Yes |
| Fragments for identification | | Additional dimension/techniques | - |

| Fragment name |
|---|
| Neutral loss of methyl moiety |
| Fatty acid fragment |
| Oxidized fatty acid fragment |
| Oxidized fatty acid -H2O fragment |

| | | | |
|---|---|---|---|
| Isotope correction at MS1 | No | Lipid Identification Software | MS-DIAL |
| Isotope correction at MS2 | No | Data manipulation | Smoothing, Centroiding |
| MS1 verified by standard | No | Nomenclature for intact lipid molecule | Yes |
| MS2 verified by standard | No | Nomenclature for fragment ions | No |
| Background check at MS1 | Yes | Further identification remarks | - |

#### 103) Oxidized phosphatidylcholine (OxPC)[M+CH3COO]- / Lipid quantification

| | | | |
|---|---|---|---|
| Quantitative | Yes | Limit of quantification | No |
| MS Level for quantification | MS1 | Normalization to reference | No |
| Internal lipid standard(s) MS1 | | Lipid Quantification Software | MS-DIAL |

| Internal standard | Endogenous subclass |
|---|---|
| PC 15:0_18:1(d7) | OxPC subclass |

| | | | |
|---|---|---|---|
| Type of quantification | Internal standard amount | Batch correction | No |
| Response correction | No | Further quantification remarks | - |
| Type I isotope correction | No | | |

#### 104) Oxidized phosphatidylethanolamine (OxPE)[M-H]- / Lipid identification

| | | | |
|---|---|---|---|
| Lipid class | Oxidized phosphatidylethanolamine (OxPE) | Background check at MS2 | No |
| Derivatization | - | Did you presume assumptions for identification? | No |
| MS Level for identification | MS1, MS2 | Check isomer overlap | No |
| Identification level | Molecular species level | RT verified by standard | Yes |
| Polarity mode | Negative | Separation of isobaric/isomeric interferece confirmed | Yes |
| Type of negative (precursor)ion | [M-H]- | Model for separation prediction | Yes |
| Fragments for identification | | Additional dimension/techniques | - |

| Fragment name |
|---|
| Neutral loss of H2O |
| Fatty acid fragment |
| Oxidized fatty acid fragment |
| Oxidized fatty acid -H2O fragment |

| | | | |
|---|---|---|---|
| Isotope correction at MS1 | No | Lipid Identification Software | MS-DIAL |
| Isotope correction at MS2 | No | Data manipulation | Smoothing, Centroiding |
| MS1 verified by standard | No | Nomenclature for intact lipid molecule | Yes |
| MS2 verified by standard | No | Nomenclature for fragment ions | No |
| Background check at MS1 | Yes | Further identification remarks | - |

#### 104) Oxidized phosphatidylethanolamine (OxPE)[M-H]- / Lipid quantification

| | | | |
|---|---|---|---|
| Quantitative | Yes | Limit of quantification | No |
| MS Level for quantification | MS1 | Normalization to reference | No |
| Internal lipid standard(s) MS1 | | Lipid Quantification Software | MS-DIAL |

| Internal standard | Endogenous subclass |
|---|---|
| PE 15:0_18:1(d7) | OxPE subclass |

| | | | |
|---|---|---|---|
| Type of quantification | Internal standard amount | Batch correction | No |
| Response correction | No | Further quantification remarks | - |
| Type I isotope correction | No | | |

#### 105) Oxidized phosphatidylglycerol (OxPG)[M-H]- / Lipid identification

| | | | |
|---|---|---|---|
| Lipid class | Oxidized phosphatidylglycerol (OxPG) | Background check at MS2 | No |
| Derivatization | - | Did you presume assumptions for identification? | No |
| MS Level for identification | MS1, MS2 | Check isomer overlap | No |
| Identification level | Molecular species level | RT verified by standard | Yes |
| Polarity mode | Negative | Separation of isobaric/isomeric interferece confirmed | Yes |
| Type of negative (precursor)ion | [M-H]- | Model for separation prediction | Yes |
| Fragments for identification | | Additional dimension/techniques | - |

| Fragment name |
|---|
| Fatty acid fragment |
| Oxidized fatty acid fragment |
| Oxidized fatty acid -H2O fragment |

| | | | |
|---|---|---|---|
| Isotope correction at MS1 | No | Lipid Identification Software | MS-DIAL |
| Isotope correction at MS2 | No | Data manipulation | Smoothing, Centroiding |
| MS1 verified by standard | No | Nomenclature for intact lipid molecule | Yes |
| MS2 verified by standard | No | Nomenclature for fragment ions | No |
| Background check at MS1 | Yes | Further identification remarks | - |

#### 105) Oxidized phosphatidylglycerol (OxPG)[M-H]- / Lipid quantification

| | | | |
|---|---|---|---|
| Quantitative | Yes | Limit of quantification | No |
| MS Level for quantification | MS1 | Normalization to reference | No |
| Internal lipid standard(s) MS1 | | Lipid Quantification Software | MS-DIAL |

| Internal standard | Endogenous subclass |
|---|---|
| PG 15:0_18:1(d7) | OxPG subclass |

| | | | |
|---|---|---|---|
| Type of quantification | Internal standard amount | Batch correction | No |
| Response correction | No | Further quantification remarks | - |
| Type I isotope correction | No | | |

#### 106) Oxidized phosphatidylinositol (OxPI)[M-H]- / Lipid identification

| | | | |
|---|---|---|---|
| Lipid class | Oxidized phosphatidylinositol (OxPI) | Background check at MS2 | No |
| Derivatization | - | Did you presume assumptions for identification? | No |
| MS Level for identification | MS1, MS2 | Check isomer overlap | No |
| Identification level | Molecular species level | RT verified by standard | Yes |
| Polarity mode | Negative | Separation of isobaric/isomeric interferece confirmed | Yes |
| Type of negative (precursor)ion | [M-H]- | Model for separation prediction | Yes |
| Fragments for identification | | Additional dimension/techniques | - |

| Fragment name |
|---|
| Phosphoinositol -H2O fragment |
| Characteristic fragment (C9H14O9P-) |
| Neutral loss of H2O |
| Fatty acid fragment |
| Oxidized fatty acid fragment |
| Oxidized fatty acid -H2O fragment |

| | | | |
|---|---|---|---|
| Isotope correction at MS1 | No | Lipid Identification Software | MS-DIAL |
| Isotope correction at MS2 | No | Data manipulation | Smoothing, Centroiding |
| MS1 verified by standard | No | Nomenclature for intact lipid molecule | Yes |
| MS2 verified by standard | No | Nomenclature for fragment ions | No |
| Background check at MS1 | Yes | Further identification remarks | - |

#### 106) Oxidized phosphatidylinositol (OxPI)[M-H]- / Lipid quantification

| | | | |
|---|---|---|---|
| Quantitative | Yes | Limit of quantification | No |
| MS Level for quantification | MS1 | Normalization to reference | No |
| Internal lipid standard(s) MS1 | | Lipid Quantification Software | MS-DIAL |

| Internal standard | Endogenous subclass |
|---|---|
| PI 15:0_18:1(d7) | OxPI subclass |

| | | | |
|---|---|---|---|
| Type of quantification | Internal standard amount | Batch correction | No |
| Response correction | No | Further quantification remarks | - |
| Type I isotope correction | No | | |

#### 107) Oxidized phosphatidylserine (OxPS)[M-H]- / Lipid identification

| | | | |
|---|---|---|---|
| Lipid class | Oxidized phosphatidylserine (OxPS) | Background check at MS2 | No |
| Derivatization | - | Did you presume assumptions for identification? | No |
| MS Level for identification | MS1, MS2 | Check isomer overlap | No |
| Identification level | Molecular species level | RT verified by standard | Yes |
| Polarity mode | Negative | Separation of isobaric/isomeric interferece confirmed | Yes |
| Type of negative (precursor)ion | [M-H]- | Model for separation prediction | Yes |
| Fragments for identification | | Additional dimension/techniques | - |

| Fragment name |
|---|
| Neutral loss of C3H6NO2 |
| Neutral loss of H2O |
| Fatty acid fragment |
| Oxidized fatty acid fragment |
| Oxidized fatty acid -H2O fragment |

| | | | |
|---|---|---|---|
| Isotope correction at MS1 | No | Lipid Identification Software | MS-DIAL |
| Isotope correction at MS2 | No | Data manipulation | Smoothing, Centroiding |
| MS1 verified by standard | No | Nomenclature for intact lipid molecule | Yes |
| MS2 verified by standard | No | Nomenclature for fragment ions | No |
| Background check at MS1 | Yes | Further identification remarks | - |

#### 107) Oxidized phosphatidylserine (OxPS)[M-H]- / Lipid quantification

| | | | |
|---|---|---|---|
| Quantitative | Yes | Limit of quantification | No |
| MS Level for quantification | MS1 | Normalization to reference | No |
| Internal lipid standard(s) MS1 | | Lipid Quantification Software | MS-DIAL |

| Internal standard | Endogenous subclass |
|---|---|
| PS 15:0_18:1(d7) | OxPS subclass |

| | | | |
|---|---|---|---|
| Type of quantification | Internal standard amount | Batch correction | No |
| Response correction | No | Further quantification remarks | - |
| Type I isotope correction | No | | |

#### 108) Oxidized triglyceride (OxTG)[M+NH4]+ / Lipid identification

| | | | |
|---|---|---|---|
| Lipid class | Oxidized triglyceride (OxTG) | Background check at MS2 | No |
| Derivatization | - | Did you presume assumptions for identification? | No |
| MS Level for identification | MS1, MS2 | Check isomer overlap | No |
| Identification level | Molecular species level | RT verified by standard | Yes |
| Polarity mode | Positive | Separation of isobaric/isomeric interferece confirmed | Yes |
| Type of positive (precursor)ion | [M+NH4]+ | Model for separation prediction | Yes |
| Fragments for identification | | Additional dimension/techniques | - |

| Fragment name |
|---|
| Neutral loss of acyl and H2O |
| Neutral loss of acyl and 2H2O |
| Neutral loss of acyl and H2O and O |

| | | | |
|---|---|---|---|
| Isotope correction at MS1 | No | Lipid Identification Software | MS-DIAL |
| Isotope correction at MS2 | No | Data manipulation | Smoothing, Centroiding |
| MS1 verified by standard | No | Nomenclature for intact lipid molecule | Yes |
| MS2 verified by standard | No | Nomenclature for fragment ions | No |
| Background check at MS1 | Yes | Further identification remarks | - |

#### 108) Oxidized triglyceride (OxTG)[M+NH4]+ / Lipid quantification

| | | | |
|---|---|---|---|
| Quantitative | Yes | Limit of quantification | No |
| MS Level for quantification | MS1 | Normalization to reference | No |
| Internal lipid standard(s) MS1 | | Lipid Quantification Software | MS-DIAL |

| Internal standard | Endogenous subclass |
|---|---|
| TG 15:0_18:1(d7)_15:0 | OxTG subclass |

| | | | |
|---|---|---|---|
| Type of quantification | Internal standard amount | Batch correction | No |
| Response correction | No | Further quantification remarks | - |
| Type I isotope correction | No | | |

#### 109) PA[M-H]- / Lipid identification

| | | | |
|---|---|---|---|
| Lipid class | PA | Background check at MS2 | No |
| Derivatization | - | Did you presume assumptions for identification? | No |
| MS Level for identification | MS1, MS2 | Check isomer overlap | No |
| Identification level | Molecular species level | RT verified by standard | Yes |
| Polarity mode | Negative | Separation of isobaric/isomeric interferece confirmed | Yes |
| Type of negative (precursor)ion | [M-H]- | Model for separation prediction | Yes |
| Fragments for identification | | Additional dimension/techniques | - |

| Fragment name |
|---|
| Phosphoglycerol -H2O fragment |
| Fatty acid fragment |

| | | | |
|---|---|---|---|
| Isotope correction at MS1 | No | Lipid Identification Software | MS-DIAL |
| Isotope correction at MS2 | No | Data manipulation | Smoothing, Centroiding |
| MS1 verified by standard | No | Nomenclature for intact lipid molecule | Yes |
| MS2 verified by standard | No | Nomenclature for fragment ions | No |
| Background check at MS1 | Yes | Further identification remarks | - |

#### 109) PA[M-H]- / Lipid quantification

| | | | |
|---|---|---|---|
| Quantitative | Yes | Limit of quantification | No |
| MS Level for quantification | MS1 | Normalization to reference | No |
| Internal lipid standard(s) MS1 | | Lipid Quantification Software | MS-DIAL |

| Internal standard | Endogenous subclass |
|---|---|
| LPC 18:1(d7) | PA subclass |

| | | | |
|---|---|---|---|
| Type of quantification | Internal standard amount | Batch correction | No |
| Response correction | No | Further quantification remarks | - |
| Type I isotope correction | No | | |

#### 110) PC[M+CH3COO]- / Lipid identification

| | | | |
|---|---|---|---|
| Lipid class | PC | Background check at MS2 | No |
| Derivatization | - | Did you presume assumptions for identification? | No |
| MS Level for identification | MS1, MS2 | Check isomer overlap | No |
| Identification level | Molecular species level | RT verified by standard | Yes |
| Polarity mode | Negative | Separation of isobaric/isomeric interferece confirmed | Yes |
| Type of negative (precursor)ion | [M+CH3COO]- | Model for separation prediction | Yes |
| Fragments for identification | | Additional dimension/techniques | - |

| Fragment name |
|---|
| Neutral loss of methyl moiety |
| Fatty acid fragment |

| | | | |
|---|---|---|---|
| Isotope correction at MS1 | No | Lipid Identification Software | MS-DIAL |
| Isotope correction at MS2 | No | Data manipulation | Smoothing, Centroiding |
| MS1 verified by standard | Yes | Nomenclature for intact lipid molecule | Yes |
| MS2 verified by standard | Yes | Nomenclature for fragment ions | No |
| Background check at MS1 | Yes | Further identification remarks | - |

#### 110) PC[M+CH3COO]- / Lipid quantification

| | | | |
|---|---|---|---|
| Quantitative | Yes | Limit of quantification | No |
| MS Level for quantification | MS1 | Normalization to reference | No |
| Internal lipid standard(s) MS1 | | Lipid Quantification Software | MS-DIAL |

| Internal standard | Endogenous subclass |
|---|---|
| PC 15:0_18:1(d7) | PC subclass |

| | | | |
|---|---|---|---|
| Type of quantification | Internal standard amount | Batch correction | No |
| Response correction | No | Further quantification remarks | - |
| Type I isotope correction | No | | |

#### 111) Phosphatidylethanol (PEtOH)[M-H]- / Lipid identification

| | | | |
|---|---|---|---|
| Lipid class | Phosphatidylethanol (PEtOH) | Background check at MS2 | No |
| Derivatization | - | Did you presume assumptions for identification? | No |
| MS Level for identification | MS1, MS2 | Check isomer overlap | No |
| Identification level | Molecular species level | RT verified by standard | Yes |
| Polarity mode | Negative | Separation of isobaric/isomeric interferece confirmed | Yes |
| Type of negative (precursor)ion | [M-H]- | Model for separation prediction | Yes |
| Fragments for identification | | Additional dimension/techniques | - |

| Fragment name |
|---|
| Phosphoethanol |
| Fatty acid fragment |

| | | | |
|---|---|---|---|
| Isotope correction at MS1 | No | Lipid Identification Software | MS-DIAL |
| Isotope correction at MS2 | No | Data manipulation | Smoothing, Centroiding |
| MS1 verified by standard | No | Nomenclature for intact lipid molecule | Yes |
| MS2 verified by standard | No | Nomenclature for fragment ions | No |
| Background check at MS1 | Yes | Further identification remarks | - |

#### 111) Phosphatidylethanol (PEtOH)[M-H]- / Lipid quantification

| | | | |
|---|---|---|---|
| Quantitative | Yes | Limit of quantification | No |
| MS Level for quantification | MS1 | Normalization to reference | No |
| Internal lipid standard(s) MS1 | | Lipid Quantification Software | MS-DIAL |

| Internal standard | Endogenous subclass |
|---|---|
| LPC 18:1(d7) | PEtOH subclass |

| | | | |
|---|---|---|---|
| Type of quantification | Internal standard amount | Batch correction | No |
| Response correction | No | Further quantification remarks | - |
| Type I isotope correction | No | | |

#### 112) PE[M-H]- / Lipid identification

| | | | |
|---|---|---|---|
| Lipid class | PE | Background check at MS2 | No |
| Derivatization | - | Did you presume assumptions for identification? | No |
| MS Level for identification | MS1, MS2 | Check isomer overlap | No |
| Identification level | Molecular species level | RT verified by standard | Yes |
| Polarity mode | Negative | Separation of isobaric/isomeric interferece confirmed | Yes |
| Type of negative (precursor)ion | [M-H]- | Model for separation prediction | Yes |
| Fragments for identification | | Additional dimension/techniques | - |

| Fragment name |
|---|
| Characteristic fragment (C5H11NO5P-) |
| Fatty acid fragment |

| | | | |
|---|---|---|---|
| Isotope correction at MS1 | No | Lipid Identification Software | MS-DIAL |
| Isotope correction at MS2 | No | Data manipulation | Smoothing, Centroiding |
| MS1 verified by standard | Yes | Nomenclature for intact lipid molecule | Yes |
| MS2 verified by standard | Yes | Nomenclature for fragment ions | No |
| Background check at MS1 | Yes | Further identification remarks | - |

#### 112) PE[M-H]- / Lipid quantification

| | | | |
|---|---|---|---|
| Quantitative | Yes | Limit of quantification | No |
| MS Level for quantification | MS1 | Normalization to reference | No |
| Internal lipid standard(s) MS1 | | Lipid Quantification Software | MS-DIAL |

| Internal standard | Endogenous subclass |
|---|---|
| PE 15:0_18:1(d7) | PE subclass |

| | | | |
|---|---|---|---|
| Type of quantification | Internal standard amount | Batch correction | No |
| Response correction | No | Further quantification remarks | - |
| Type I isotope correction | No | | |

#### 113) PG[M-H]- / Lipid identification

| | | | |
|---|---|---|---|
| Lipid class | PG | Background check at MS2 | No |
| Derivatization | - | Did you presume assumptions for identification? | No |
| MS Level for identification | MS1, MS2 | Check isomer overlap | No |
| Identification level | Molecular species level | RT verified by standard | Yes |
| Polarity mode | Negative | Separation of isobaric/isomeric interferece confirmed | Yes |
| Type of negative (precursor)ion | [M-H]- | Model for separation prediction | Yes |
| Fragments for identification | | Additional dimension/techniques | - |

| Fragment name |
|---|
| Phosphoglycerol -H2O |
| Fatty acid fragment |

| | | | |
|---|---|---|---|
| Isotope correction at MS1 | No | Lipid Identification Software | MS-DIAL |
| Isotope correction at MS2 | No | Data manipulation | Smoothing, Centroiding |
| MS1 verified by standard | Yes | Nomenclature for intact lipid molecule | Yes |
| MS2 verified by standard | Yes | Nomenclature for fragment ions | No |
| Background check at MS1 | Yes | Further identification remarks | - |

#### 113) PG[M-H]- / Lipid quantification

| | | | |
|---|---|---|---|
| Quantitative | Yes | Limit of quantification | No |
| MS Level for quantification | MS1 | Normalization to reference | No |
| Internal lipid standard(s) MS1 | | Lipid Quantification Software | MS-DIAL |

| Internal standard | Endogenous subclass |
|---|---|
| PG 15:0_18:1(d7) | PG subclass |

| | | | |
|---|---|---|---|
| Type of quantification | Internal standard amount | Batch correction | No |
| Response correction | No | Further quantification remarks | - |
| Type I isotope correction | No | | |

#### 114) PI[M-H]- / Lipid identification

| | | | |
|---|---|---|---|
| Lipid class | PI | Background check at MS2 | No |
| Derivatization | - | Did you presume assumptions for identification? | No |
| MS Level for identification | MS1, MS2 | Check isomer overlap | No |
| Identification level | Molecular species level | RT verified by standard | Yes |
| Polarity mode | Negative | Separation of isobaric/isomeric interferece confirmed | Yes |
| Type of negative (precursor)ion | [M-H]- | Model for separation prediction | Yes |
| Fragments for identification | | Additional dimension/techniques | - |

| Fragment name |
|---|
| Phosphoinositol -H2O |
| Characteristic fragment (C9H14O9P-) |
| Fatty acid fragment |

| | | | |
|---|---|---|---|
| Isotope correction at MS1 | No | Lipid Identification Software | MS-DIAL |
| Isotope correction at MS2 | No | Data manipulation | Smoothing, Centroiding |
| MS1 verified by standard | Yes | Nomenclature for intact lipid molecule | Yes |
| MS2 verified by standard | Yes | Nomenclature for fragment ions | No |
| Background check at MS1 | Yes | Further identification remarks | - |

#### 114) PI[M-H]- / Lipid quantification

| | | | |
|---|---|---|---|
| Quantitative | Yes | Limit of quantification | No |
| MS Level for quantification | MS1 | Normalization to reference | No |
| Internal lipid standard(s) MS1 | | Lipid Quantification Software | MS-DIAL |

| Internal standard | Endogenous subclass |
|---|---|
| PI 15:0_18:1(d7) | PI subclass |

| | | | |
|---|---|---|---|
| Type of quantification | Internal standard amount | Batch correction | No |
| Response correction | No | Further quantification remarks | - |
| Type I isotope correction | No | | |

#### 115) Phosphatidylmethanol (PMeOH)[M-H]- / Lipid identification

| | | | |
|---|---|---|---|
| Lipid class | Phosphatidylmethanol (PMeOH) | Background check at MS2 | No |
| Derivatization | - | Did you presume assumptions for identification? | No |
| MS Level for identification | MS1, MS2 | Check isomer overlap | No |
| Identification level | Molecular species level | RT verified by standard | Yes |
| Polarity mode | Negative | Separation of isobaric/isomeric interferece confirmed | Yes |
| Type of negative (precursor)ion | [M-H]- | Model for separation prediction | Yes |
| Fragments for identification | | Additional dimension/techniques | - |

| Fragment name |
|---|
| Phosphomethanol |
| Fatty acid fragment |

| | | | |
|---|---|---|---|
| Isotope correction at MS1 | No | Lipid Identification Software | MS-DIAL |
| Isotope correction at MS2 | No | Data manipulation | Smoothing, Centroiding |
| MS1 verified by standard | No | Nomenclature for intact lipid molecule | Yes |
| MS2 verified by standard | No | Nomenclature for fragment ions | No |
| Background check at MS1 | Yes | Further identification remarks | - |

#### 115) Phosphatidylmethanol (PMeOH)[M-H]- / Lipid quantification

| | | | |
|---|---|---|---|
| Quantitative | Yes | Limit of quantification | No |
| MS Level for quantification | MS1 | Normalization to reference | No |
| Internal lipid standard(s) MS1 | | Lipid Quantification Software | MS-DIAL |

| Internal standard | Endogenous subclass |
|---|---|
| LPC 18:1(d7) | PMeOH subclass |

| | | | |
|---|---|---|---|
| Type of quantification | Internal standard amount | Batch correction | No |
| Response correction | No | Further quantification remarks | - |
| Type I isotope correction | No | | |

#### 116) PS[M-H]- / Lipid identification

| | | | |
|---|---|---|---|
| Lipid class | PS | Background check at MS2 | No |
| Derivatization | - | Did you presume assumptions for identification? | No |
| MS Level for identification | MS1, MS2 | Check isomer overlap | No |
| Identification level | Molecular species level | RT verified by standard | Yes |
| Polarity mode | Negative | Separation of isobaric/isomeric interferece confirmed | Yes |
| Type of negative (precursor)ion | [M-H]- | Model for separation prediction | Yes |
| Fragments for identification | | Additional dimension/techniques | - |

| Fragment name |
|---|
| Neutral loss of C3H6NO2 |
| Fatty acid fragment |

| | | | |
|---|---|---|---|
| Isotope correction at MS1 | No | Lipid Identification Software | MS-DIAL |
| Isotope correction at MS2 | No | Data manipulation | Smoothing, Centroiding |
| MS1 verified by standard | Yes | Nomenclature for intact lipid molecule | Yes |
| MS2 verified by standard | Yes | Nomenclature for fragment ions | No |
| Background check at MS1 | Yes | Further identification remarks | - |

#### 116) PS[M-H]- / Lipid quantification

| | | | |
|---|---|---|---|
| Quantitative | Yes | Limit of quantification | No |
| MS Level for quantification | MS1 | Normalization to reference | No |
| Internal lipid standard(s) MS1 | | Lipid Quantification Software | MS-DIAL |

| Internal standard | Endogenous subclass |
|---|---|
| PS 15:0_18:1(d7) | PS subclass |

| | | | |
|---|---|---|---|
| Type of quantification | Internal standard amount | Batch correction | No |
| Response correction | No | Further quantification remarks | - |
| Type I isotope correction | No | | |

#### 117) Phytosphingosine (PhytoSph)[M+H]+ / Lipid identification

| | | | |
|---|---|---|---|
| Lipid class | Phytosphingosine (PhytoSph) | Background check at MS2 | No |
| Derivatization | - | Did you presume assumptions for identification? | No |
| MS Level for identification | MS1, MS2 | Check isomer overlap | No |
| Identification level | Molecular species level | RT verified by standard | Yes |
| Polarity mode | Positive | Separation of isobaric/isomeric interferece confirmed | Yes |
| Type of positive (precursor)ion | [M+H]+ | Model for separation prediction | Yes |
| Fragments for identification | | Additional dimension/techniques | - |

| Fragment name |
|---|
| Neutral loss of $H_2O$ |
| Neutral loss of $2H_2O$ |
| Neutral loss of $3H_2O$ |
| Neutral loss of $CH_4O_2$ |

| | | | |
|---|---|---|---|
| Isotope correction at MS1 | No | Lipid Identification Software | MS-DIAL |
| Isotope correction at MS2 | No | Data manipulation | Smoothing, Centroiding |
| MS1 verified by standard | No | Nomenclature for intact lipid molecule | Yes |
| MS2 verified by standard | No | Nomenclature for fragment ions | No |
| Background check at MS1 | Yes | Further identification remarks | - |

#### 117) Phytosphingosine (PhytoSph)[M+H]+ / Lipid quantification

| | | | |
|---|---|---|---|
| Quantitative | Yes | Limit of quantification | No |
| MS Level for quantification | MS1 | Normalization to reference | No |
| Internal lipid standard(s) MS1 | | Lipid Quantification Software | MS-DIAL |

| Internal standard | Endogenous subclass |
|---|---|
| Cer 18:1;2O/15:0(d7) | PhytoSph subclass |

| | | | |
|---|---|---|---|
| Type of quantification | Internal standard amount | Batch correction | No |
| Response correction | No | Further quantification remarks | - |
| Type I isotope correction | No | | |

#### 118) Semino lipid (EtherSMGDG)[M-H]- / Lipid identification

| | | | |
|---|---|---|---|
| Lipid class | Semino lipid (EtherSMGDG) | Background check at MS2 | No |
| Derivatization | - | Did you presume assumptions for identification? | No |
| MS Level for identification | MS1, MS2 | Check isomer overlap | No |
| Identification level | Molecular species level | RT verified by standard | Yes |
| Polarity mode | Negative | Separation of isobaric/isomeric interferece confirmed | Yes |
| Type of negative (precursor)ion | [M-H]- | Model for separation prediction | Yes |
| Fragments for identification | | Additional dimension/techniques | - |

| Fragment name |
|---|
| Sulfate |
| Hexosylsulfate |
| Neutral loss of acyl and H2O |

| | | | |
|---|---|---|---|
| Isotope correction at MS1 | No | Lipid Identification Software | MS-DIAL |
| Isotope correction at MS2 | No | Data manipulation | Smoothing, Centroiding |
| MS1 verified by standard | No | Nomenclature for intact lipid molecule | Yes |
| MS2 verified by standard | No | Nomenclature for fragment ions | No |
| Background check at MS1 | Yes | Further identification remarks | - |

#### 118) Semino lipid (EtherSMGDG)[M-H]- / Lipid quantification

| | | | |
|---|---|---|---|
| Quantitative | Yes | Limit of quantification | No |
| MS Level for quantification | MS1 | Normalization to reference | No |
| Internal lipid standard(s) MS1 | | Lipid Quantification Software | MS-DIAL |

| Internal standard | Endogenous subclass |
|---|---|
| Cer 18:1;2O/15:0(d7) | EtherSMGDG subclass |

| | | | |
|---|---|---|---|
| Type of quantification | Internal standard amount | Batch correction | No |
| Response correction | No | Further quantification remarks | - |
| Type I isotope correction | No | | |

#### 119) Semino lipid (SMGDG)[M-H]- / Lipid identification

| | | | |
|---|---|---|---|
| Lipid class | Semino lipid (SMGDG) | Background check at MS2 | No |
| Derivatization | - | Did you presume assumptions for identification? | No |
| MS Level for identification | MS1, MS2 | Check isomer overlap | No |
| Identification level | Molecular species level | RT verified by standard | Yes |
| Polarity mode | Negative | Separation of isobaric/isomeric interferece confirmed | Yes |
| Type of negative (precursor)ion | [M-H]- | Model for separation prediction | Yes |
| Fragments for identification | | Additional dimension/techniques | - |

| Fragment name |
|---|
| Sulfate |
| Hexosylsulfate |
| Neutral loss of acyl and H2O |
| Fatty acid fragment |

| | | | |
|---|---|---|---|
| Isotope correction at MS1 | No | Lipid Identification Software | MS-DIAL |
| Isotope correction at MS2 | No | Data manipulation | Smoothing, Centroiding |
| MS1 verified by standard | No | Nomenclature for intact lipid molecule | Yes |
| MS2 verified by standard | No | Nomenclature for fragment ions | No |
| Background check at MS1 | Yes | Further identification remarks | - |

#### 119) Semino lipid (SMGDG)[M-H]- / Lipid quantification

| | | | |
|---|---|---|---|
| Quantitative | Yes | Limit of quantification | No |
| MS Level for quantification | MS1 | Normalization to reference | No |
| Internal lipid standard(s) MS1 | | Lipid Quantification Software | MS-DIAL |

| Internal standard | Endogenous subclass |
|---|---|
| Cer 18:1;2O/15:0(d7) | SMGDG subclass |

| | | | |
|---|---|---|---|
| Type of quantification | Internal standard amount | Batch correction | No |
| Response correction | No | Further quantification remarks | - |
| Type I isotope correction | No | | |

#### 120) Sitosterol ester (SISE)[M+NH4]+ / Lipid identification

| | | | |
|---|---|---|---|
| Lipid class | Sitosterol ester (SISE) | Background check at MS2 | No |
| Derivatization | - | Did you presume assumptions for identification? | No |
| MS Level for identification | MS1, MS2 | Check isomer overlap | No |
| Identification level | Molecular species level | RT verified by standard | Yes |
| Polarity mode | Positive | Separation of isobaric/isomeric interferece confirmed | Yes |
| Type of positive (precursor)ion | [M+NH4]+ | Model for separation prediction | Yes |
| Fragments for identification | | Additional dimension/techniques | - |

| Fragment name |
|---|
| Neutral loss of fatty acid |

| | | | |
|---|---|---|---|
| Isotope correction at MS1 | No | Lipid Identification Software | MS-DIAL |
| Isotope correction at MS2 | No | Data manipulation | Smoothing, Centroiding |
| MS1 verified by standard | No | Nomenclature for intact lipid molecule | Yes |
| MS2 verified by standard | No | Nomenclature for fragment ions | No |
| Background check at MS1 | Yes | Further identification remarks | - |

#### 120) Sitosterol ester (SISE)[M+NH4]+ / Lipid quantification

| | | | |
|---|---|---|---|
| Quantitative | Yes | Limit of quantification | No |
| MS Level for quantification | MS1 | Normalization to reference | No |
| Internal lipid standard(s) MS1 | | Lipid Quantification Software | MS-DIAL |

| Internal standard | Endogenous subclass |
|---|---|
| CE 18:1(d7) | SISE subclass |

| | | | |
|---|---|---|---|
| Type of quantification | Internal standard amount | Batch correction | No |
| Response correction | No | Further quantification remarks | - |
| Type I isotope correction | No | | |

#### 121) Sphinganine (DHSph)[M+H]+ / Lipid identification

| | | | |
|---|---|---|---|
| Lipid class | Sphinganine (DHSph) | Background check at MS2 | No |
| Derivatization | - | Did you presume assumptions for identification? | No |
| MS Level for identification | MS1, MS2 | Check isomer overlap | No |
| Identification level | Molecular species level | RT verified by standard | Yes |
| Polarity mode | Positive | Separation of isobaric/isomeric interferece confirmed | Yes |
| Type of positive (precursor)ion | [M+H]+ | Model for separation prediction | Yes |
| Fragments for identification | | Additional dimension/techniques | - |

| Fragment name |
|---|
| Neutral loss of H2O |
| Neutral loss of 2H2O |
| Neutral loss of CH4O2 |

| | | | |
|---|---|---|---|
| Isotope correction at MS1 | No | Lipid Identification Software | MS-DIAL |
| Isotope correction at MS2 | No | Data manipulation | Smoothing, Centroiding |
| MS1 verified by standard | No | Nomenclature for intact lipid molecule | Yes |
| MS2 verified by standard | No | Nomenclature for fragment ions | No |
| Background check at MS1 | Yes | Further identification remarks | - |

#### 121) Sphinganine (DHSph)[M+H]+ / Lipid quantification

| | | | |
|---|---|---|---|
| Quantitative | Yes | Limit of quantification | No |
| MS Level for quantification | MS1 | Normalization to reference | No |
| Internal lipid standard(s) MS1 | | Lipid Quantification Software | MS-DIAL |

| Internal standard | Endogenous subclass |
|---|---|
| Cer 18:1;2O/15:0(d7) | DHSph subclass |

| | | | |
|---|---|---|---|
| Type of quantification | Internal standard amount | Batch correction | No |
| Response correction | No | Further quantification remarks | - |
| Type I isotope correction | No | | |

#### 122) Sphingomyelin (SM)[M+CH3COO]- / Lipid identification

| | | | |
|---|---|---|---|
| Lipid class | Sphingomyelin (SM) | Background check at MS2 | No |
| Derivatization | - | Did you presume assumptions for identification? | No |
| MS Level for identification | MS1, MS2 | Check isomer overlap | No |
| Identification level | Molecular species level | RT verified by standard | Yes |
| Polarity mode | Negative | Separation of isobaric/isomeric interferece confirmed | Yes |
| Type of negative (precursor)ion | [M+CH3COO]- | Model for separation prediction | Yes |
| Fragments for identification | | Additional dimension/techniques | - |

| Fragment name |
|---|
| Neutral loss of methyl moiety |
| Characteristic fragment (C4H11NO4P-) |
| Neutral loss of methyl moiety and acyl |

| | | | |
|---|---|---|---|
| Isotope correction at MS1 | No | Lipid Identification Software | MS-DIAL |
| Isotope correction at MS2 | No | Data manipulation | Smoothing, Centroiding |
| MS1 verified by standard | Yes | Nomenclature for intact lipid molecule | Yes |
| MS2 verified by standard | Yes | Nomenclature for fragment ions | No |
| Background check at MS1 | Yes | Further identification remarks | - |

#### 122) Sphingomyelin (SM)[M+CH3COO]- / Lipid quantification

| | | | |
|---|---|---|---|
| Quantitative | Yes | Limit of quantification | No |
| MS Level for quantification | MS1 | Normalization to reference | No |
| Internal lipid standard(s) MS1 | | Lipid Quantification Software | MS-DIAL |

| Internal standard | Endogenous subclass |
|---|---|
| SM 18:1;2O/18:1(d9) | SM subclass |

| | | | |
|---|---|---|---|
| Type of quantification | Internal standard amount | Batch correction | No |
| Response correction | No | Further quantification remarks | - |
| Type I isotope correction | No | | |

#### 123) SPB[M+H]+ / Lipid identification

| | | | |
|---|---|---|---|
| Lipid class | SPB | Background check at MS2 | No |
| Derivatization | - | Did you presume assumptions for identification? | No |
| MS Level for identification | MS1, MS2 | Check isomer overlap | No |
| Identification level | Molecular species level | RT verified by standard | Yes |
| Polarity mode | Positive | Separation of isobaric/isomeric interferece confirmed | Yes |
| Type of positive (precursor)ion | [M+H]+ | Model for separation prediction | Yes |
| Fragments for identification | | Additional dimension/techniques | - |

| Fragment name |
|---|
| Neutral loss of H2O |
| Neutral loss of 2H2O |
| Neutral loss of CH4O2 |

| | | | |
|---|---|---|---|
| Isotope correction at MS1 | No | Lipid Identification Software | MS-DIAL |
| Isotope correction at MS2 | No | Data manipulation | Smoothing, Centroiding |
| MS1 verified by standard | No | Nomenclature for intact lipid molecule | Yes |
| MS2 verified by standard | No | Nomenclature for fragment ions | No |
| Background check at MS1 | Yes | Further identification remarks | - |

#### 123) SPB[M+H]+ / Lipid quantification

| | | | |
|---|---|---|---|
| Quantitative | Yes | Limit of quantification | No |
| MS Level for quantification | MS1 | Normalization to reference | No |
| Internal lipid standard(s) MS1 | | Lipid Quantification Software | MS-DIAL |

| Internal standard | Endogenous subclass |
|---|---|
| Cer 18:1;2O/15:0(d7) | Sph subclass |

| | | | |
|---|---|---|---|
| Type of quantification | Internal standard amount | Batch correction | No |
| Response correction | No | Further quantification remarks | - |
| Type I isotope correction | No | | |

#### 124) Sterol sulfate (SSulfate)[M-H]- / Lipid identification

| | | | |
|---|---|---|---|
| Lipid class | Sterol sulfate (SSulfate) | Background check at MS2 | No |
| Derivatization | - | Did you presume assumptions for identification? | No |
| MS Level for identification | MS1, MS2 | Check isomer overlap | No |
| Identification level | Molecular species level | RT verified by standard | Yes |
| Polarity mode | Negative | Separation of isobaric/isomeric interferece confirmed | Yes |
| Type of negative (precursor)ion | [M-H]- | Model for separation prediction | Yes |
| Fragments for identification | | Additional dimension/techniques | - |

| Fragment name |
|---|
| Cholesterol sulfate |
| Sulfate |

| | | | |
|---|---|---|---|
| Isotope correction at MS1 | No | Lipid Identification Software | MS-DIAL |
| Isotope correction at MS2 | No | Data manipulation | Smoothing, Centroiding |
| MS1 verified by standard | No | Nomenclature for intact lipid molecule | Yes |
| MS2 verified by standard | No | Nomenclature for fragment ions | No |
| Background check at MS1 | Yes | Further identification remarks | - |

#### 124) Sterol sulfate (SSulfate)[M-H]- / Lipid quantification

| | | | |
|---|---|---|---|
| Quantitative | Yes | Limit of quantification | No |
| MS Level for quantification | MS1 | Normalization to reference | No |
| Internal lipid standard(s) MS1 | | Lipid Quantification Software | MS-DIAL |

| Internal standard | Endogenous subclass |
|---|---|
| LPC 18:1(d7) | SSulfate subclass |

| | | | |
|---|---|---|---|
| Type of quantification | Internal standard amount | Batch correction | No |
| Response correction | No | Further quantification remarks | - |
| Type I isotope correction | No | | |

#### 125) Stigmasterol ester (STSE)[M+NH4]+ / Lipid identification

| | | | |
|---|---|---|---|
| Lipid class | Stigmasterol ester (STSE) | Background check at MS2 | No |
| Derivatization | - | Did you presume assumptions for identification? | No |
| MS Level for identification | MS1, MS2 | Check isomer overlap | No |
| Identification level | Molecular species level | RT verified by standard | Yes |
| Polarity mode | Positive | Separation of isobaric/isomeric interferece confirmed | Yes |
| Type of positive (precursor)ion | [M+NH4]+ | Model for separation prediction | Yes |
| Fragments for identification | | Additional dimension/techniques | - |

| Fragment name |
|---|
| Neutral loss of fatty acid |

| | | | |
|---|---|---|---|
| Isotope correction at MS1 | No | Lipid Identification Software | MS-DIAL |
| Isotope correction at MS2 | No | Data manipulation | Smoothing, Centroiding |
| MS1 verified by standard | No | Nomenclature for intact lipid molecule | Yes |
| MS2 verified by standard | No | Nomenclature for fragment ions | No |
| Background check at MS1 | Yes | Further identification remarks | - |

#### 125) Stigmasterol ester (STSE)[M+NH4]+ / Lipid quantification

| | | | |
|---|---|---|---|
| Quantitative | Yes | Limit of quantification | No |
| MS Level for quantification | MS1 | Normalization to reference | No |
| Internal lipid standard(s) MS1 | | Lipid Quantification Software | MS-DIAL |

| Internal standard | Endogenous subclass |
|---|---|
| CE 18:1(d7) | STSE subclass |

| | | | |
|---|---|---|---|
| Type of quantification | Internal standard amount | Batch correction | No |
| Response correction | No | Further quantification remarks | - |
| Type I isotope correction | No | | |

#### 126) Stigmasterol hexoside (SHex)[M+CH3COO]- / Lipid identification

| | | | |
|---|---|---|---|
| Lipid class | Stigmasterol hexoside (SHex) | Background check at MS2 | No |
| Derivatization | - | Did you presume assumptions for identification? | No |
| MS Level for identification | MS1, MS2 | Check isomer overlap | No |
| Identification level | Molecular species level | RT verified by standard | Yes |
| Polarity mode | Negative | Separation of isobaric/isomeric interferece confirmed | Yes |
| Type of negative (precursor)ion | [M+CH3COO]- | Model for separation prediction | Yes |
| Fragments for identification | | Additional dimension/techniques | - |

| Fragment name |
|---|
| Hexose |

| | | | |
|---|---|---|---|
| Isotope correction at MS1 | No | Lipid Identification Software | MS-DIAL |
| Isotope correction at MS2 | No | Data manipulation | Smoothing, Centroiding |
| MS1 verified by standard | No | Nomenclature for intact lipid molecule | Yes |
| MS2 verified by standard | No | Nomenclature for fragment ions | No |
| Background check at MS1 | Yes | Further identification remarks | - |

#### 126) Stigmasterol hexoside (SHex)[M+CH3COO]- / Lipid quantification

| | | | |
|---|---|---|---|
| Quantitative | Yes | Limit of quantification | No |
| MS Level for quantification | MS1 | Normalization to reference | No |
| Internal lipid standard(s) MS1 | | Lipid Quantification Software | MS-DIAL |

| Internal standard | Endogenous subclass |
|---|---|
| LPC 18:1(d7) | SHex subclass |

| | | | |
|---|---|---|---|
| Type of quantification | Internal standard amount | Batch correction | No |
| Response correction | No | Further quantification remarks | - |
| Type I isotope correction | No | | |

#### 127) SHexCer[M+H]+ / Lipid identification

| | | | |
|---|---|---|---|
| Lipid class | SHexCer | Background check at MS2 | No |
| Derivatization | - | Did you presume assumptions for identification? | No |
| MS Level for identification | MS1, MS2 | Check isomer overlap | No |
| Identification level | Molecular species level | RT verified by standard | Yes |
| Polarity mode | Positive | Separation of isobaric/isomeric interferece confirmed | Yes |
| Type of positive (precursor)ion | [M+H]+ | Model for separation prediction | Yes |
| Fragments for identification | | Additional dimension/techniques | - |

| Fragment name |
|---|
| Neutral loss of sulfate |
| Neutral loss of sulfate and hexose |
| Sphingosine -H2O fragment |
| Sphingosine -2H2O fragment |
| Sphingosine -CH4O2 fragment |

| | | | |
|---|---|---|---|
| Isotope correction at MS1 | No | Lipid Identification Software | MS-DIAL |
| Isotope correction at MS2 | No | Data manipulation | Smoothing, Centroiding |
| MS1 verified by standard | No | Nomenclature for intact lipid molecule | Yes |
| MS2 verified by standard | No | Nomenclature for fragment ions | No |
| Background check at MS1 | Yes | Further identification remarks | - |

#### 127) SHexCer[M+H]+ / Lipid quantification

| | | | |
|---|---|---|---|
| Quantitative | Yes | Limit of quantification | No |
| MS Level for quantification | MS1 | Normalization to reference | No |
| Internal lipid standard(s) MS1 | | Lipid Quantification Software | MS-DIAL |

| Internal standard | Endogenous subclass |
|---|---|
| Cer 18:1;2O/15:0(d7) | SHexCer subclass |

| | | | |
|---|---|---|---|
| Type of quantification | Internal standard amount | Batch correction | No |
| Response correction | No | Further quantification remarks | - |
| Type I isotope correction | No | | |

#### 128) SQDG[M+NH4]+ / Lipid identification

| | | | |
|---|---|---|---|
| Lipid class | SQDG | Background check at MS2 | No |
| Derivatization | - | Did you presume assumptions for identification? | No |
| MS Level for identification | MS1, MS2 | Check isomer overlap | No |
| Identification level | Molecular species level | RT verified by standard | Yes |
| Polarity mode | Positive | Separation of isobaric/isomeric interferece confirmed | Yes |
| Type of positive (precursor)ion | [M+NH4]+ | Model for separation prediction | Yes |
| Fragments for identification | | Additional dimension/techniques | - |

| Fragment name |
|---|
| Neutral loss of C6H10O7S |
| Dehydro-monoacyl glycerols |

| | | | |
|---|---|---|---|
| Isotope correction at MS1 | No | Lipid Identification Software | MS-DIAL |
| Isotope correction at MS2 | No | Data manipulation | Smoothing, Centroiding |
| MS1 verified by standard | No | Nomenclature for intact lipid molecule | Yes |
| MS2 verified by standard | No | Nomenclature for fragment ions | No |
| Background check at MS1 | Yes | Further identification remarks | - |

#### 128) SQDG[M+NH4]+ / Lipid quantification

| | | | |
|---|---|---|---|
| Quantitative | Yes | Limit of quantification | No |
| MS Level for quantification | MS1 | Normalization to reference | No |
| Internal lipid standard(s) MS1 | | Lipid Quantification Software | MS-DIAL |

| Internal standard | Endogenous subclass |
|---|---|
| LPC 18:1(d7) | SQDG subclass |

| | | | |
|---|---|---|---|
| Type of quantification | Internal standard amount | Batch correction | No |
| Response correction | No | Further quantification remarks | - |
| Type I isotope correction | No | | |

#### 129) TG[M+NH4]+ / Lipid identification

| | | | |
|---|---|---|---|
| Lipid class | TG | Background check at MS2 | No |
| Derivatization | - | Did you presume assumptions for identification? | No |
| MS Level for identification | MS1, MS2 | Check isomer overlap | No |
| Identification level | Molecular species level | RT verified by standard | Yes |
| Polarity mode | Positive | Separation of isobaric/isomeric interferece confirmed | Yes |
| Type of positive (precursor)ion | [M+NH4]+ | Model for separation prediction | Yes |
| Fragments for identification | | Additional dimension/techniques | - |

| Fragment name |
|---|
| Neutral loss of acyl and H2O |

| | | | |
|---|---|---|---|
| Isotope correction at MS1 | No | Lipid Identification Software | MS-DIAL |
| Isotope correction at MS2 | No | Data manipulation | Smoothing, Centroiding |
| MS1 verified by standard | Yes | Nomenclature for intact lipid molecule | Yes |
| MS2 verified by standard | Yes | Nomenclature for fragment ions | No |
| Background check at MS1 | Yes | Further identification remarks | - |

#### 129) TG[M+NH4]+ / Lipid quantification

| | | | |
|---|---|---|---|
| Quantitative | Yes | Limit of quantification | No |
| MS Level for quantification | MS1 | Normalization to reference | No |
| Internal lipid standard(s) MS1 | | Lipid Quantification Software | MS-DIAL |

| Internal standard | Endogenous subclass |
|---|---|
| TG 15:0_18:1(d7)_15:0 | TG subclass |

| | | | |
|---|---|---|---|
| Type of quantification | Internal standard amount | Batch correction | No |
| Response correction | No | Further quantification remarks | - |
| Type I isotope correction | No | | |

#### 130) Triacylglycerol estolides (TG_EST)[M+NH4]+ / Lipid identification

| | | | |
|---|---|---|---|
| Lipid class | Triacylglycerol estolides (TG_EST) | Background check at MS2 | No |
| Derivatization | - | Did you presume assumptions for identification? | No |
| MS Level for identification | MS1, MS2 | Check isomer overlap | No |
| Identification level | Molecular species level | RT verified by standard | Yes |
| Polarity mode | Positive | Separation of isobaric/isomeric interferece confirmed | Yes |
| Type of positive (precursor)ion | [M+NH4]+ | Model for separation prediction | Yes |
| Fragments for identification | | Additional dimension/techniques | - |

| Fragment name |
|---|
| Neutral loss of acyl and H2O |
| Neutral loss of oxidized acyl and H2O |
| Neutral loss of FAHFA |

| | | | |
|---|---|---|---|
| Isotope correction at MS1 | No | Lipid Identification Software | MS-DIAL |
| Isotope correction at MS2 | No | Data manipulation | Smoothing, Centroiding |
| MS1 verified by standard | No | Nomenclature for intact lipid molecule | Yes |
| MS2 verified by standard | No | Nomenclature for fragment ions | No |
| Background check at MS1 | Yes | Further identification remarks | - |

#### 130) Triacylglycerol estolides (TG_EST)[M+NH4]+ / Lipid quantification

| | | | |
|---|---|---|---|
| Quantitative | Yes | Limit of quantification | No |
| MS Level for quantification | MS1 | Normalization to reference | No |
| Internal lipid standard(s) MS1 | | Lipid Quantification Software | MS-DIAL |

| Internal standard | Endogenous subclass |
|---|---|
| TG 15:0_18:1(d7)_15:0 | TG_EST subclass |

| | | | |
|---|---|---|---|
| Type of quantification | Internal standard amount | Batch correction | No |
| Response correction | No | Further quantification remarks | - |
| Type I isotope correction | No | | |

#### 131) Hex3Cer[M+H]+ / Lipid identification

| | | | |
|---|---|---|---|
| Lipid class | Hex3Cer | Background check at MS2 | No |
| Derivatization | - | Did you presume assumptions for identification? | No |
| MS Level for identification | MS1, MS2 | Check isomer overlap | No |
| Identification level | Molecular species level | RT verified by standard | Yes |
| Polarity mode | Positive | Separation of isobaric/isomeric interferece confirmed | Yes |
| Type of positive (precursor)ion | [M+H]+ | Model for separation prediction | Yes |
| Fragments for identification | | Additional dimension/techniques | - |

| Fragment name |
|---|
| Neutral loss of hexose |
| Neutral loss of 2hexose |
| Neutral loss of 3hexose |
| Neutral loss of 3hexose and H2O |
| Sphingosine -H2O fragment |
| Sphingosine -2H2O fragment |
| Sphingosine -CH4O2 fragment |

| | | | |
|---|---|---|---|
| Isotope correction at MS1 | No | Lipid Identification Software | MS-DIAL |
| Isotope correction at MS2 | No | Data manipulation | Smoothing, Centroiding |
| MS1 verified by standard | No | Nomenclature for intact lipid molecule | Yes |
| MS2 verified by standard | No | Nomenclature for fragment ions | No |
| Background check at MS1 | Yes | Further identification remarks | - |

#### 131) Hex3Cer[M+H]+ / Lipid quantification

| | | | |
|---|---|---|---|
| Quantitative | Yes | Limit of quantification | No |
| MS Level for quantification | MS1 | Normalization to reference | No |
| Internal lipid standard(s) MS1 | | Lipid Quantification Software | MS-DIAL |

| Internal standard | Endogenous subclass |
|---|---|
| Cer 18:1;2O/15:0(d7) | Hex3Cer subclass |

| | | | |
|---|---|---|---|
| Type of quantification | Internal standard amount | Batch correction | No |
| Response correction | No | Further quantification remarks | - |
| Type I isotope correction | No | | |

#### 132) Vitamin A fatty acid ester (VAE)[M+Na]+ / Lipid identification

| | | | |
|---|---|---|---|
| Lipid class | Vitamin A fatty acid ester (VAE) | Background check at MS2 | No |
| Derivatization | - | Did you presume assumptions for identification? | No |
| MS Level for identification | MS1, MS2 | Check isomer overlap | No |
| Identification level | Molecular species level | RT verified by standard | Yes |
| Polarity mode | Positive | Separation of isobaric/isomeric interferece confirmed | Yes |
| Type of positive (precursor)ion | [M+Na]+ | Model for separation prediction | Yes |
| Fragments for identification | | Additional dimension/techniques | - |

| Fragment name |
|---|
| Characteristic fragment (C20H29+) |
| Characteristic fragment (C9H11+) |

| | | | |
|---|---|---|---|
| Isotope correction at MS1 | No | Lipid Identification Software | MS-DIAL |
| Isotope correction at MS2 | No | Data manipulation | Smoothing, Centroiding |
| MS1 verified by standard | No | Nomenclature for intact lipid molecule | Yes |
| MS2 verified by standard | No | Nomenclature for fragment ions | No |
| Background check at MS1 | Yes | Further identification remarks | - |

#### 132) Vitamin A fatty acid ester (VAE)[M+Na]+ / Lipid quantification

| | | | |
|---|---|---|---|
| Quantitative | Yes | Limit of quantification | No |
| MS Level for quantification | MS1 | Normalization to reference | No |
| Internal lipid standard(s) MS1 | | Lipid Quantification Software | MS-DIAL |

| Internal standard | Endogenous subclass |
|---|---|
| LPC 18:1(d7) | VAE subclass |

| | | | |
|---|---|---|---|
| Type of quantification | Internal standard amount | Batch correction | No |
| Response correction | No | Further quantification remarks | - |
| Type I isotope correction | No | | |

#### 133) Vitamin D[M+H]+ / Lipid identification

| | | | |
|---|---|---|---|
| Lipid class | Vitamin D | Background check at MS2 | No |
| Derivatization | - | Did you presume assumptions for identification? | No |
| MS Level for identification | MS1, MS2 | Check isomer overlap | No |
| Identification level | Species level | RT verified by standard | Yes |
| Polarity mode | Positive | Separation of isobaric/isomeric interferece confirmed | Yes |
| Type of positive (precursor)ion | [M+H]+ | Model for separation prediction | Yes |
| Fragments for identification | | Additional dimension/techniques | - |

| Fragment name |
|---|
| Neutral loss of H2O |
| Neutral loss of 2H2O |

| | | | |
|---|---|---|---|
| Isotope correction at MS1 | No | Lipid Identification Software | MS-DIAL |
| Isotope correction at MS2 | No | Data manipulation | Smoothing, Centroiding |
| MS1 verified by standard | No | Nomenclature for intact lipid molecule | Yes |
| MS2 verified by standard | No | Nomenclature for fragment ions | No |
| Background check at MS1 | Yes | Further identification remarks | - |

#### 133) Vitamin D[M+H]+ / Lipid quantification

| | | | |
|---|---|---|---|
| Quantitative | Yes | Limit of quantification | No |
| MS Level for quantification | MS1 | Normalization to reference | No |
| Internal lipid standard(s) MS1 | | Lipid Quantification Software | MS-DIAL |

| Internal standard | Endogenous subclass |
|---|---|
| LPC 18:1(d7) | Vitamin D subclass |

| | | | |
|---|---|---|---|
| Type of quantification | Internal standard amount | Batch correction | No |
| Response correction | No | Further quantification remarks | - |
| Type I isotope correction | No | | |

#### 134) Vitamin E[M+CH3COO]- / Lipid identification

| | | | |
|---|---|---|---|
| Lipid class | Vitamin E | Background check at MS2 | No |
| Derivatization | - | Did you presume assumptions for identification? | No |
| MS Level for identification | MS1, MS2 | Check isomer overlap | No |
| Identification level | Species level | RT verified by standard | Yes |
| Polarity mode | Negative | Separation of isobaric/isomeric interferece confirmed | Yes |
| Type of negative (precursor)ion | [M+CH3COO]- | Model for separation prediction | Yes |
| Fragments for identification | | Additional dimension/techniques | - |

| Fragment name |
|---|
| Characteristic fragment (C10H11O2-) |

| | | | |
|---|---|---|---|
| Isotope correction at MS1 | No | Lipid Identification Software | MS-DIAL |
| Isotope correction at MS2 | No | Data manipulation | Smoothing, Centroiding |
| MS1 verified by standard | No | Nomenclature for intact lipid molecule | Yes |
| MS2 verified by standard | No | Nomenclature for fragment ions | No |
| Background check at MS1 | Yes | Further identification remarks | - |

#### 134) Vitamin E[M+CH3COO]- / Lipid quantification

| | | | |
|---|---|---|---|
| Quantitative | Yes | Limit of quantification | No |
| MS Level for quantification | MS1 | Normalization to reference | No |
| Internal lipid standard(s) MS1 | | Lipid Quantification Software | MS-DIAL |

| Internal standard | Endogenous subclass |
|---|---|
| LPC 18:1(d7) | Vitamin E subclass |

| | | | |
|---|---|---|---|
| Type of quantification | Internal standard amount | Batch correction | No |
| Response correction | No | Further quantification remarks | - |
| Type I isotope correction | No | | |

#### 135) PE P[M+H]+ / Lipid identification

| | | | |
|---|---|---|---|
| Lipid class | PE P | Background check at MS2 | No |
| Derivatization | - | Did you presume assumptions for identification? | No |
| MS Level for identification | MS1, MS2 | Check isomer overlap | No |
| Identification level | Molecular species level | RT verified by standard | Yes |
| Polarity mode | Positive | Separation of isobaric/isomeric interferece confirmed | Yes |
| Type of positive (precursor)ion | [M+H]+ | Model for separation prediction | Yes |
| Fragments for identification | | Additional dimension/techniques | - |

| Fragment name |
|---|
| Neutral loss of C2H8NO4P |
| Alkyl ether +C2H8NO3P fragment |
| Dehydro-monoacyl glycerols |

| | | | |
|---|---|---|---|
| Isotope correction at MS1 | No | Lipid Identification Software | MS-DIAL |
| Isotope correction at MS2 | No | Data manipulation | Smoothing, Centroiding |
| MS1 verified by standard | No | Nomenclature for intact lipid molecule | Yes |
| MS2 verified by standard | No | Nomenclature for fragment ions | No |
| Background check at MS1 | Yes | Further identification remarks | - |

#### 135) PE P[M+H]+ / Lipid quantification

| | | | |
|---|---|---|---|
| Quantitative | Yes | Limit of quantification | No |
| MS Level for quantification | MS1 | Normalization to reference | No |
| Internal lipid standard(s) MS1 | | Lipid Quantification Software | MS-DIAL |

| Internal standard | Endogenous subclass |
|---|---|
| PE 15:0_18:1(d7) | EtherPE subclass |

| | | | |
|---|---|---|---|
| Type of quantification | Internal standard amount | Batch correction | No |
| Response correction | No | Further quantification remarks | - |
| Type I isotope correction | No | | |
