## Supplementary Note 4 for "MS-DIAL 5 multimodal mass spectrometry data mining unveils lipidome complexities"

### Separation Workflow

Created by https://lipidomicstandards.org, version v2.3.2

#### Overall study design

| | | | |
|---|---|---|---|
| Title of the study | Lysophospholipid profiling by using trimethylsilyl-diazomethane for HeLa cells | | |
| Document creation date | 02/07/2024 | Corresponding Email | |
| Principle investigator | Hiroshi Tsugawa | Is the workflow targeted or untargeted? | Targeted |
| Institution | Tokyo University of Agriculture and Technology | Clinical | No |

#### Lipid extraction

| | | | |
|---|---|---|---|
| Extraction method | 1-phase system | 1-phase system | Methanol |
| pH adjustment | None | Were internal standards added prior extraction? | No |

#### Analytical platform

| | | | |
|---|---|---|---|
| Which solvents were used | (A) acetonitrile (ACN):MeOH:H2O (1:1:3, v/v/v) and (B) ACN:IPA (1:9, v/v). Both the solvents contained 10 nM ethylenediaminetetraacetic acid and 5 mM ammonium acetate. | Mass resolution for detected ion at MS1 | High resolution |
| Number of separation dimensions | One dimension | Resolution at m/z 200 at MS1 | 25148 |
| Separation type 1 | LC | Mass accuracy in ppm at MS1 | 0.3 |
| Separation mode 1 (liquid) | RP | Mass window for precursor ion isolation (in Da total isolation window) | 1 |
| Detector | Mass spectrometer | Mass resolution for detected ion at MS2 | High resolution |
| MS type | QTOF | Resolution at m/z 200 at MS2 | 26373 |
| MS vendor | SCIEX | Mass accuracy in ppm at MS2 | 0.67 |
| Ion source | ESI | Was/Were additional dimension/techniques used | No |
| MS Level | MS1, MS2 | | |

#### Quality control

| | | | |
|---|---|---|---|
| Blanks | Yes | Quality control | No |
| Type of Blanks | Extraction blank, Injection blank | | |

#### Method qualification and validation

| | | | |
|---|---|---|---|
| Method validation | Yes | Precison | Yes |
| Lipid recovery | Yes | Accuracy | Yes |
| Dynamic quantification range | No | Guidelines followed | None |
| Limit of quantitation (LOQ)/Limit of detection (LOD) | No | | |

#### Reporting

| | | | |
|---|---|---|---|
| Are reported raw data uploaded into repository? | Yes | Summary data | Quantification and identification data |
| Link to repository / ID to entry | http://prime.psc.riken.jp/menta.cgi/prime/drop_index | Raw data upload | Yes |
| Are metadata available? | Yes | Additional comments | The resolution and accuracy (ppm) for MS1 are those of m/z 132.9049 which were determined by the mass calibration in SCIEX OS. In addition, the resolution and accuracy for MS2 are those of m/z 185.1284, which were also determined by the mass calibration in SCIEX OS. |

### Sample Descriptions

#### HeLa cell with the supplementation of VLC-PUFA (FA 32:6) / Human / Cells

| | | | |
|---|---|---|---|
| Provided information | Time to freeze (min), Storage time (month), Freeze-thaw cycles | Storage time (month) | 1 |
| Temperature handling original sample | 4-8 °C | Freeze-thaw cycles | 0 |
| Instant sample preparation | No | Additives | None |
| Time to freeze (min) | 10 | Were samples stored under inert gas? | No |
| Snap freezing in liquid N2 | Yes | Additional preservation methods | No |
| Storage temperature | -80 °C | Biobank samples | No |

### Lipid Class Descriptions

#### 1) LPA[M+H]+ / Lipid identification

| | | | |
|---|---|---|---|
| Lipid class | LPA | Background check at MS2 | Yes |
| Derivatization | Trimethylsilyl (TMS)-diazomethane | Did you presume assumptions for identification? | No |
| MS Level for identification | MS1, MS2 | Check isomer overlap | Yes |
| Identification level | Molecular species level | RT verified by standard | Yes |
| Polarity mode | Positive | Separation of isobaric/isomeric interferece confirmed | Yes |
| Type of positive (precursor)ion | [M+H]+ | Model for separation prediction | No |
| Fragments for identification | | Additional dimension/techniques | - |

| Fragment name |
|---|
| Product ion of m/z 127.01547 (theoretical value) meaning C2H8O4P (Bismethyl PO4) |
| Neutral loss of 126.0082 (theoretical value) meaning the loss of bismethyl PO4 |

| | | | |
|---|---|---|---|
| Isotope correction at MS1 | No | Lipid Identification Software | MS-DIAL |
| Isotope correction at MS2 | No | Data manipulation | Smoothing, Centroiding |
| MS1 verified by standard | Yes | Nomenclature for intact lipid molecule | Yes |
| MS2 verified by standard | Yes | Nomenclature for fragment ions | N/A |
| Background check at MS1 | Yes | Further identification remarks | - |

#### 1) LPA[M+H]+ / Lipid quantification

| | | | |
|---|---|---|---|
| Quantitative | Yes | Limit of quantification | No |
| MS Level for quantification | MS1 | Normalization to reference | No |
| Internal lipid standard(s) MS1 | | Lipid Quantification Software | MS-DIAL |

| Internal standard | Endogenous subclass |
|---|---|
| LPA 17:1 | LPA |

| | | | |
|---|---|---|---|
| Type of quantification | Internal standard amount | Batch correction | No |
| Response correction | No | Further quantification remarks | In the paper, the original peak height are described. |
| Type I isotope correction | No | | |
