## Supplementary material for "MS-DIAL 5 multimodal mass spectrometry data mining unveils lipidome complexities": Source data: readme.txt

#annofile contains the annotation results in each file. The "no MS/MS" information is retrived from this file.
#pairfile contains the definition of annotation levels.
#peaknamefile contains the annotation results in each file. The annotation name is compared to the list of pairfile.
#resultexport contains the result, which is made by integrating annofile, pairfile, and peaknamefile, to create the figure.
#productiontables folder contains the ion abundances of product ions of each lipid structure in each file. The file name means the alignment spot ID. For example, AlignmentID000291.txt contains the spectra of each file aligned to the alignedspot.spotid of 291 (CE 18:1) in peaknamefile.txt. 
